# Neither global nor consistent: a technical comment on the tree diversity-productivity analysis of Liang et al. (2016)

**DOI:** 10.1101/524363

**Authors:** Carsten F. Dormann, Helge Schneider, Jonas Gorges

## Abstract

The publication of Liang *et al.* (2016, Science) seems to demonstrate very clearly that increasing tree species richness substantially increases forest productivity. To combine data from very different ecoregions, the authors constructed a relative measure of tree species richness. This relative richness however confounds plot-level tree species richness and the polar-tropical gradient of tree species richness. We re-analysed their orginal data, computing a regional measure of tree species richness and addressing several other issues in their analysis. We find that there is virtually no effect of relative tree species richness on productivity when computing species richness at the local scale. Also, different ecoregions have very different relationships between tree species richness and productivity. Thus, neither the “global” consistency nor the actual effect can be confirmed.

---

Observational studies of the species-richness-ecosystem functioning relationship are correlational and not causal. Observing a consistent correlative effect of tree species richness on forest productivity (TSP-P) across a wide range of biomes, however, would make this relationship a reliable principle on which to build recommendations for forestry and forest conservation. The publication of Liang *et al.* (2016) report on such a consistent, global TSP-P-relationship, based on forest inventory data from over 600,000 plots in 12 different ecoregions.

There are several logical and technical flaws in their analysis; correcting them also removes the reported global relationship and thus its role as principle for forest management. We identified the following problems with the original analysis: (1) The authors computed “relative tree species richness” as proportion of the maximum value. Thereby it represents a gradient from boreal to tropical plots, rather than in *local* species richness. When instead computing species richness relative to the maximum value in the region the effect of species richness on productivity is dramatically reduced. (2) Plots are overwhelmingly from temperate forest; indeed only some 2500 plots are from the tropics (equivalent to 0.4%), despite these forests representing around 30% of the world’s forest. Stratifying the plots accordingly weakens the TSR-P-relationship. (3) In the spatial regression model, distances between plots were computed without taking the spherical nature of earth into account. This had little effect on the slope estimate of the TSR-P-relationship. (4) The computational burden of the spatial model required subsampling the data to 500 data points. The authors did not correctly compute confidence intervals for this approach, wrongly interpreting subsampling as bootstrapping and additionally incorrectly computing bootstrap standard errors. A correct subsampling-based estimation led to approximate trippling of the reported confidence interval. (5) As noted earlier (Schulze *et al.,* 2018), some 4% of the plots had productivity values (far) beyond what is biologically plausible (Stape *et al.,* 2010). The likely reason is that small plots with large inventory errors in the productivity may lead to erratically high values. Not taking this into account in the analysis, e.g. by down-weighting plots with productivities above 30 m^2^ha^−1^y^−1^ at least indicates an unreflected use of data. It also leads to an overestimation of the absolute productivity. Whether that is the main reason for productivity increases dramatically higher than any reported from experimental setups (Zhang *et al.,* 2012a; Vilà *et al.,* 2013; Huang *et al.,* 2018; Jactel *et al.,* 2018; Ammer, 2019) is unclear.

Combining corrections for these points led to a new global analysis, with a negligible effect of tree species richness on a site’s volumetric productivity (Fig. 1 left; see supplementary material for the full analysis including R-code). Furthermore, fitting the model to the 11 ecoregions with sufficient data separately shows that the global model is not consistent and provides a poor description of the ecoregions’ specific TSR-P-relationships (Fig. 1 right). Also, there is no obvious benefit of scaling species richness, as the curves are more difficult to interpret when displayed as percentage of some maximal value (Fig. 2). Indeed, using a relative scale makes the species richness-effect (the *θ*-estimate) harder to compare across biomes and insignificant in several cases.

**Figure 1:**
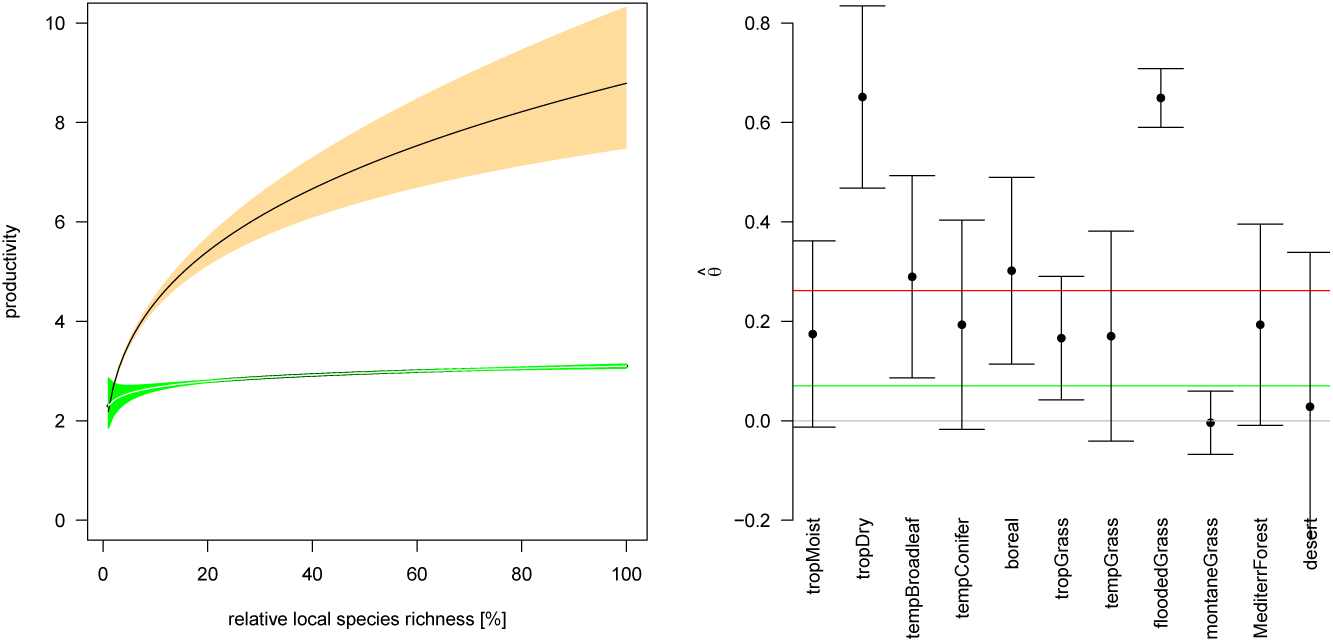
Comparison of reproduced analysis of Liang *et al.* (2016) (black) and corrected analysis of the same data (green), left, and re-analysis of the estimated species richness effect 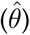 for each ecoregion separately, right. Red and green lines, right, are the original and the global re-analysis, respectively.

**Figure 2:**
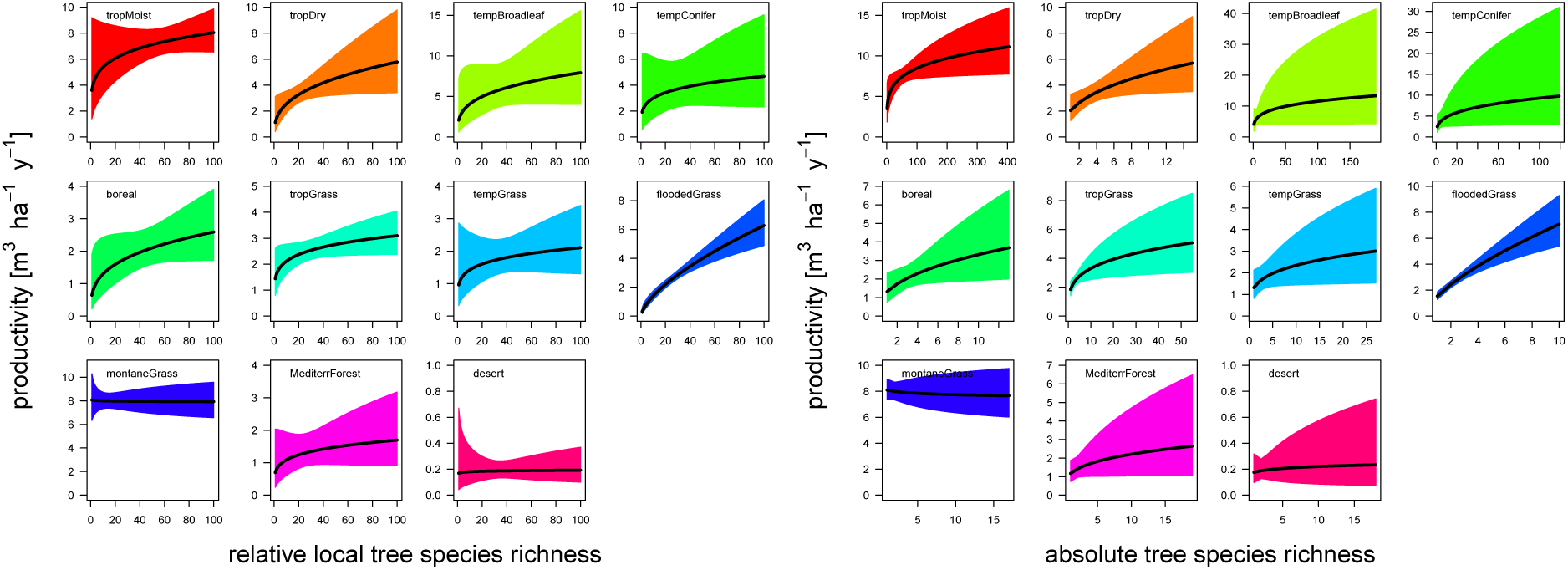
Biome-specific observed tree-species richness-productivity relationships (and 95% confidence interval). (Left) Species richness expressed as percentage of regional maximum. (Right) Productivity as changing with absolute species richness. Note that the shape of the curves can change when re-scaling species richness, as the values enter the analysis log-transformed.

In summary, the analysis presented by Liang *et al.* (2016) is flawed in several respects, leading to a spuriously strong effect of tree species richness on forest productivity at the global level. While a re-analysis correlatively confirms a positive TSR-P-relationship for most ecoregion, these effects vary substantially in their strength and should be examined for each system separately. Overall, the original publication has thus added very little to our understanding, as both positive and biome-specific effects of species richness were already known before their publication (see, e.g., Whittaker & Heegaard, 2003; Vilà *et al.,* 2007; Paquette & Messier, 2010; Whittaker, 2010; Zhang *et al.,* 2012b), and the global consistency could not be upheld.

## Appendix: Reproducible analysis with comments and conclu-sions for each step

### A Introduction

The positive effect of species richness on various measure of ecosystem functioning is generally undisputed (Cardinale *et al.,* 2012), even though this effect levels off fairly rapidly as the number of species beyond “few” (Cardinale *et al.,* 2006; Isbell *et al.,* 2011). Such fundamental understanding does not necessarily translate into applicable knowledge, and hence the publication of Liang *et al.* (2016) is a landmark step in converting academic biodiversity-ecosystem functioning into on-the-ground forest management. They claim to describe a globally applicable relationship between tree species richness and forest productivity, which, if correct, could guide forest management around the world without specific understanding of local or regional specifics.

On re-analysing the excellent data base compiled by Liang *et al.* (2016), we were originally curious to investigate an unanswered question, namely whether the global tree-richness-productivity-relationship is actually anywhere near as good as one built, on the same data, for a specific biome. Furthermore, we were also concerned with describing a partial dependence plot for the effect of tree species richness, rather than the conditional plot presented by Liang *et al.* (2016); the difference being that in their plot, the environmental predictors used in the model (such as basal area, temperature, precipitation and elevation) must be known, while in a partial dependency plot the *marginal* effect of tree species richness, independent of the actual local setting, would be depicted.

During the analysis, we realised that these initial questions were of secondary importance, as we discovered conceptual and statistical issues that cast substantial doubt on the results presented by Liang *et al.* (2016). Since the first author initially did not provide his code, we had to rely on the method description and guessing to reproduce the analysis.^1^ From the reconstruction we modified the original model to correct for some mistakes, as detailed below, to derive our new version of the global relationship. In the next step, we predicted from the global model to the average conditions of each biome, and compared this prediction with those of a biome-specific model from the same data base.

In the following, we present first the original analysis, then the modifications we introduce. We close with a new global and a biome-specific analysis.

### B Reproducing the original tree richness-productivity relationship

#### B.1 Data preparation

The data are available under https://figshare.com/articles/GFB1_data_figshare_xlsx/4286552 for all countries except New Zealand, whose data are available from https://datastore.landcareresearch.co.nz/dataset/new-zealand-forest-plot-data-in-global-forest-biodiversity-dataset, both as Excel files. Upon downloading and merging the files, here is a summary of its content (in the following, code used for checking is outcommented):

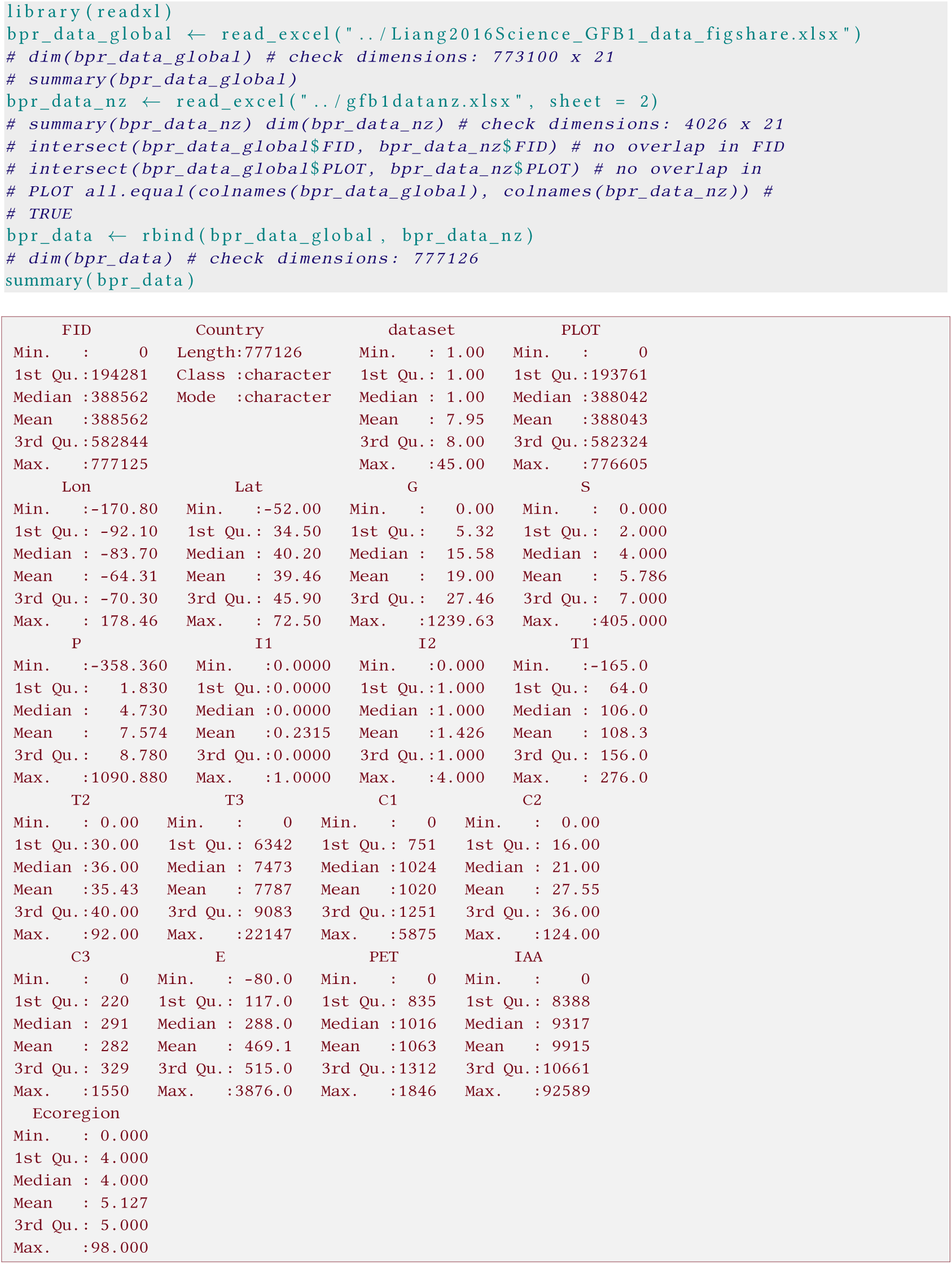

The meaning of the different column titles is provided in the meta-data to the figshare download. Most importantly, the response, productivity [m^3^ha^-1^y^-1^] is given in column P, and the number of tree species observed in a plot is given in column S. As we can see from the summary, productivity ranges from −360 to 1090 m^3^ha^-1^y^-1^, with most data between 1 and 9 m^3^ha^-1^y^-1^. Species numbers range from 0 to 405 (we cannot provide a density here, as plot sizes are not provided and range from < 100 to > 1100 m^2^, only the 95% sampling interval was provided in the paper), with most plots between 2 and 7 species.

Liang *et al.* (2016) excluded plots with harvest volume > 50% of stocking volume (page 5), but do not provide the IDs of these plots. We thus excluded all plots with negative productivity and also those with productivity of 0 (as our model including them has a very different AIC to those reported by the authors).

Additionally, we remove plots with a species richness of 0, basal area of 0, those with a T3-value of 0 (temperature seasonality as standard deviation times 1000, making a value of 0 indicating no data, rather than no seasonality), IAA (indexed annual aridity times 10^-4^)-values of 0, and manually assigned four plots with an ecoregion value > 13 based on their coordinates in googleEarth. (Although there are 14 ecoregions in WWF’s definition, the 14th in this data set was wrongly coded in the data and in fact plots were in rainforests.)

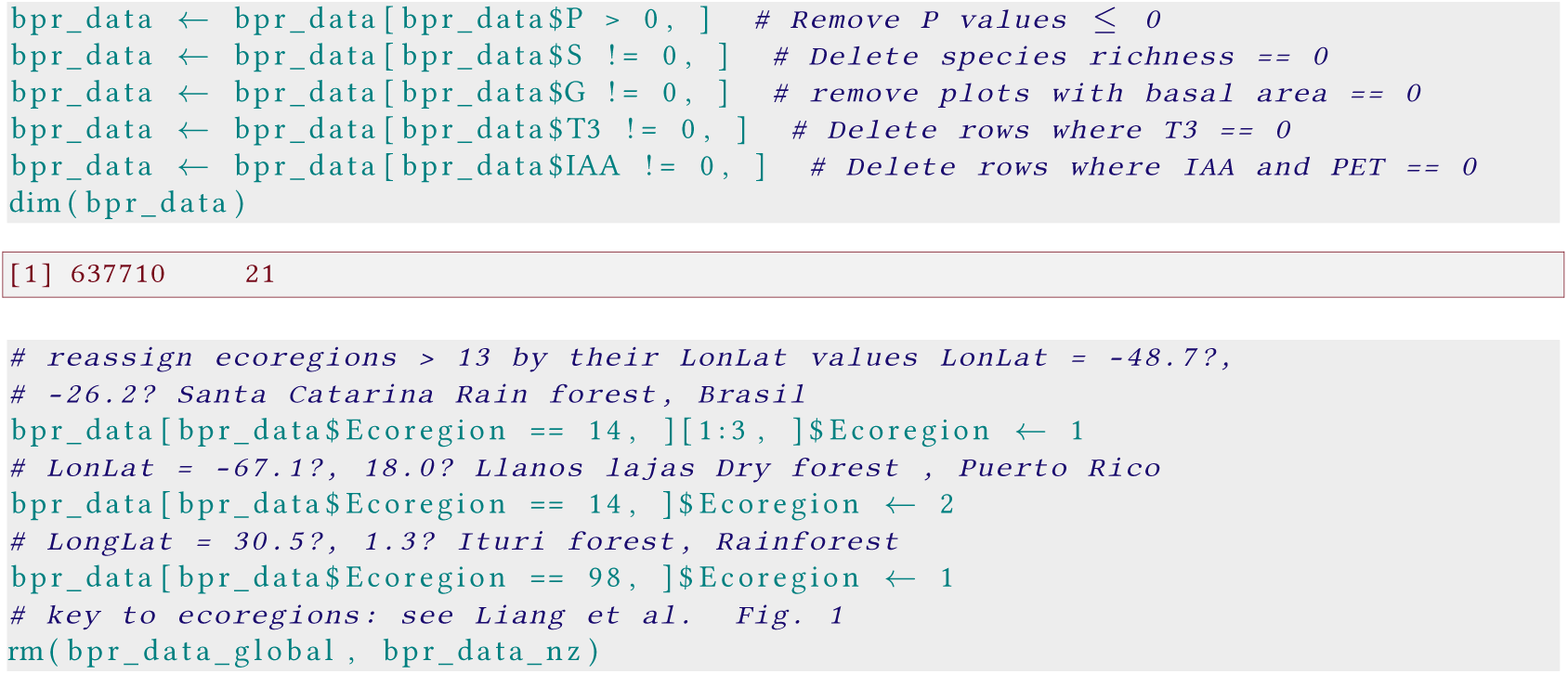

Some 1139 plots were recorded as ecoregion 0, which, when plotted, were distributed along the coasts across the world, i.e. belonging to very different ecoregions. We have no idea why these are classified as “0”, but excluded them rather than manually and labouriously assigning them to appropriate ecoregions. Some plots were recorded repeatedly in the database (e.g. plot 0 occurs 370 times), but we kept them all in the data, as FIDs were unique, and so are their geographical coordinates.

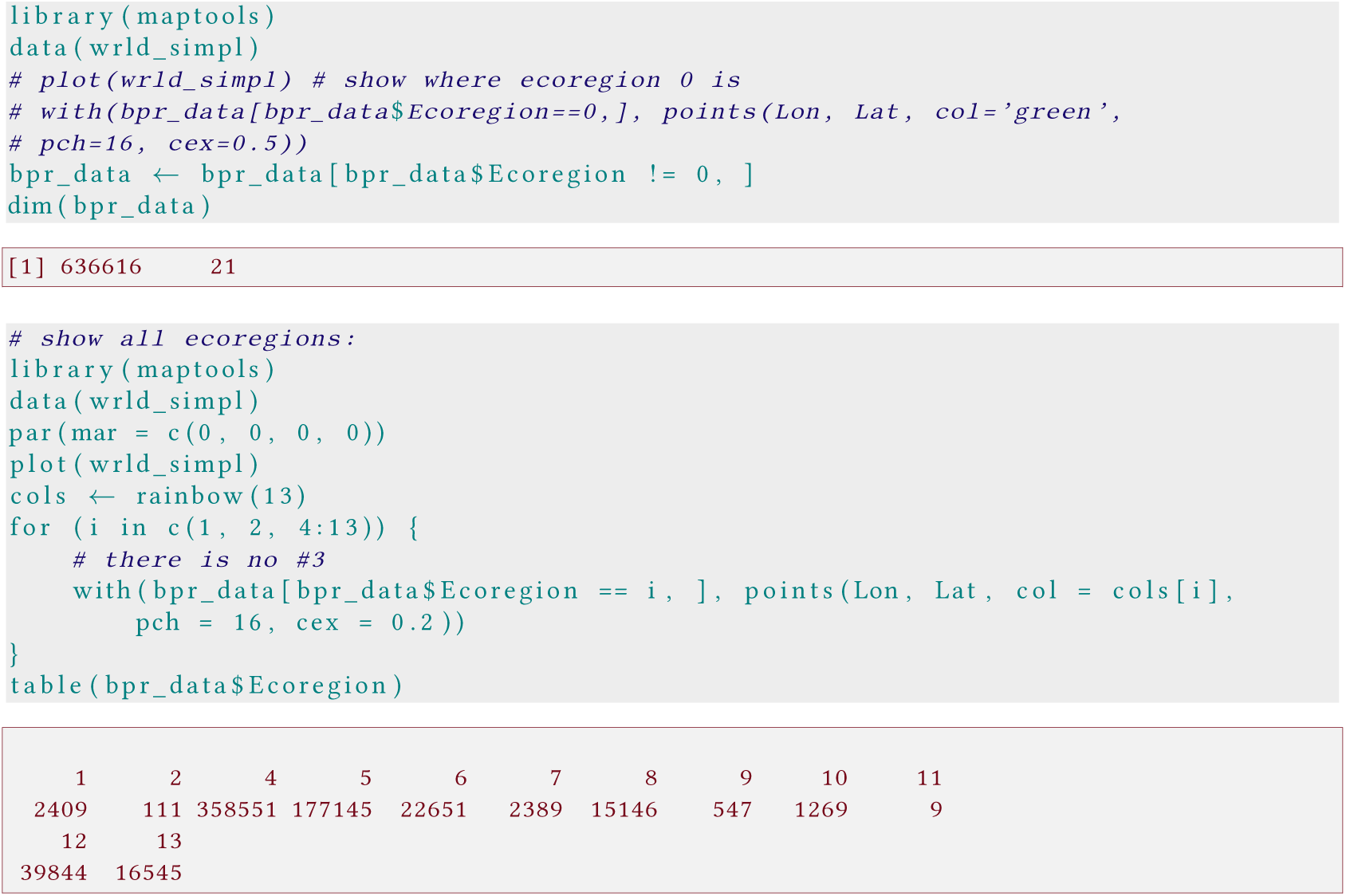

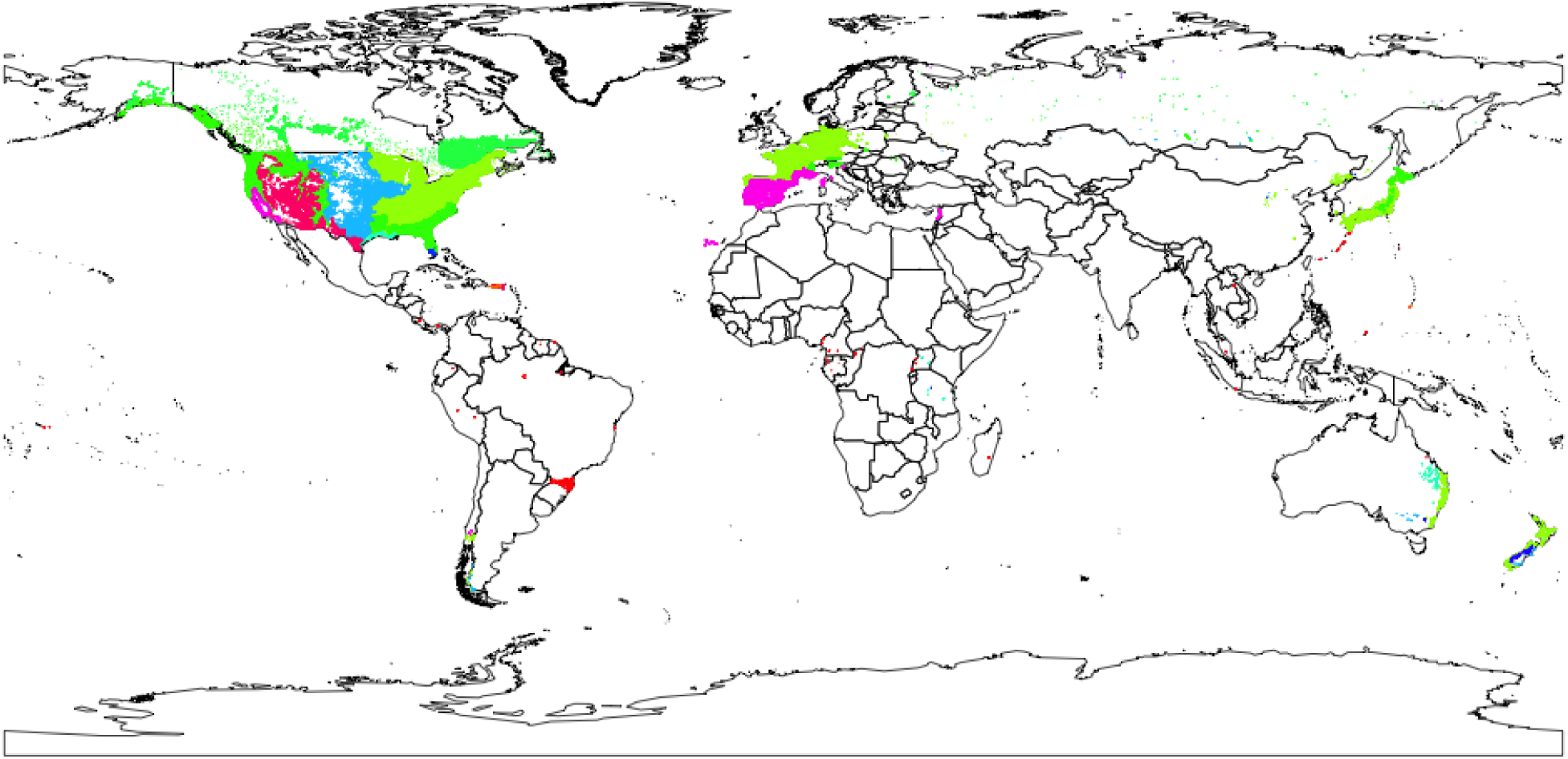

At the end we are left with a data set of 636616 rows, heavily relying on the forest inventory data of temperate forests (ecoregions 4 and 5 comprise 615576 plots, i.e. 97% of all data points). Since for tundra, ecoregion 11, only 9 data points are available, we exclude it from the biome-specific analyses lateron.

Before analysing these data, let’s have a look at them, separately for each ecoregion:

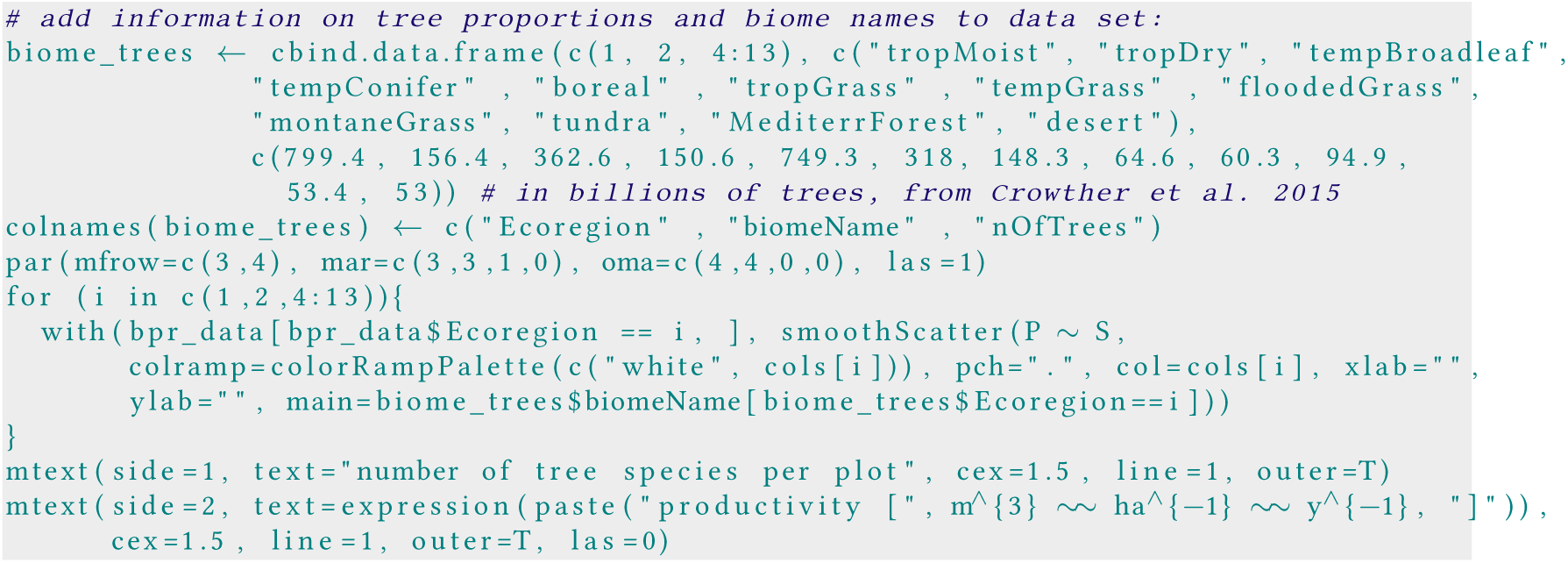

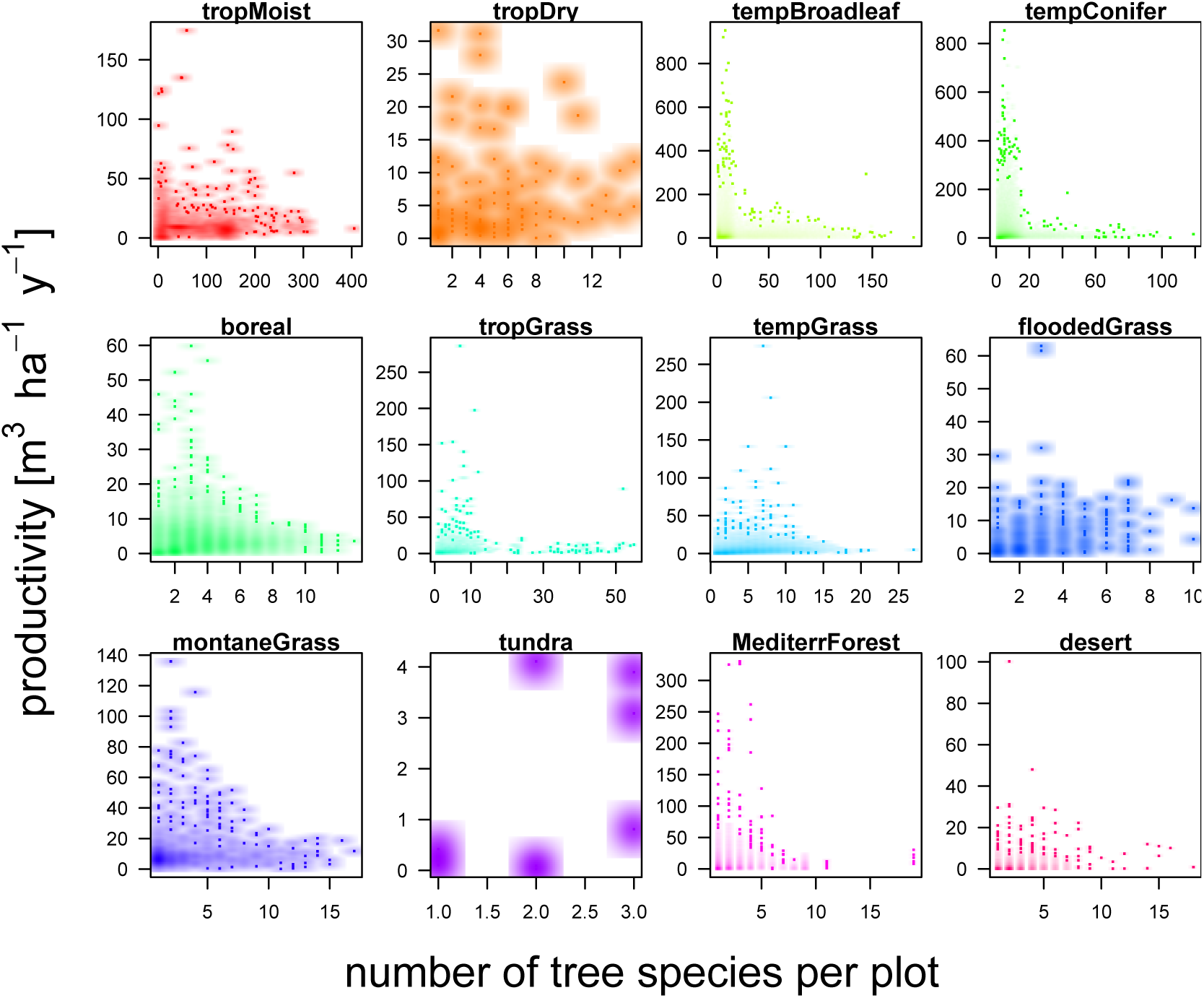

It is visible, at first glance, that these data are suspicious. The highest volume productivity ever measured in situ (rather than estimated from inventories), in a fertilised Eucalypt plantation in Brazil, was around 30 m^3^ha^-1^y^-1^ (Stape *et al.*, 2010). In this data set, the highest value is over 1000 m^3^ha^-1^y^-1^, and there are 23561 points with values larger than 30 m^3^ha^-1^y^-1^ (over 15000 of those are from the US, but Germany, France and Japan also contribute thousands of data points, respectively):

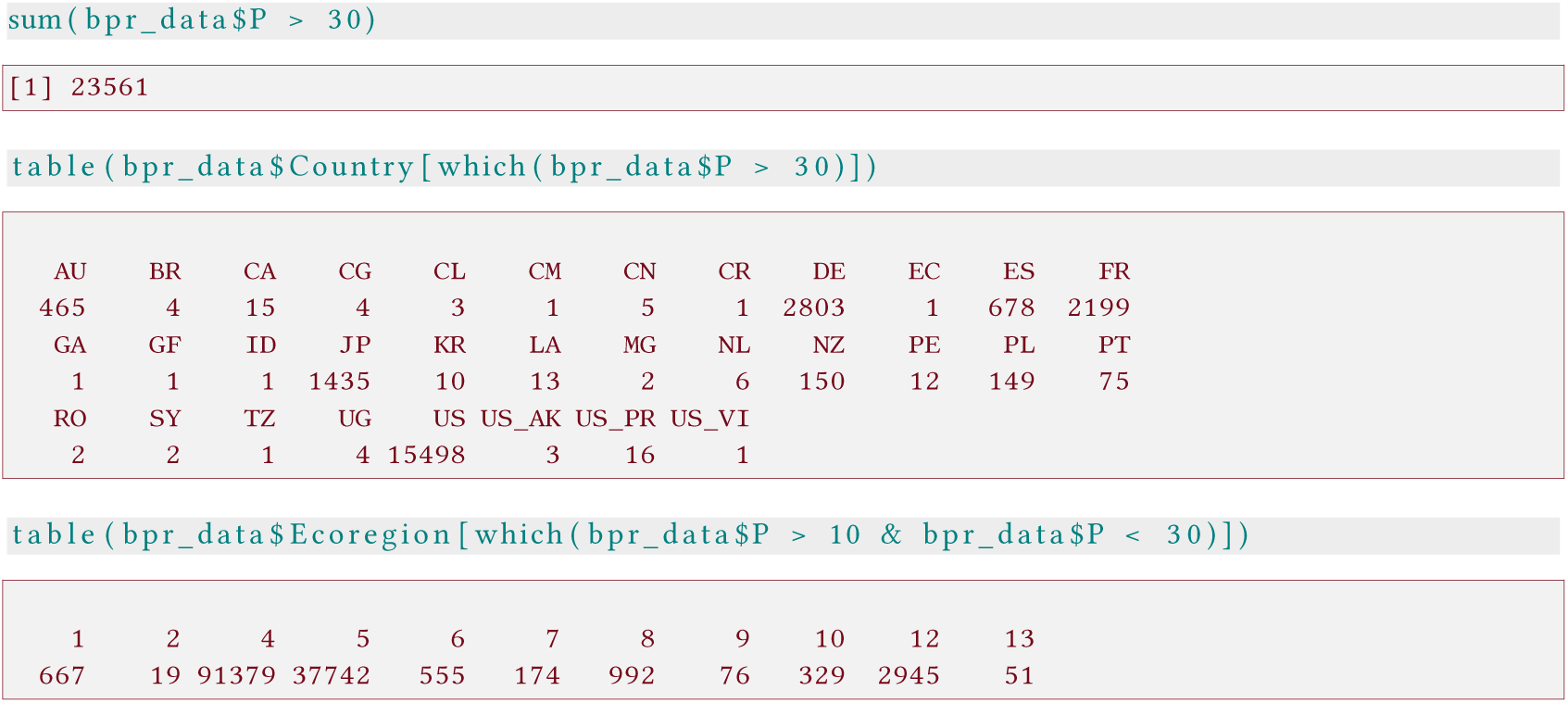

Moreover, as the last figure shows, some unlikely ecosystems contributed to high productivity sites (even after excluding obviously wrong data points): tundra, Mediterranean forest and deserts contributed dozens to thousands of plots with productivity exceeding 10 m^3^ha^-1^y^-1^. It is possible that the authors did remove some of these plots before the analysis without mentioning it explicitly in the methods (p. 5: “Intensively managed forests with harvests exceeding 50 percent of the stocking volume were excluded from this study”; there is no indication in the data, which plots are concerned).^2^

Nevertheless, as we are able to reproduce the analysis to a large extent (see below), we assume that many of these data points were still in the data set analysed.

#### B.2 Spatial model

Liang et *al.* (2016) employ a Generalised Least Squares (GLS) approach to accommodate spatial autocorrelation. Since the GLS is rather time-consuming, and it cannot handle data sets beyond a few thousand data points (depending on computer memory), they took random subsamples of 500 plots from the above data and ran the GLS on that subsample.^3^ For each such subsample, a new variable Š was computed as 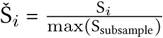. The actual model is at the log-log-scale (i.e. both P and Š were log-transformed). Šis interpreted as “relative species richness” and is claimed to “facilitate inter-biome comparison” (p. 7)^4^.

So the actual steps of the analysis are:

1. Draw a random subsample of 500 data points (without replacement).
2. Compute Š for this sample.
3. Log-transform P and Š.
4. Run a GLS with log(P) as response and log(Š), G (basal area), T3 (standard deviation of temperature), C1 (annual precipitation), C3 (precipitation of warmest quarter), PET (potential evapotranspiration), IAA (indexed annual aridity) and E (elevation) as predictors^5^. The spatial distances are implicitly computed from geographic coordinates using the corSpher correlation structure^6^. Since coordinates of the released data were rounded, we jitter them slightly to avoid distances of zero.
5. Store model estimates and repeat.

While Liang et *al.* (2016) run 10000 bootstraps, we only do 50, as the results do not change substantially anymore after that. (Note that our code is not made for speed but for intellegibility.) Also, as the authors kept the nugget and range fixed in their final run, we use these values here (0.8 and 50, respectively).

As illustration, here is a single model run:

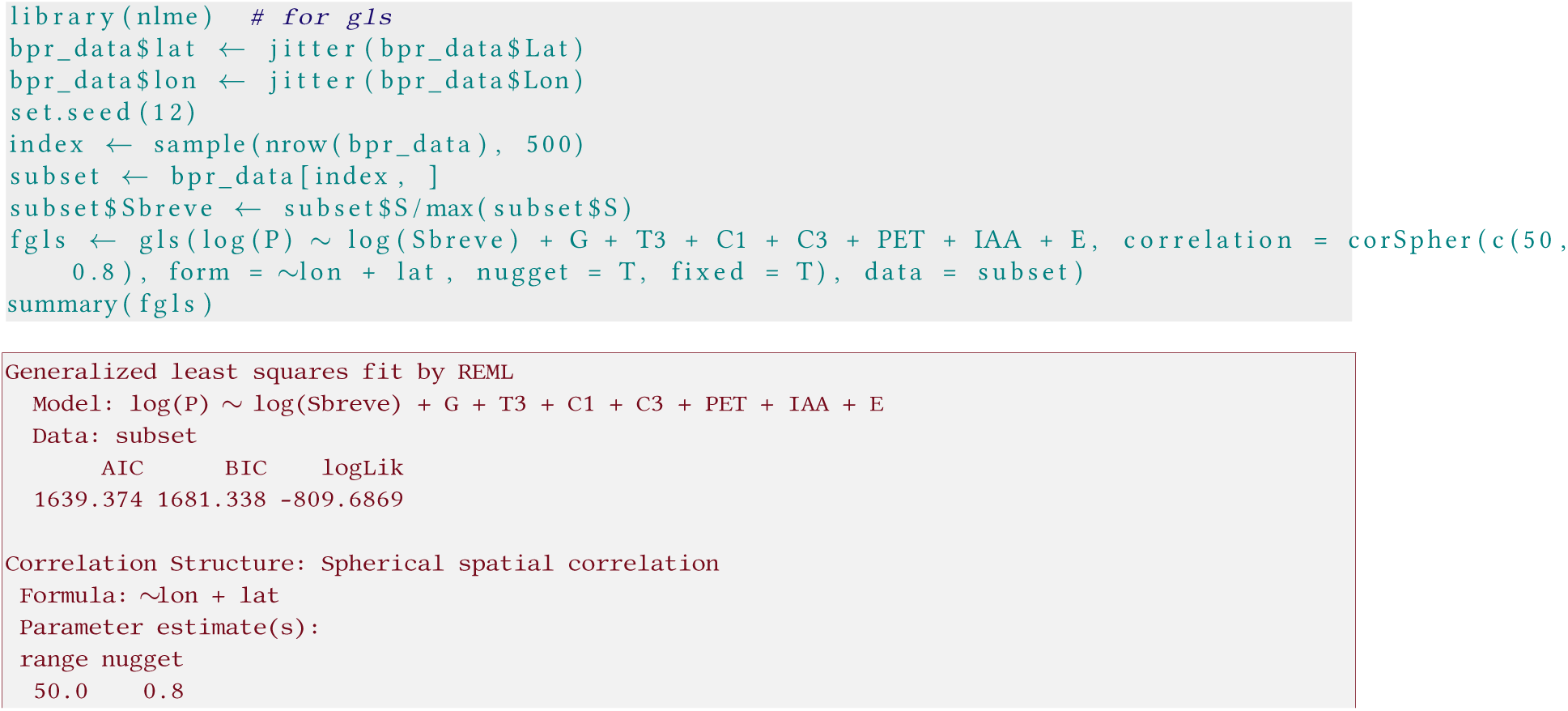

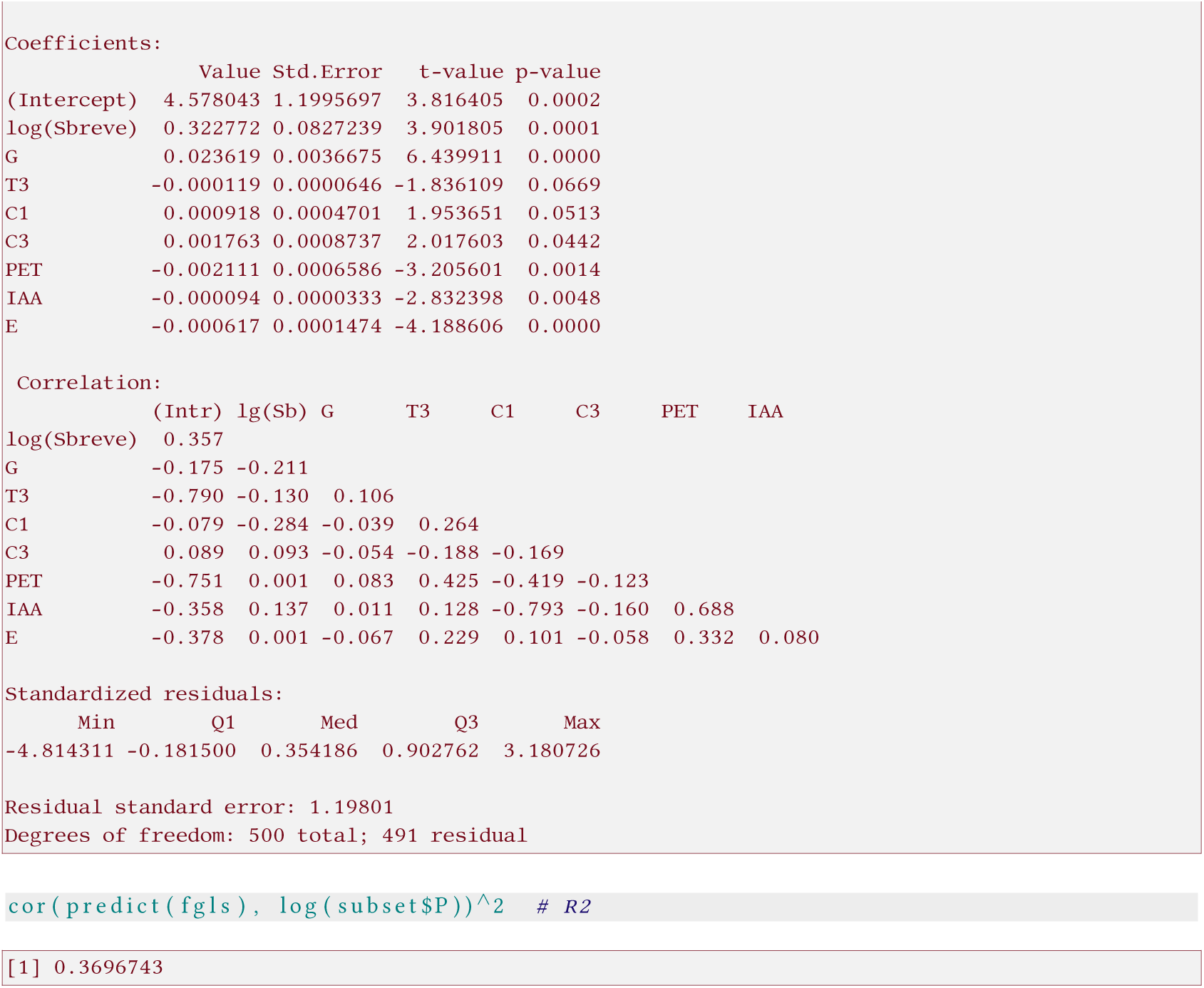

The part after setting the seed will be repeated below, and from the summary we only keep the coefficient estimates.

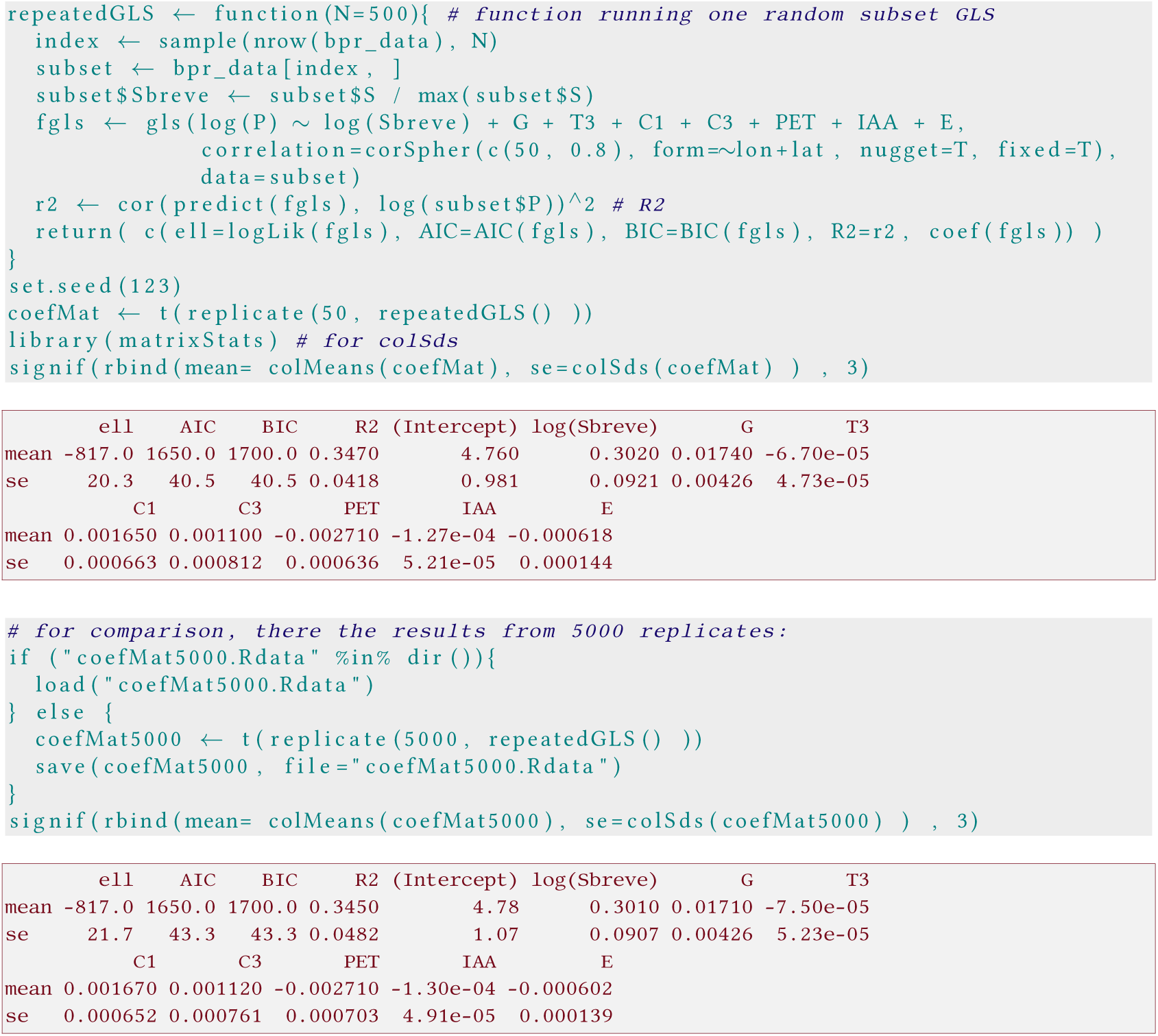

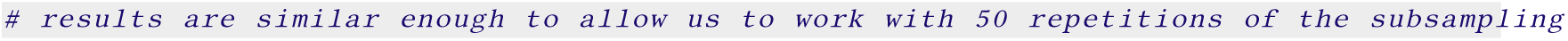

This approximates the results of Liang *et al.* (2016)’s Table 2, with the exception of the standard errors for the model coefficients, which Liang computed as incorrectly as standard deviation divided by the square root of the number of replicate models: correctly computed, the standard deviation of a bootstrap is the standard error of the statistic (here the goodness of fit measures and the coefficients). Hence, the row “SE” in the authors’ table 2 must be multiplied by 100 to get to the correct values.^7^

**Table 2:** Parameters of the *global* geospatial random forest model in 10,000 Iterations of 500 randomly selected (with replacement) GFB plots. Mean and SE of all the parameters were estimated by using bootstrapping. Effect sizes were represented by the Akaike information criterion (AIC), Bayesian information criterion (BIG), and generalized *R*^*2*^ (*G-R*^*2*^), Const, constant.

| | Loglik | AIC | BIC | $G-R^2$ | const | $\theta$ | G | T3 | C1 | C3 | PET | IAA | E |
| --- | --- | --- | --- | --- | --- | --- | --- | --- | --- | --- | --- | --- | --- |
| Mean | -761.41 | 1546.71 | 1597.08 | 0.354 | 3.816 | 0.2625243 | 0.014607 | -0.000106 | 0.001604 | 0.001739 | -0.002566 | -0.000134 | -0.000809 |
| SE | 0.54 | 1.10 | 1.13 | 0.001 | 0.011 | 0.0009512 | 0.000039 | 0.000001 | 0.000008 | 0.000008 | 0.000009 | 0.000001 | 0.000002 |

Finally, we plot this relationship with the “correct” error bars. As parameter estimates are correlated, we cannot simply use the above standard errors to compute an error interval around the line. Instead, we predict with each of the 50 parameter sets the curve and compute 95% quantiles from them as confidence intervals. ^8^

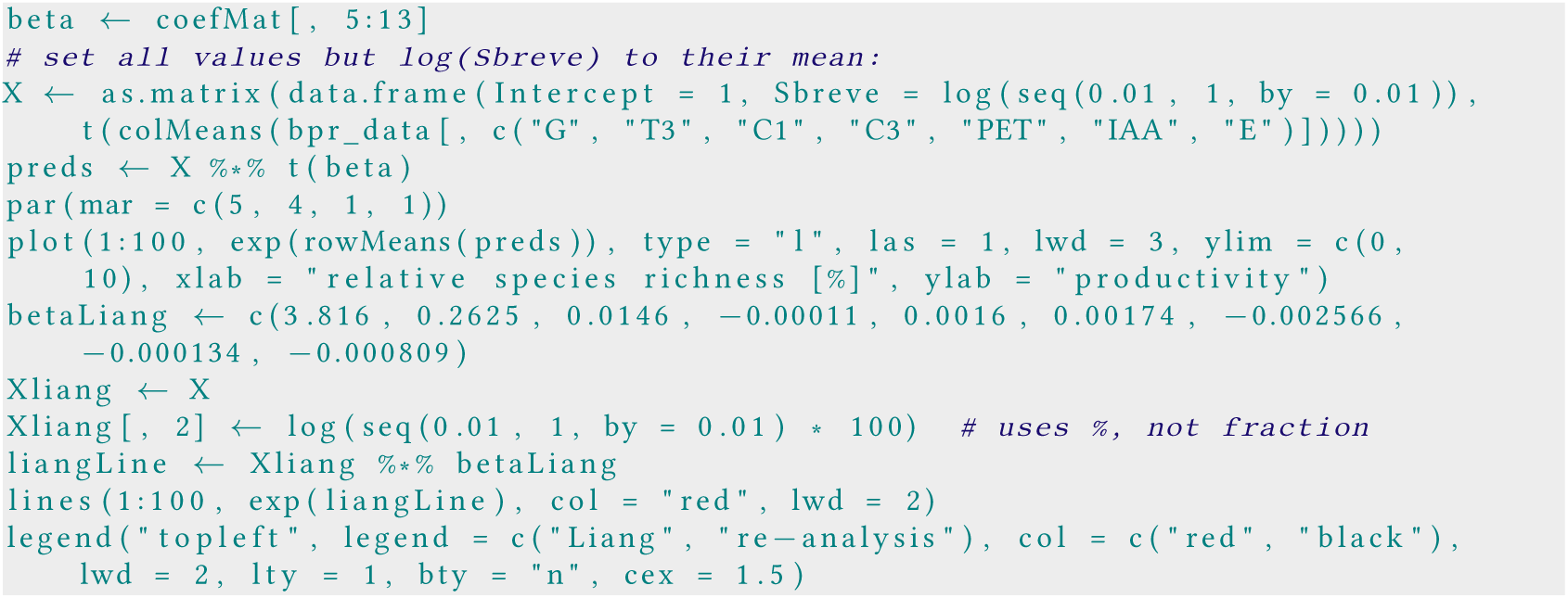

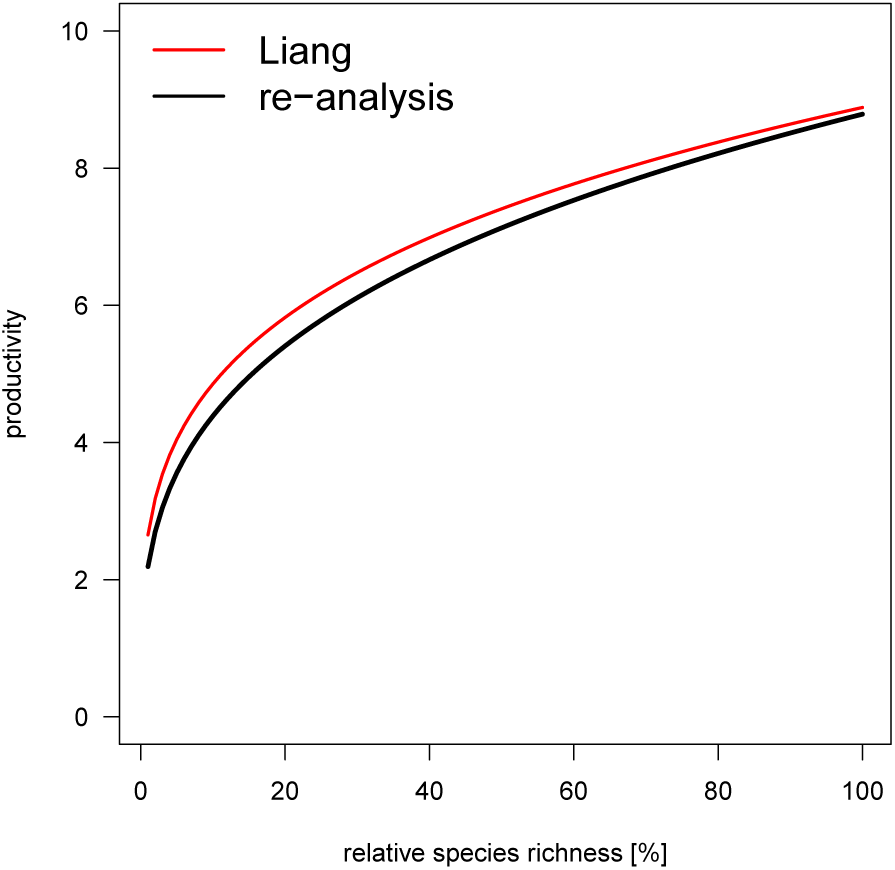

We were thus able to reproduce the analysis of Liang et al. relatively faithfully, at least with respect to the expectation (we shall return to the error envelop in due course).

Since their log-likelihood is 50 units higher, we suspect that in their analysis further plots were excluded.

### C Corrections and improvements

In summary, we have been able to reproduce the analysis of Liang *et al.* (2016).

During this first step, we may already have noticed some issues with the analysis. Here is our list of things we improve on in the next sections, starting with the most important ones first:

1. Evaluate the effect of unreasonably large values of productivity on the result.
2. Aggregating analyses of subsets, which comprise only a permille of the full data set, may introduce a bias relative to the full dataset. The mechanism behind this bias is the same that we criticised as first point, i.e. the computation of Š on the basis of highest tree species richness in the subset, thereby ranging Š differently for each subset.
3. Minor point: The GLS can easily accommodate more than 500 points, so we can run sub-samples of 1000, 2000 and 5000 points. This will slightly increase the chance of plots being relatively close to each other, and hence allows for a better estimation of the short-range spatial autocorrelation, possibly improving the estimates for nugget and range of the variogram.
4. Stratified draws of plots in proportion to the area this ecoregion’s forests cover on the world. Currently only some 2500 points (out of 636000) are representing the tropical forests (labelled 1 and 2 below), despite their proportion of the world’s forests being over 30%.

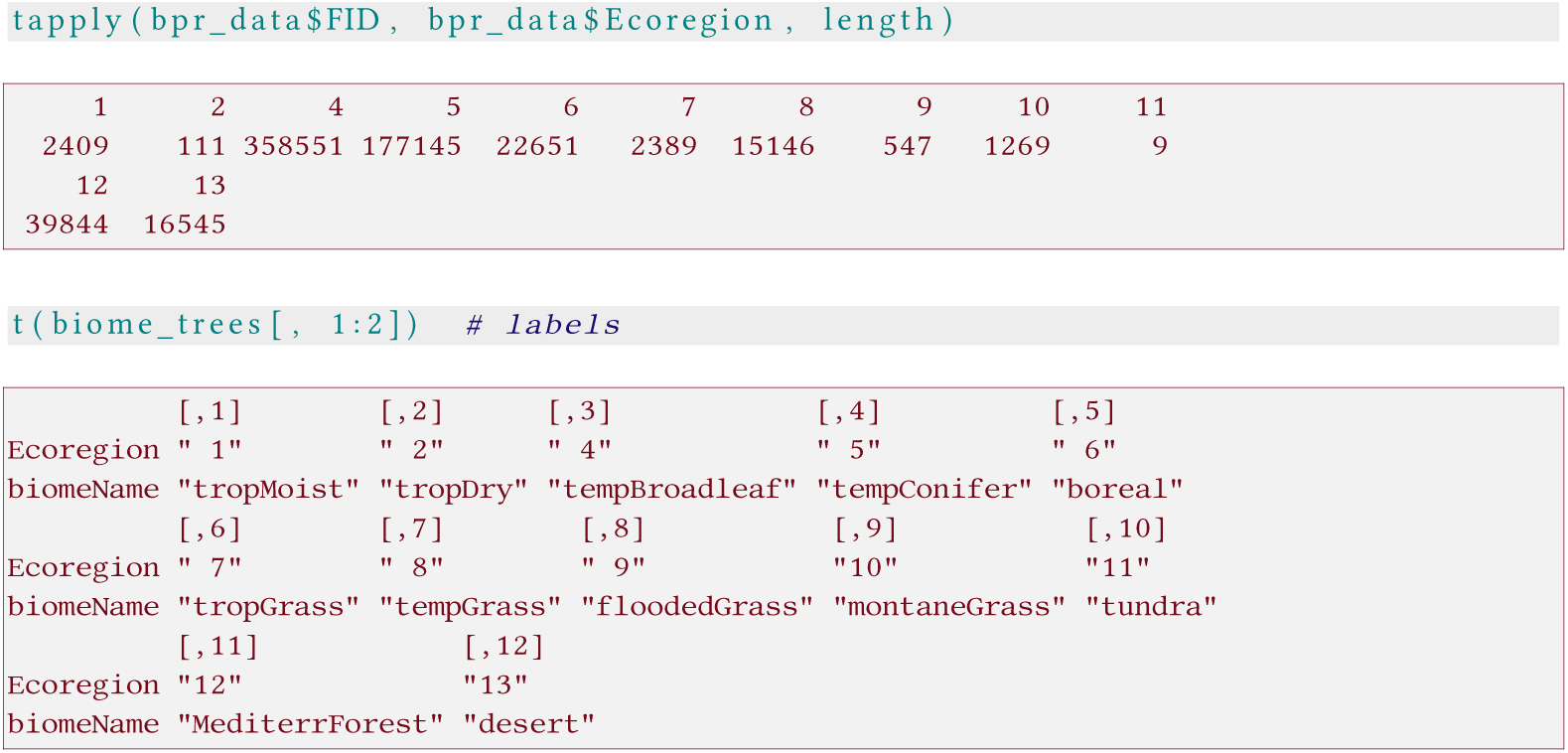
5. The corSpher-structure used by Liang *et al.* (2016) suggests, in its name, that it would be appropriate for data on a sphere. That is not the case. The spherical model is only the name for a different shape of how spatial autocorrelation decreases with distance (Pinheiro & Bates, 2000), but it does not correct for the fact that on a sphere geographic distances cannot be computed from coordinates as Euclidean distance. To do so, we have to compute the so-called great-circle (or orthodromic) distance.
6. Calculate relative species richness Š relative to what is the maximal species richness *in that region.* Currently, the definition of relative species richness is not intuitive. Since only the tropics have plots with dozens to hundreds of species, Š is effectively representing a gradient from cold to tropical plots.^9^

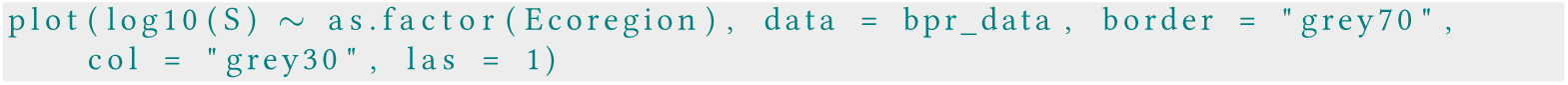

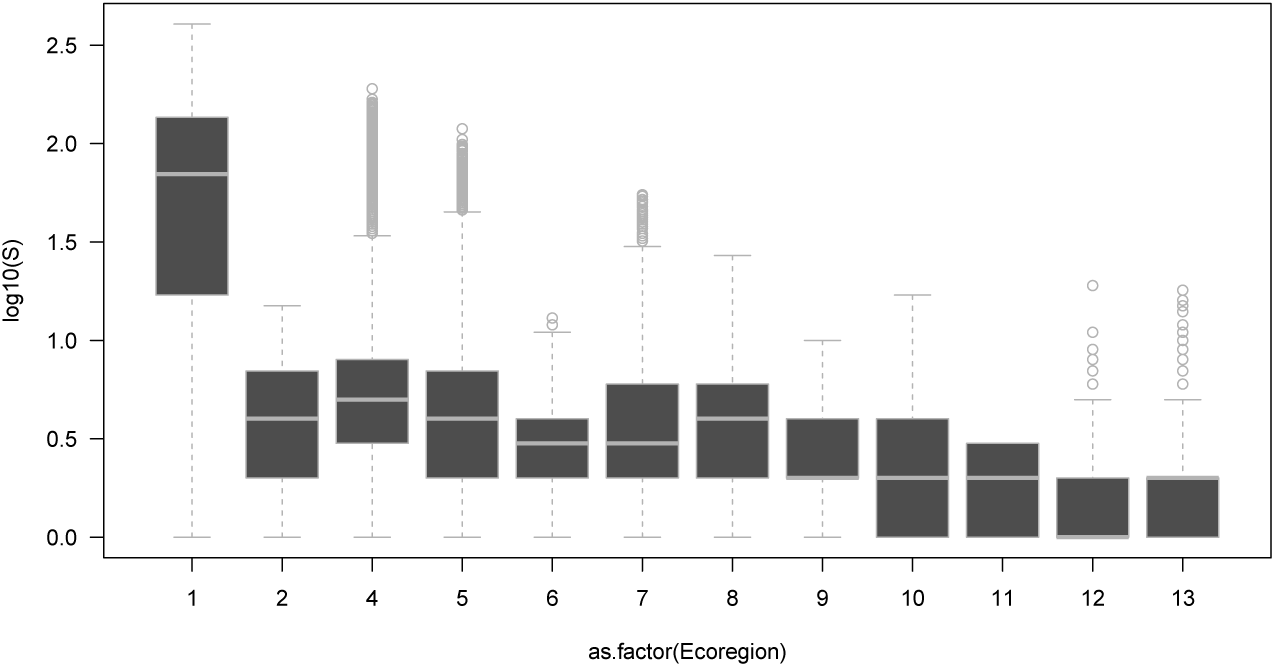 Thus, the so-called species richness gradient is in fact a latitudinal gradient. This is, in our opinion, a serious problem and leads to fundamental misinterpretation of the results. We propose to compute Š differently, namely relative to what is possible for a given plot. We could choose the maximal value for an ecoregion to scale tree species richness of each plot, or even more locally, the maximal species richness observed in, say, 100 km radius around a plot. This would lead to plots from any ecoregion being able to occupy the upper end of the relative species richness gradient, if they are species rich *relative to their wider neighbourhood.*^10^

In the following, we demonstrate the effect of each of these improvements separately, always comparing it to our analysis of the data presented above. Arguably, as the data set changed, we may have to re-estimate the coefficients of the spatial autocorrelation, which we did not.

#### C.1 Do implausible productivity values distort the analysis?

As shown above, some 23,000 data points, around 4% of the entire data set, have productivity values above the largest measured in the field. The probably reason is that forest inventories measure with a certain error, and if the error is large relative to the increase in biomass, it is easy to get unrealistically high productivity estimates, particularly in small plots with harvesting.

There are, essentially, two ways to accommodate values that are clearly implausible: (i) omit them, or (ii) give the lower weight in the model. We only investigate the effect of omitting all values where P>30 as the strongest, not necessarily most appropriate, way.

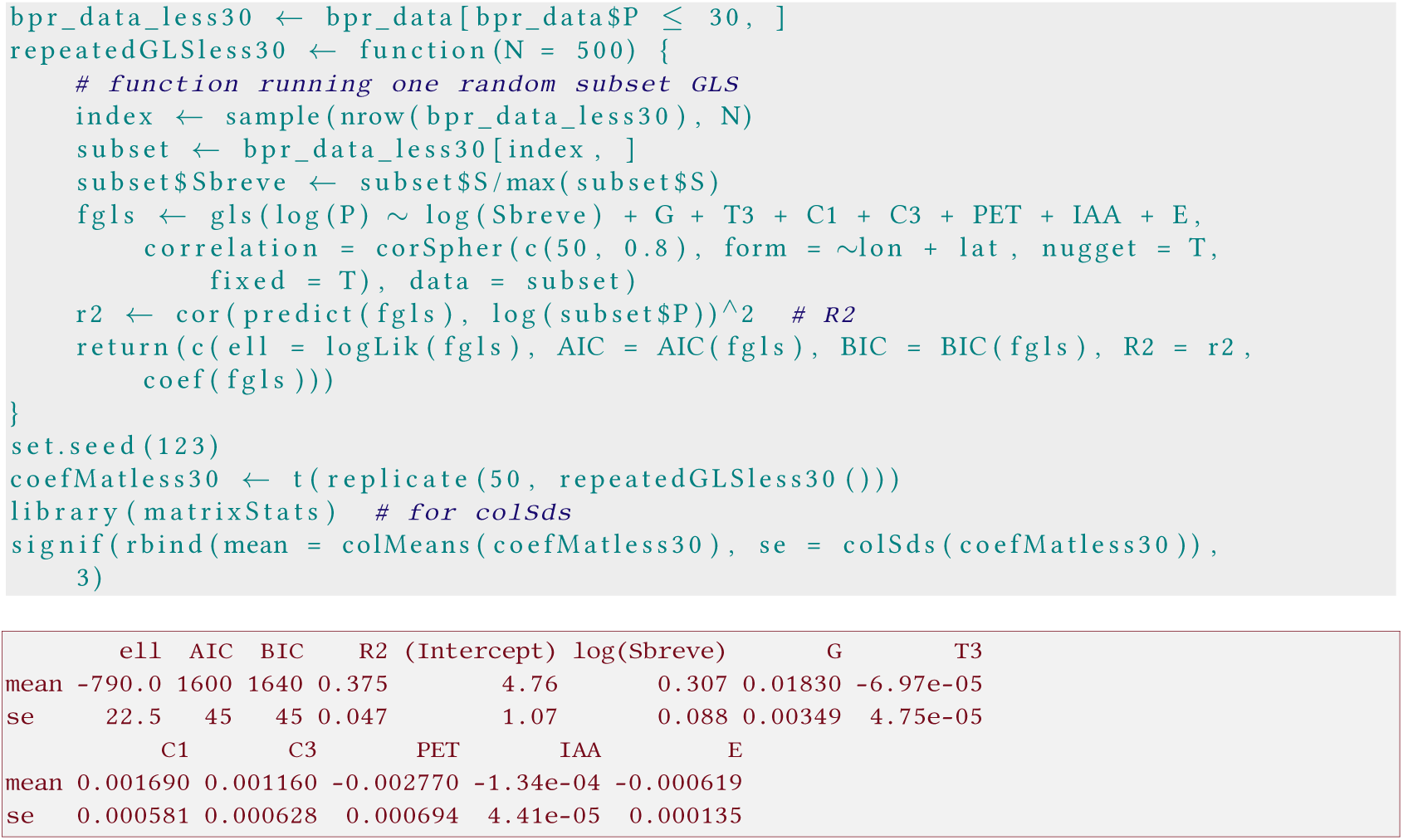

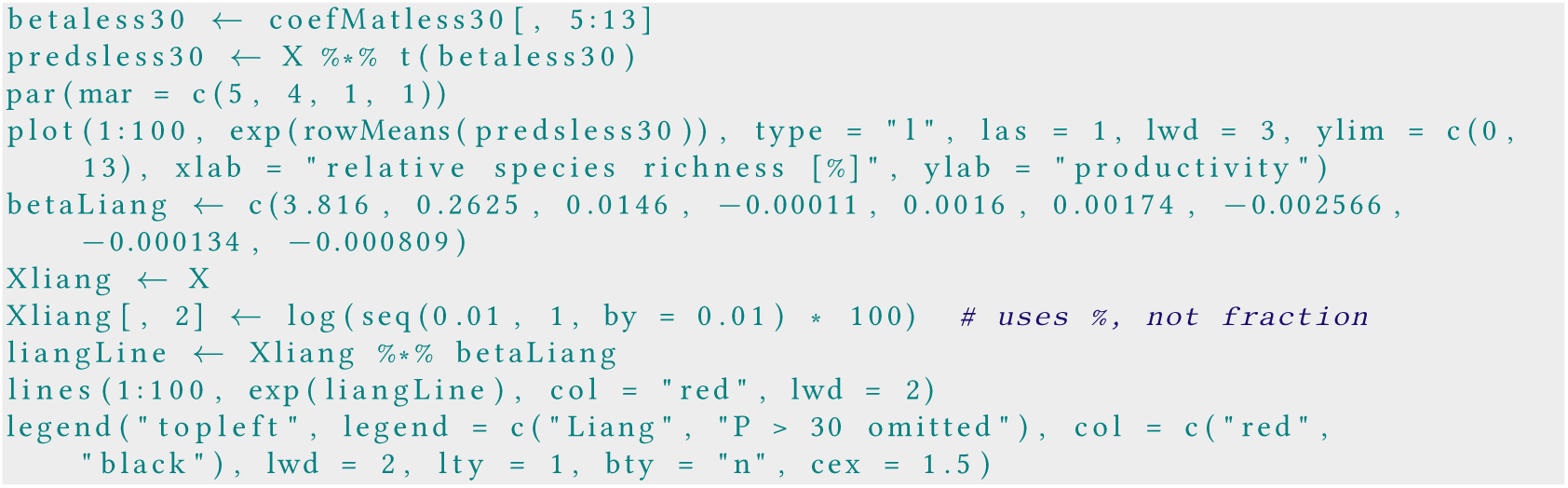

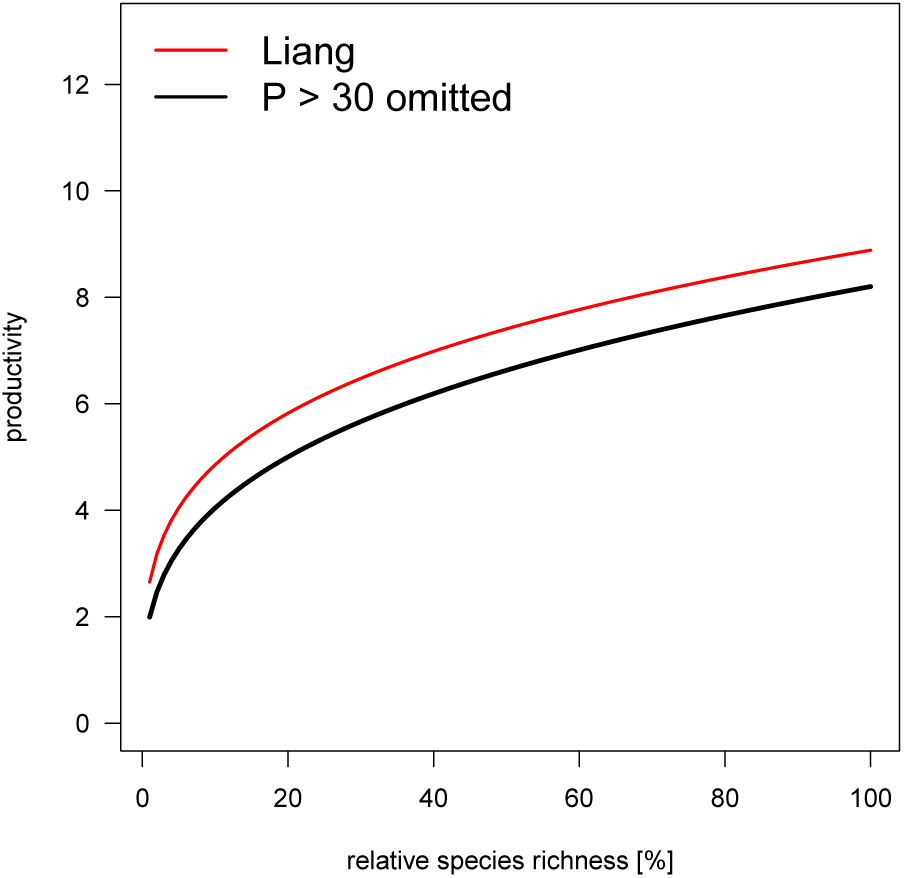

Excluding the high-value plots leads to the expected drop in the absolute values of the TSR-P- relationship, but it does not substantially affect the slope estimate.

#### C.2 Aggregated subsets vs using the entire data set: do parameter estimates match?

As sample size of a regression problem increases, standard errors of model parameters decrease, typically as a function of 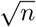. Thus, the 500-data point subset is likely to have much wider error bars than an analysis of the full 636616 data points. How much wider is somewhat difficult to estimate, as data are spatially autocorrelated, making the 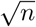 an optimistic estimate. Following the logic of data cloning (Lele *et al.,* 2010), who show that repeating their data set 100 times yields 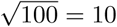 times to narrow standard errors, one can argue that the standard error computed from the subset-models will be 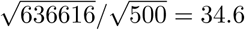 times too wide.

We cannot test this idea on the GLS, as we have no computer at our disposal that can invert a 600000 × 600000 matrix as required by the GLS. Thus, we cannot fit a GLS on the full data set (otherwise Liang et al. would have done that, too). Instead, we use the non-spatial linear model, which is biased due to spatial autocorrelation, as we shall see. Still, if our 35-fold correction worked for the non-spatial regression (the ordinary least square: OLS), we would be happy to use it for the GLS.

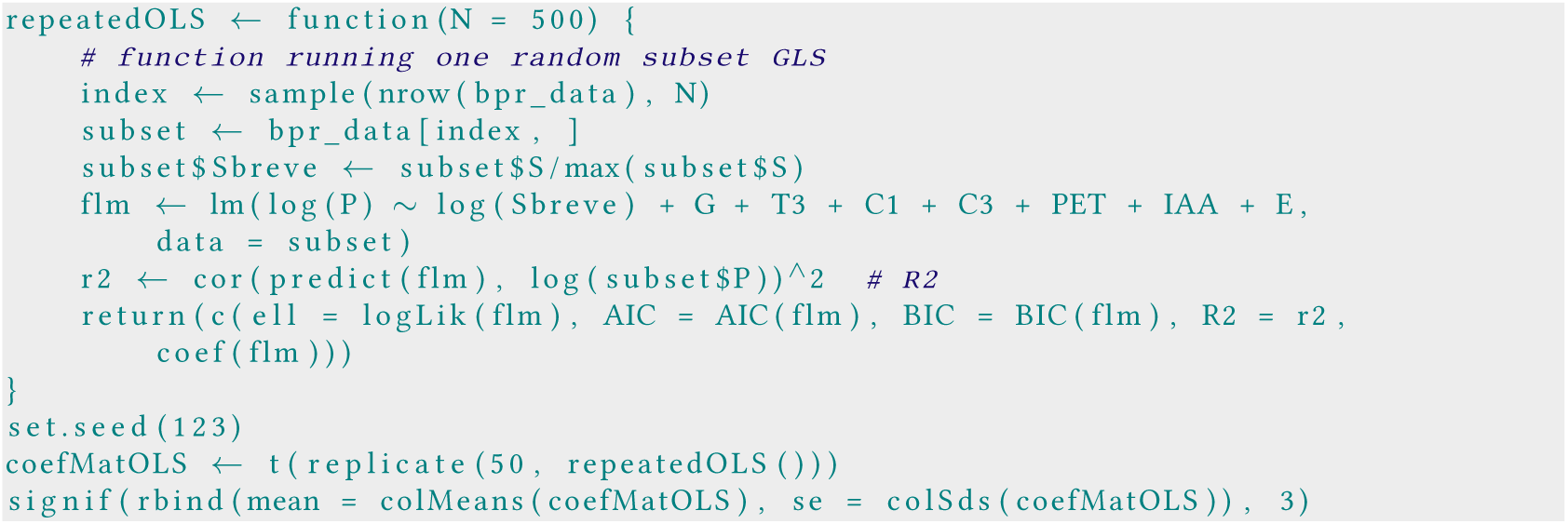

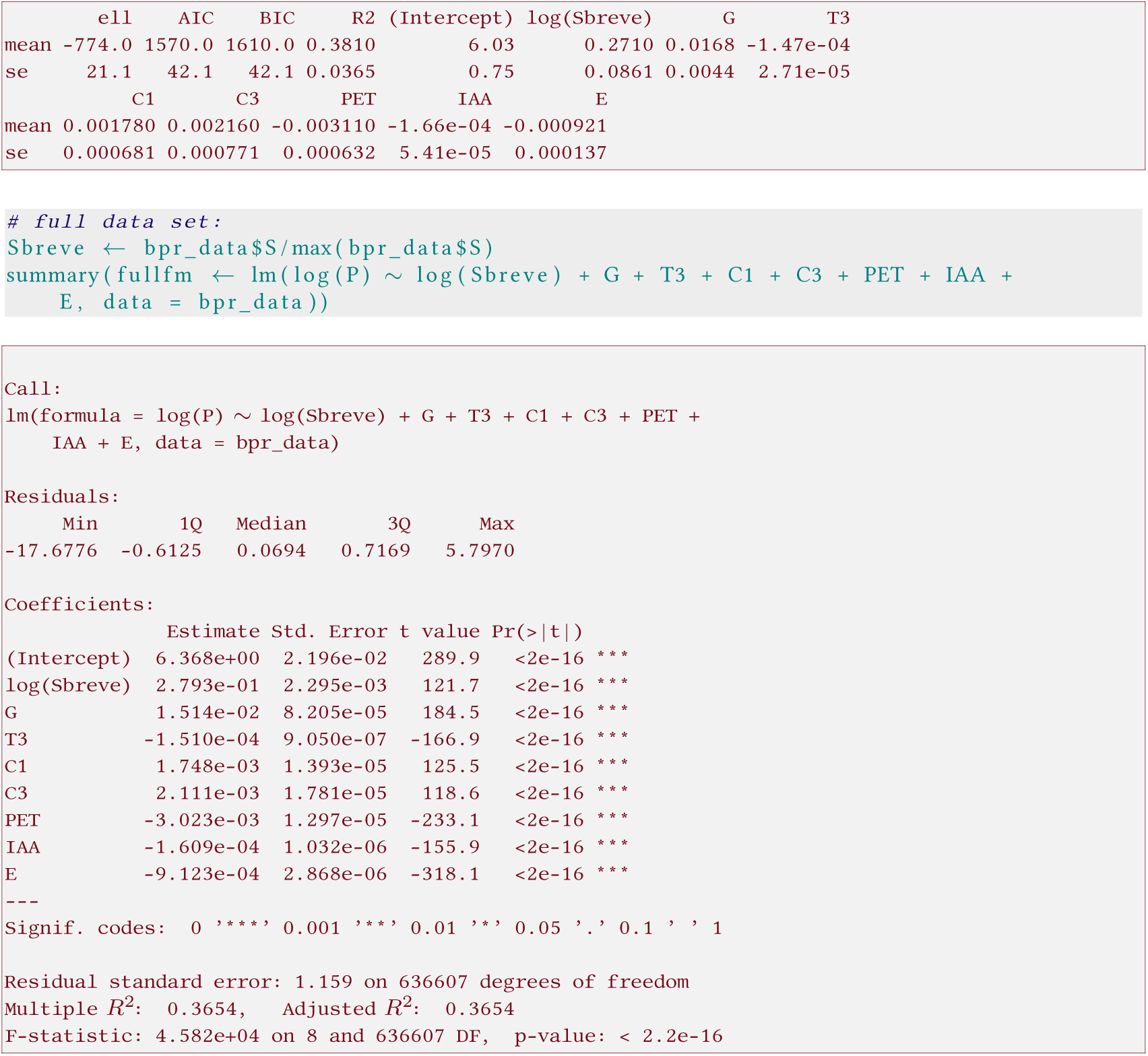

Resampling OLS and full data set analysis yield similar parameter estimates (apart from the intercept). As result, the **estimated effect of increasing “relative species richness” is substantially lower** in the subsampling approach than in the full model. We are confident that this effect will be similarly present in the GLS analysis and return to this issue in the section on increasing sample size of the subsampling.

Let’s plot the results:

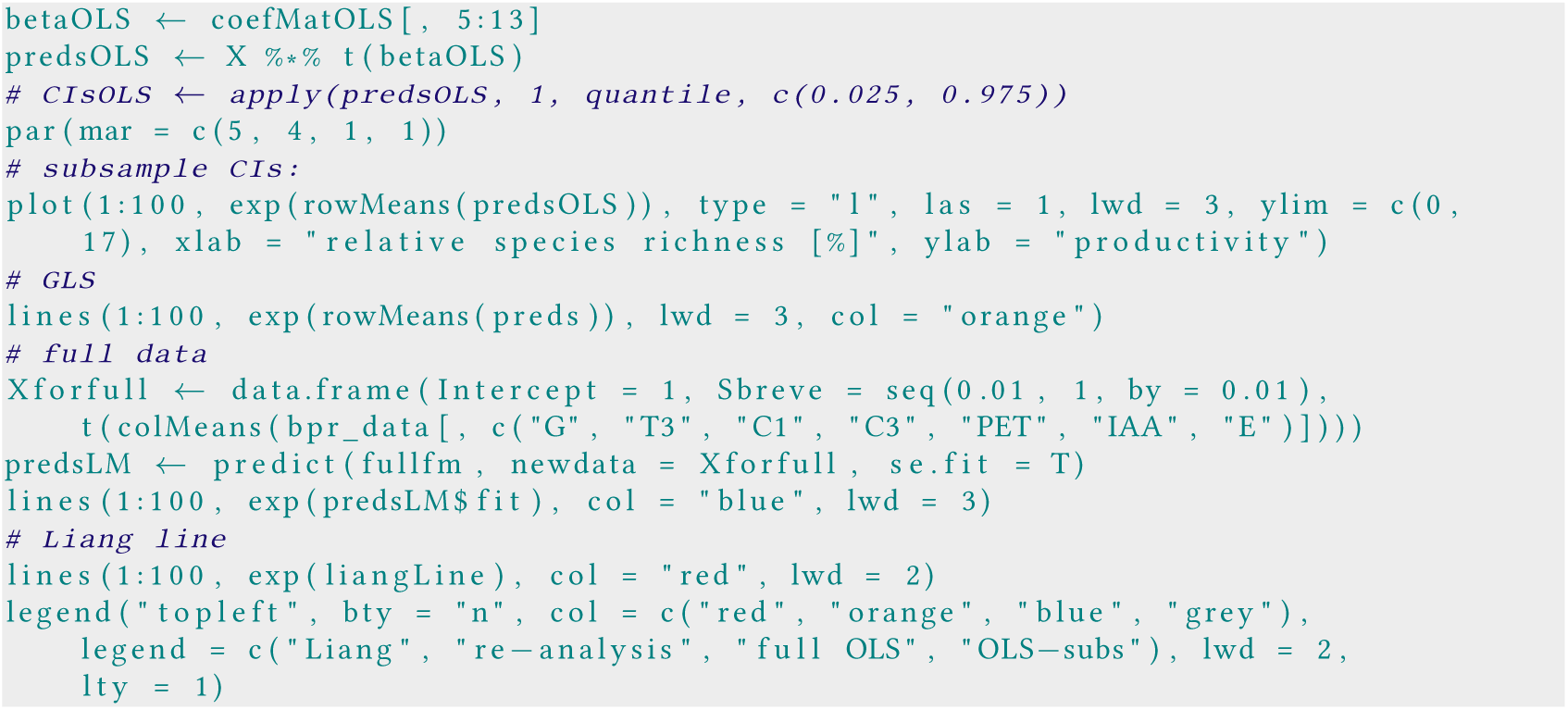

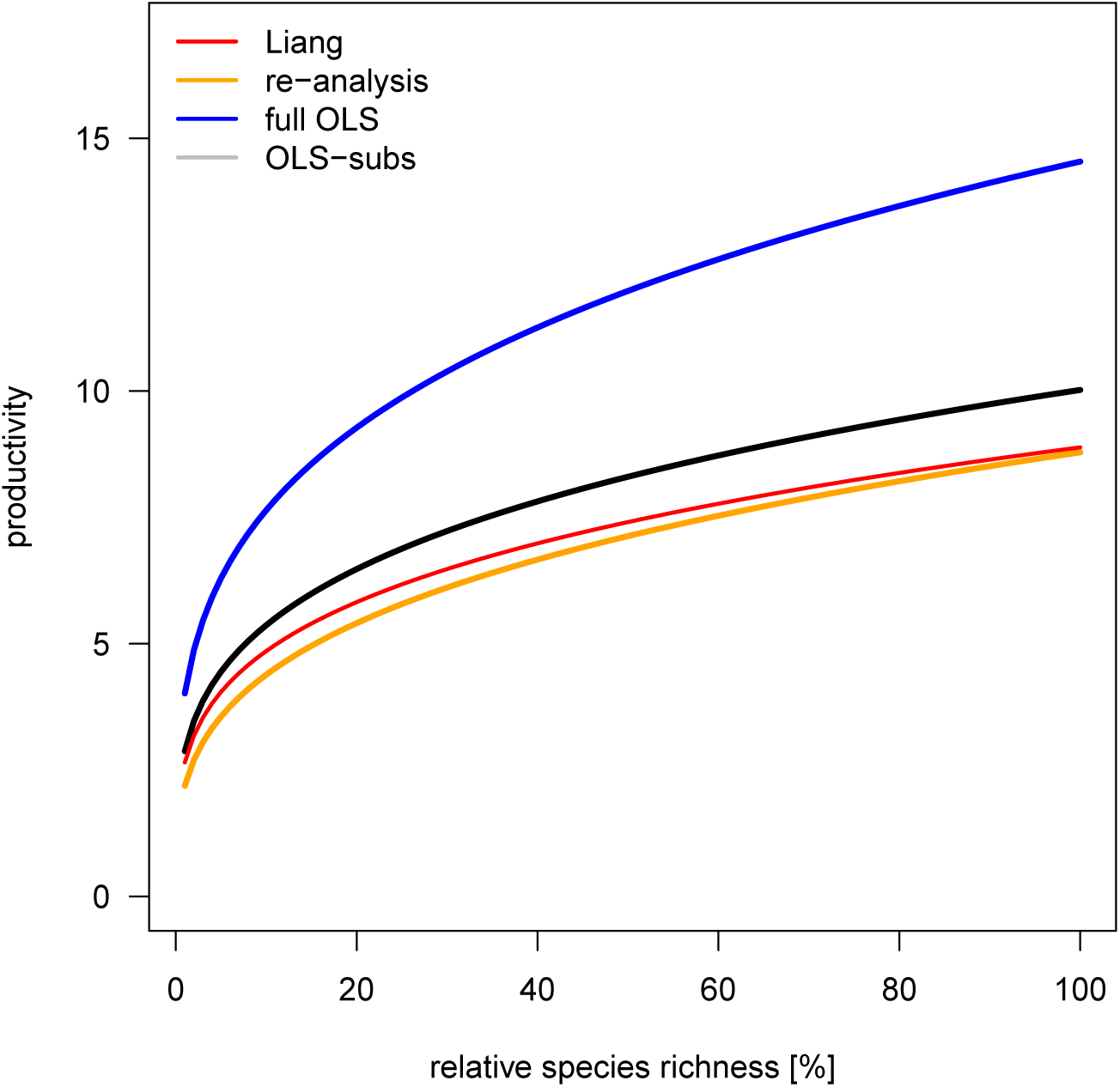

The take-home message of this is: subsampling causes bias, at least for the specific way Š is computed by Liang *et al.* (2016).

#### C.2.1 Scaling standard errors from subsamples to full data

To take the subsampling correction a step further, we now run subsamples of different size and try to identify a relationship between their (subsampling-derived) standard error for 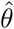 and size of the subsample, *B*. As it happens, this relationship is best presented as a straight line on a log-log-plot.

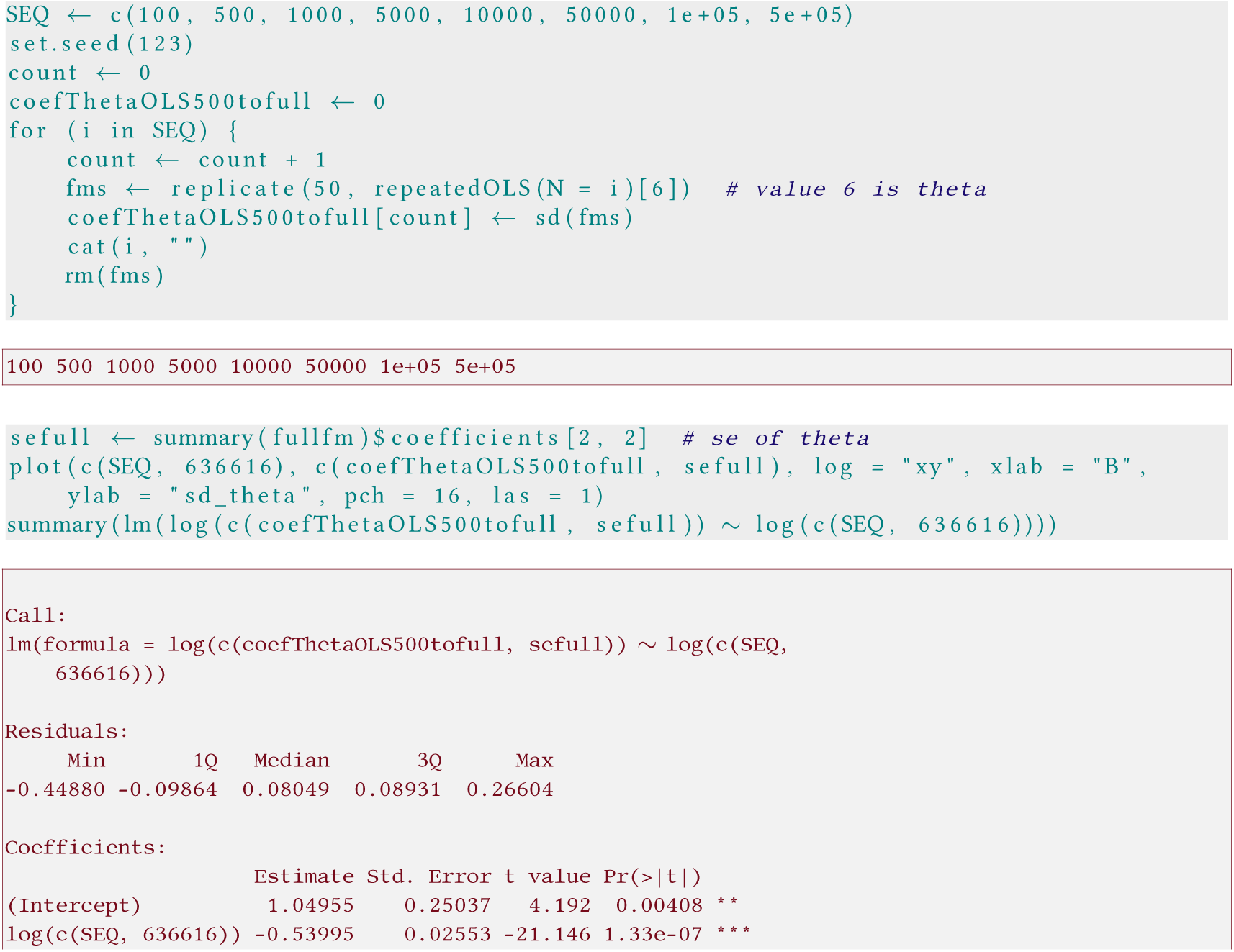

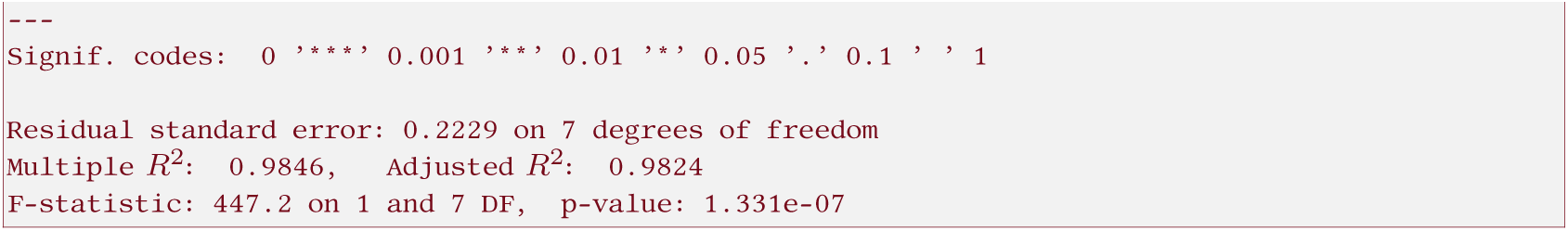

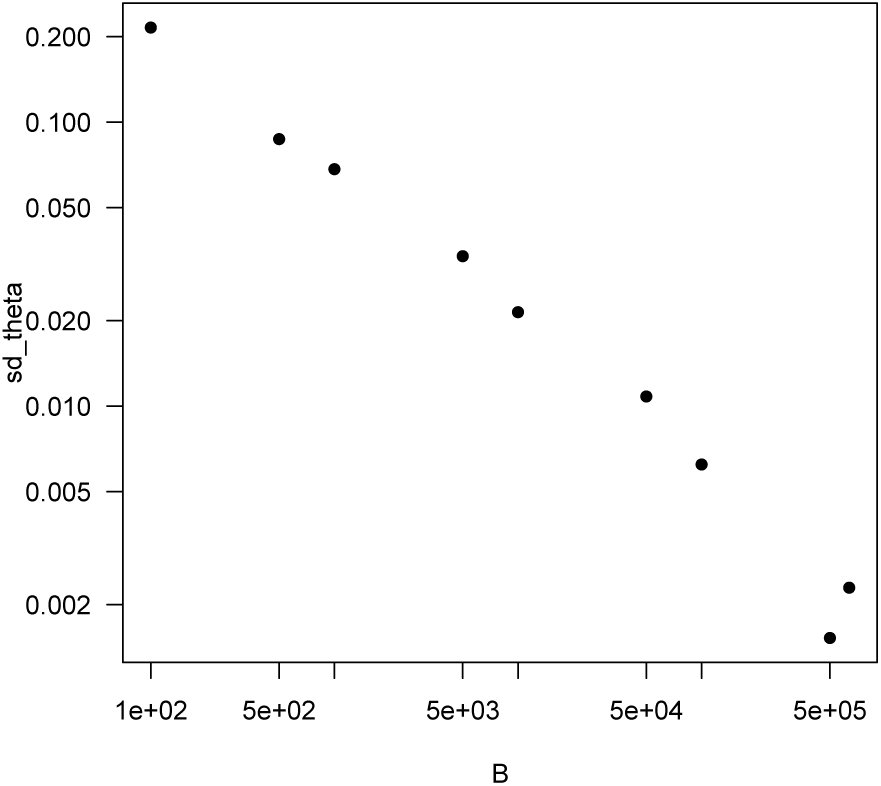

The standard error of the estimate of *θ* clearly decreases nicely as a power-law function of sample size, and fits well to the observed values:

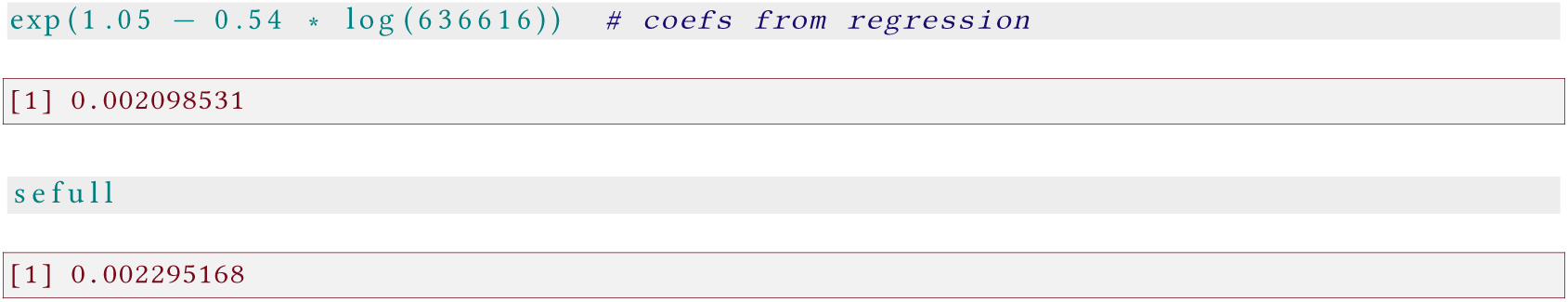

To apply this to the GLS, we have to make the assumption that the same proportionality holds for the GLS as for the 0LS. If so, we can compute the standard error of the GLS, based on the subsampling of size *B* = 500, as follows (as a simple rule of three):

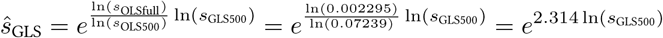

Thus, for the observed standard error of the GLS for subsampling size *B* = 500 we have *s*_GLS500_= 0.0921 (see section B.2), which leads to an estimated standard error of the full model of *ŝ* _GLSfull_ = 0.00263:

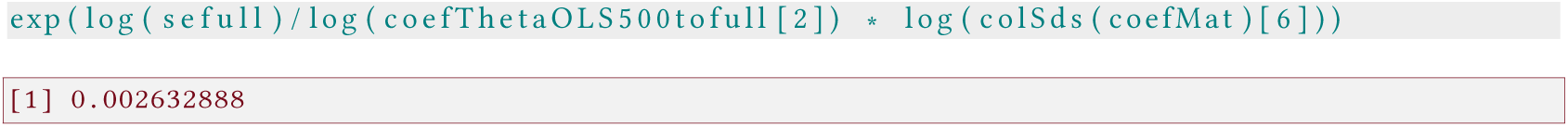

This is (obviously) dramatically better than the “bootstrap” estimate presented above, but still thrice the 0.0009 of Liang *et al.* (2016).

#### C.2.2 Increasing the size of the subsamples for the GLS

As another step, we can briefly check whether increasing the size of the subsample from 500 to 5000 for the GLS makes an appreciable difference.^11^

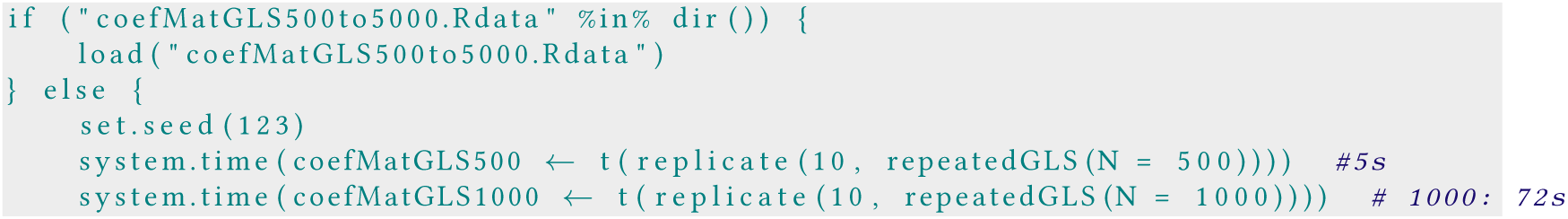

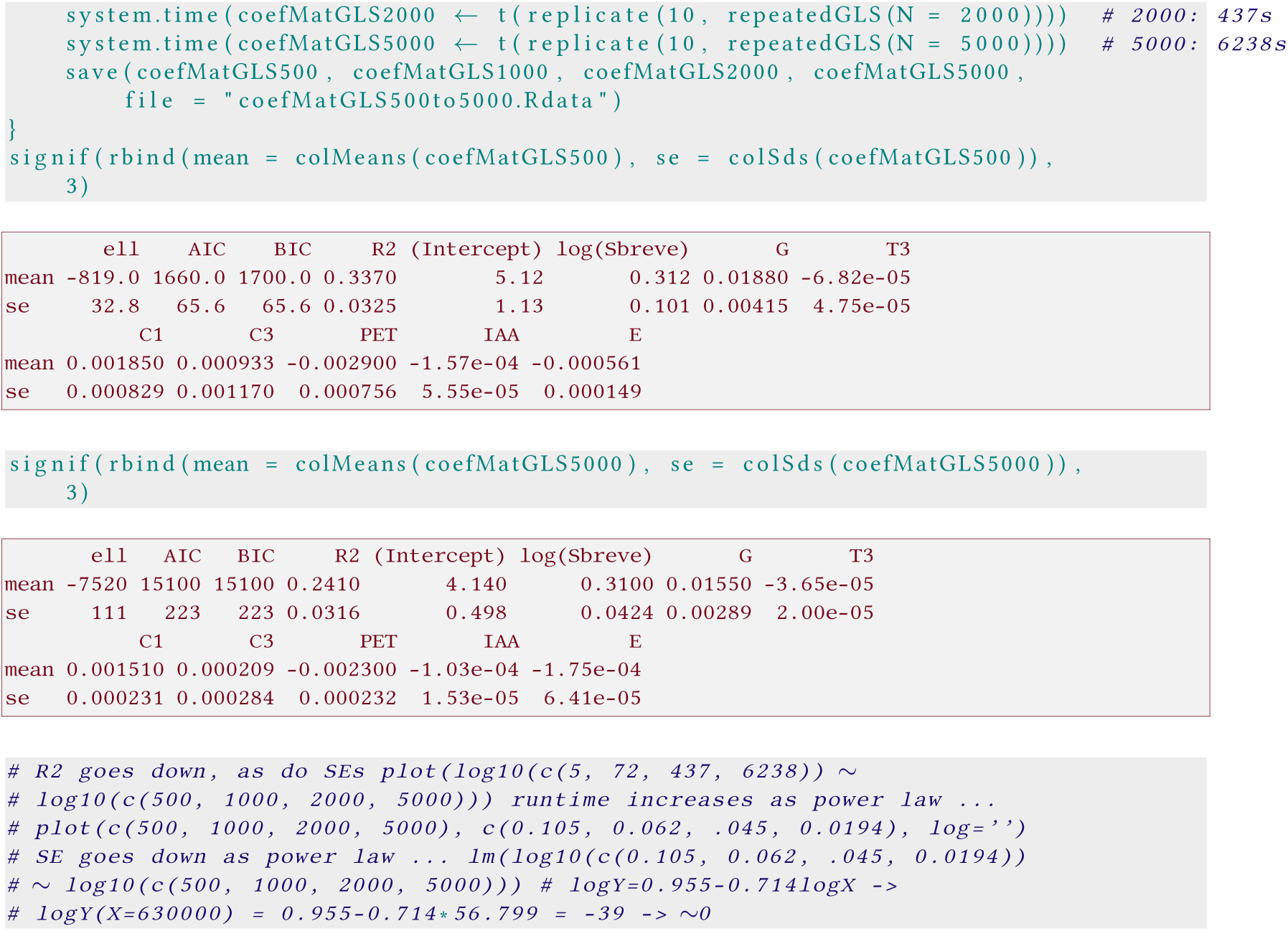

The estimate of *θ* and most other model parameters is similar for sample size 500 and 5000, while the intercept and the model’s R^2^ goes down.

What does that mean?

1. Using small subsamples of a large data set leads to dramatic increases in parameter uncertainty. Thus, the (corrected) “bootstrap” approach of Liang *et al. (2016)* is likely to be very pessimistic. Since they did an error in the computation of their standard errors, the uncertainty limits provided in the original paper cannot be trusted.
2. Using the full data set led to *very* different estimates for the model (comparing OLS for the resampled and the full data set, i.e. the grey and the blue curves above). By analogy, also the resampling-based estimates of the GLS are likely to be substantially biased.^12^
3. Correction for spatial autocorrelation leads to lower estimates for productivity, roughly 80% of the expected value of P along the tree species richness gradient (orange line).

Although we have introduced a way to estimate the uncertainty around the predicted relationship from the data by approximation in the previous section, we have not used it until the very end.

#### C.3 Stratified sampling of subsets

Each ecoregion has a certain share of the terrestrial surface, but within each ecoregion, not all area is forested. Crowther *et al.* (2015) provide estimates of the total land area as well as the number of trees for each ecoregion. We use the latter as means of stratification, i.e. draw plots proportional to the number of trees in that ecoregion.

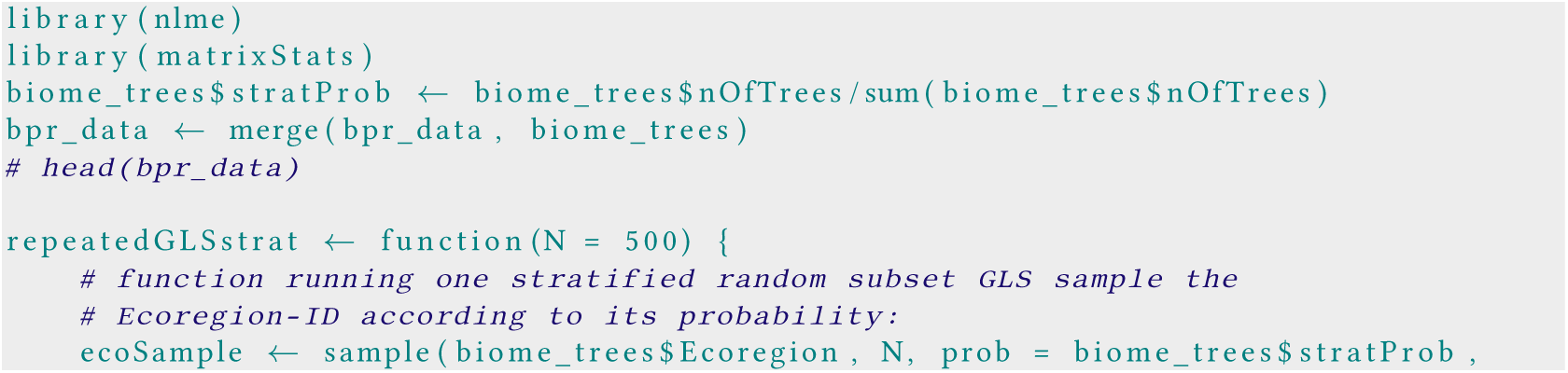

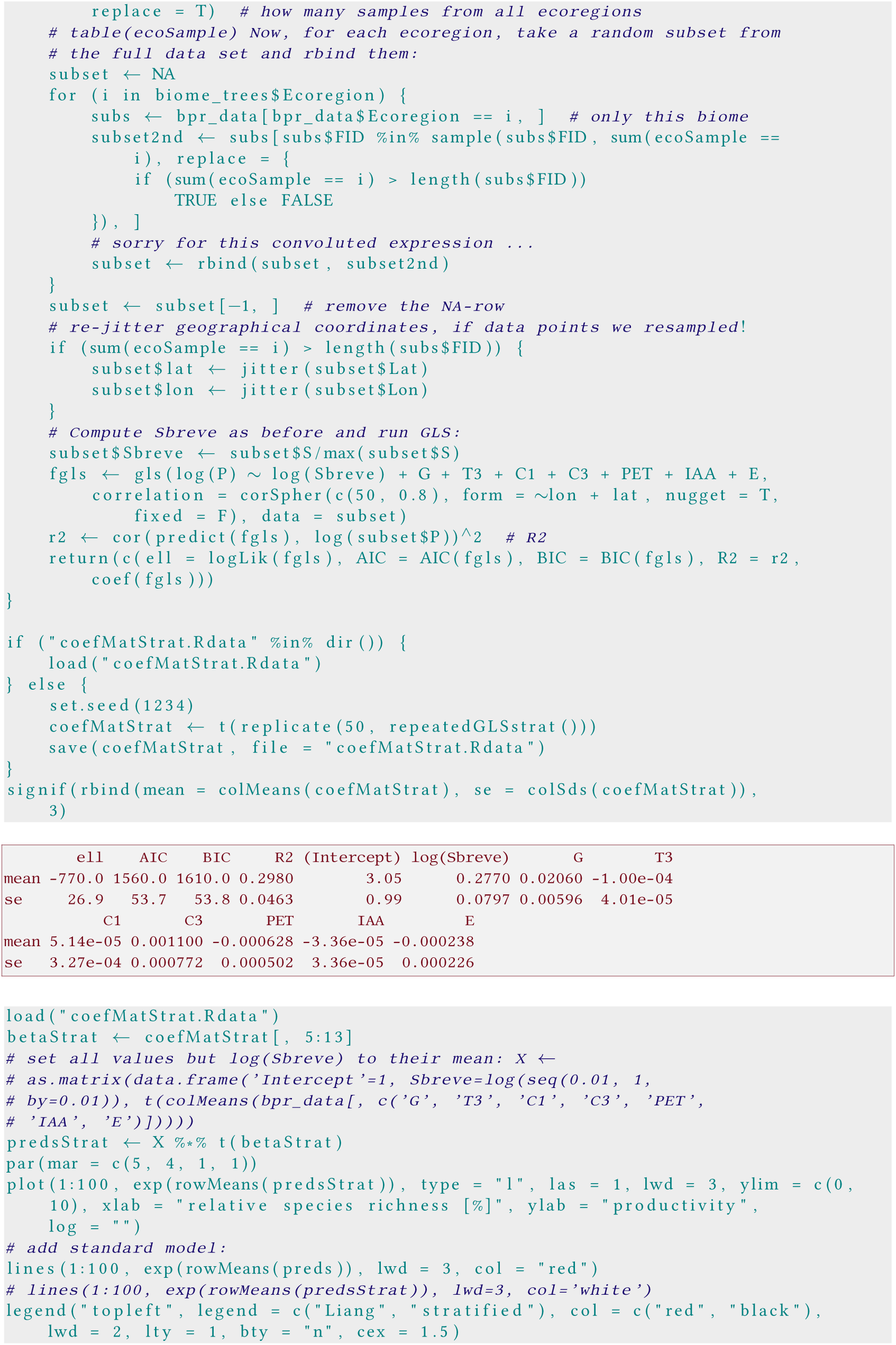

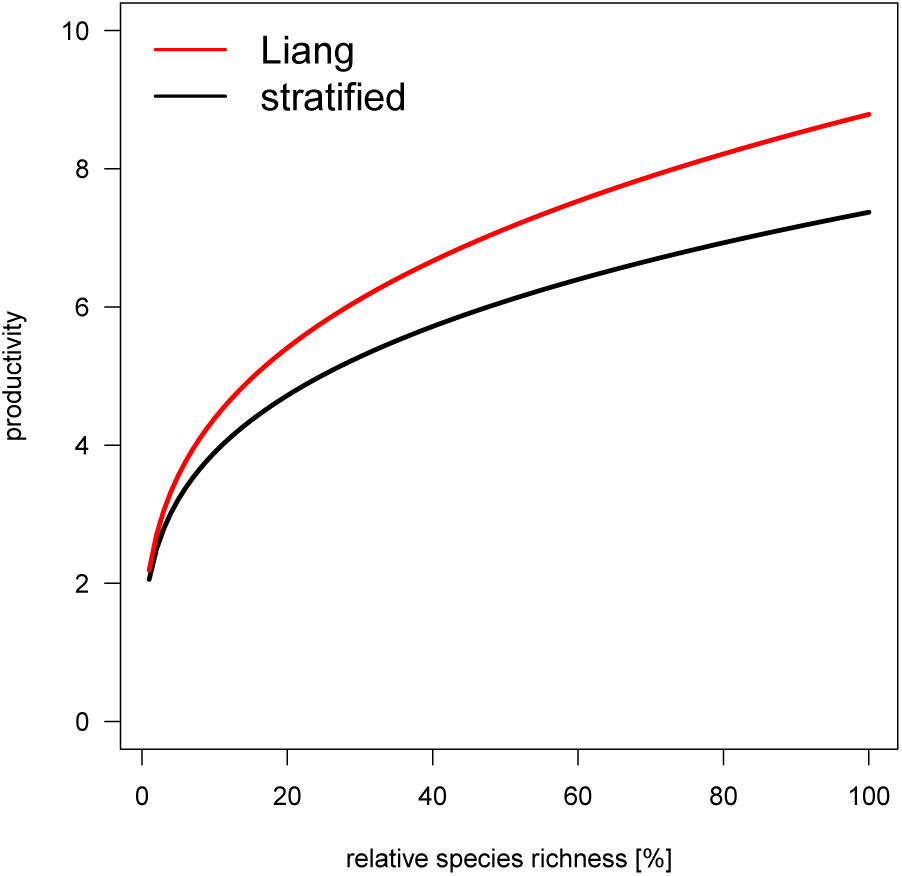

The effect is moderate, with slightly lower values than the original non-stratified approach. This result suggests that also with non-stratified sampling always some tropical plots with high species richness are drawn, making the original Š robust to unrepresentative sampling.

#### C.4 Compute distances between plots as Great Circle distances, rather than Euclidean (thereby accommodating the spherical nature of earth)

In the spatial model, the GLS, the distances between plots are computed from x-y-coordinates as Euclidean distance. On earth, the distance^13^ between two points cannot be computed as Euclidean distance, and instead requires the computation of the so-called Great Circle distance.^14^

We employ the “halversine” correlation structure provided on stackexchange to implement a correct distance measure within GLS.^15^ We define the model and run it repeatedly.

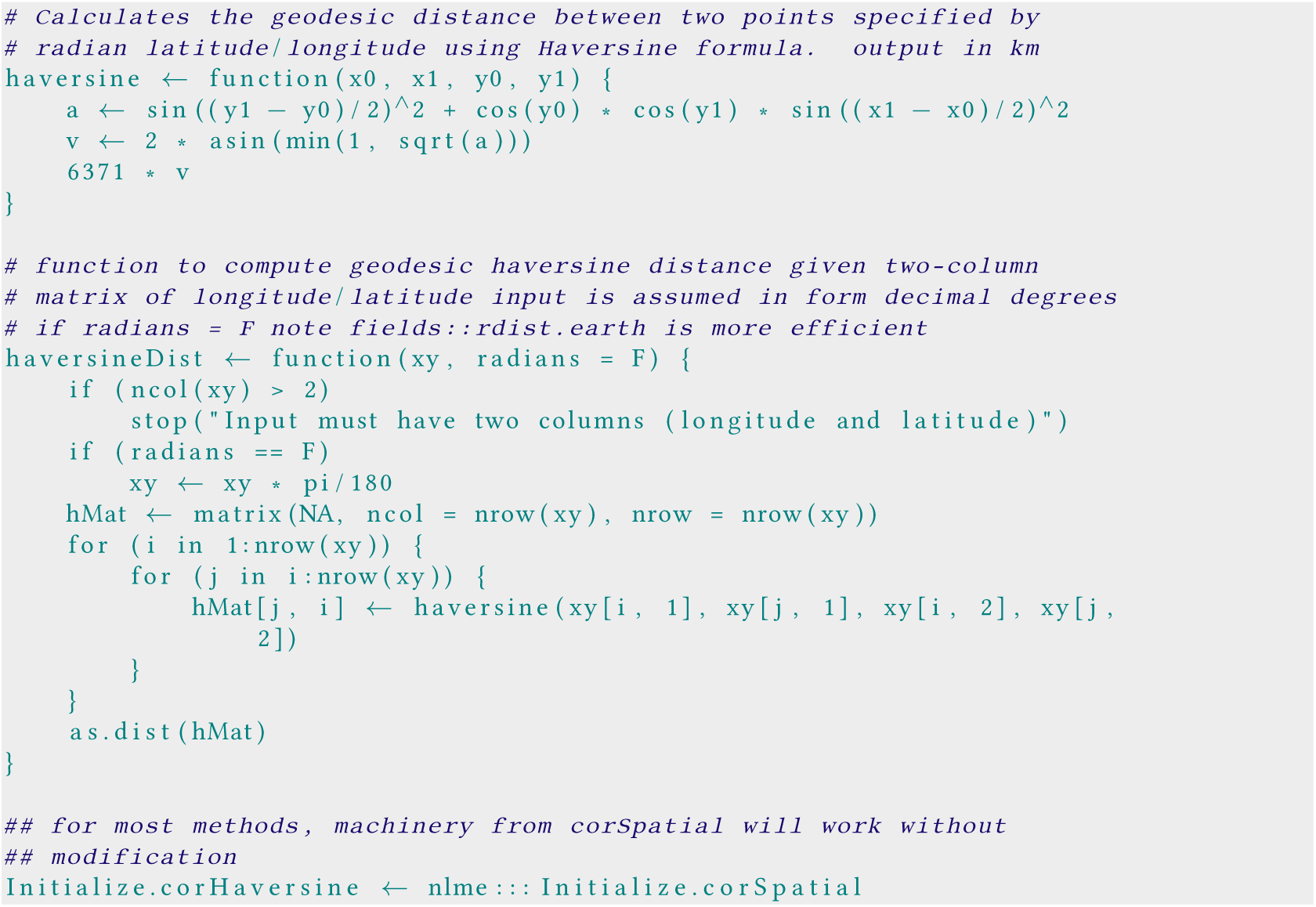

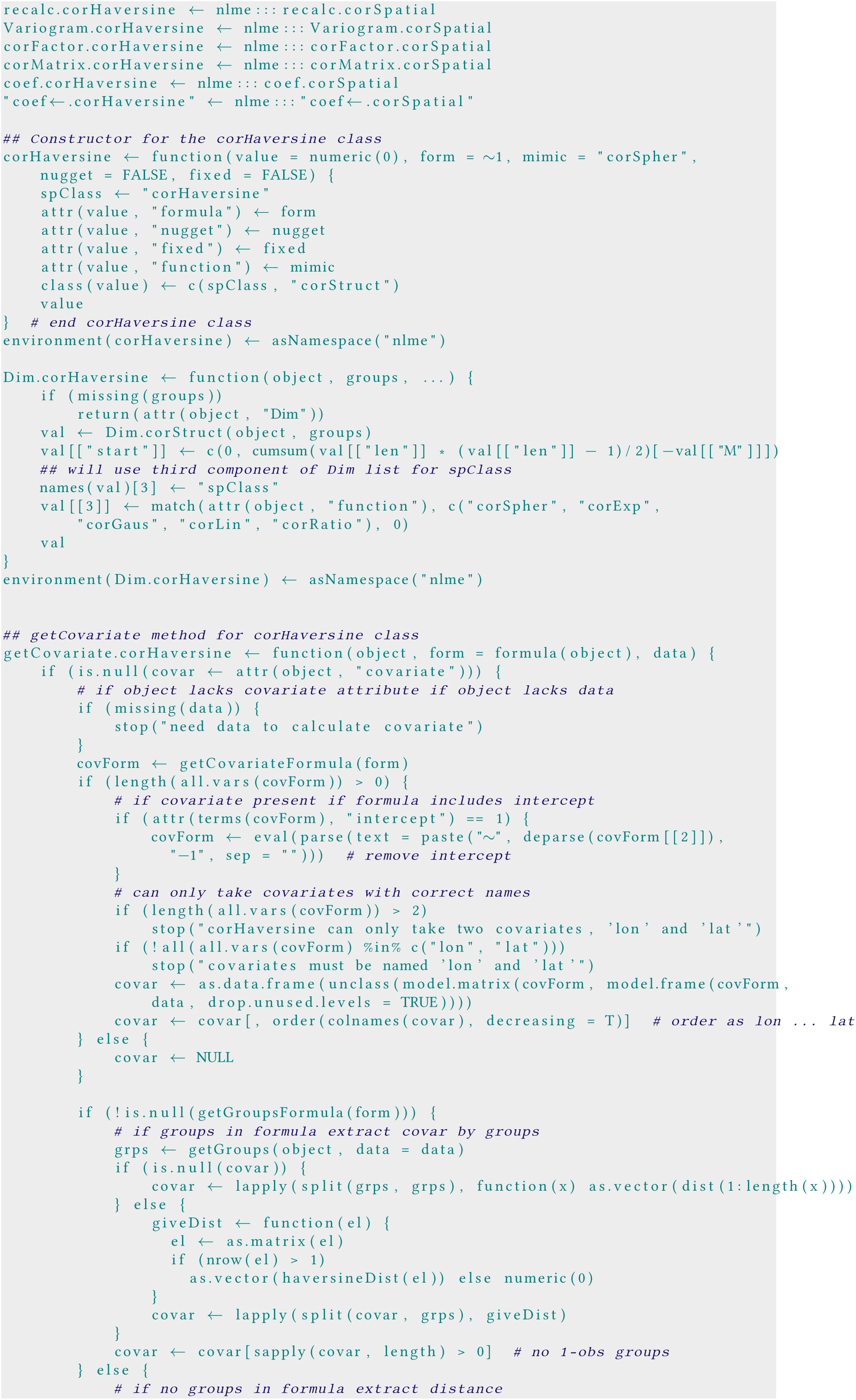

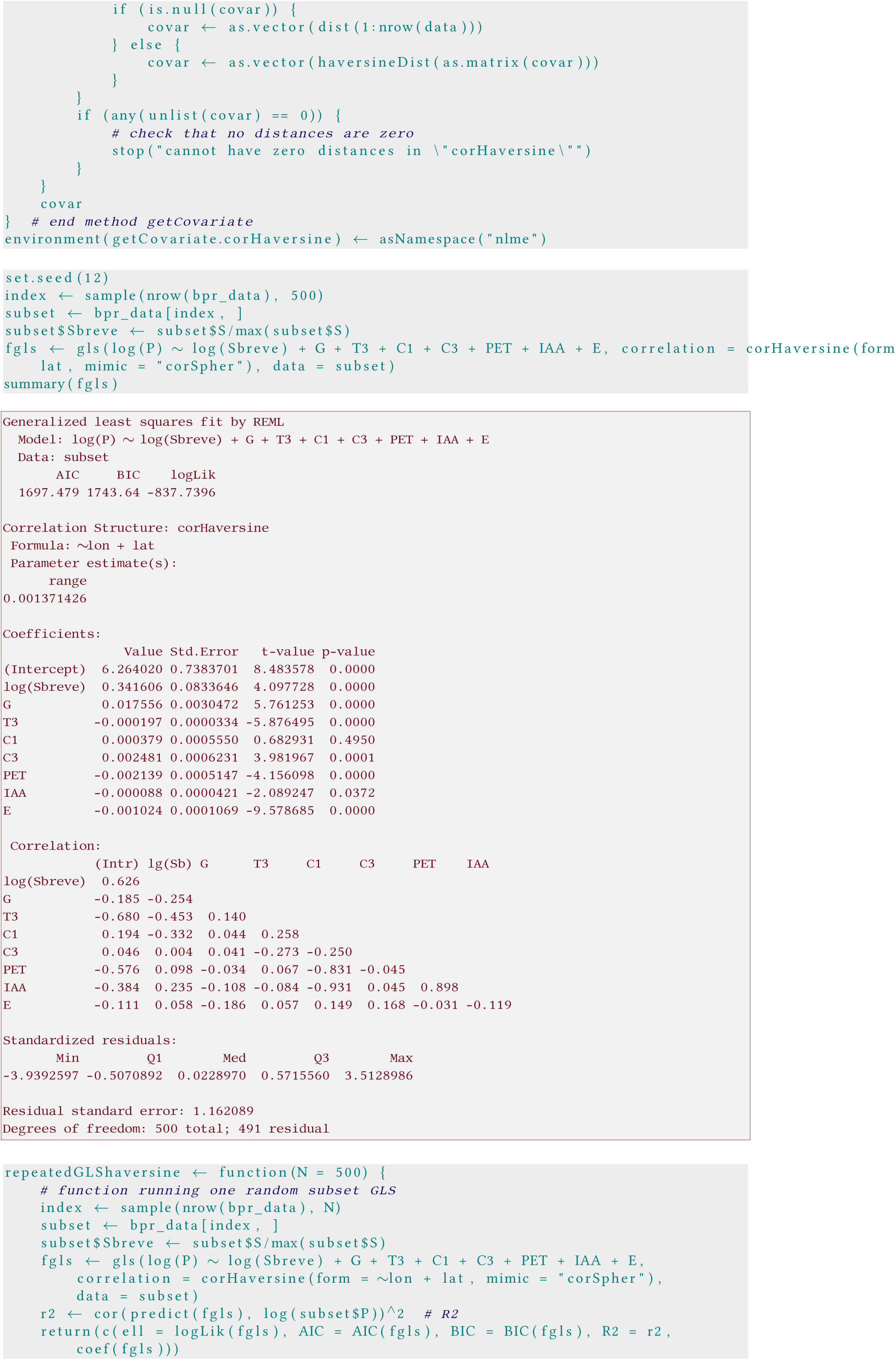

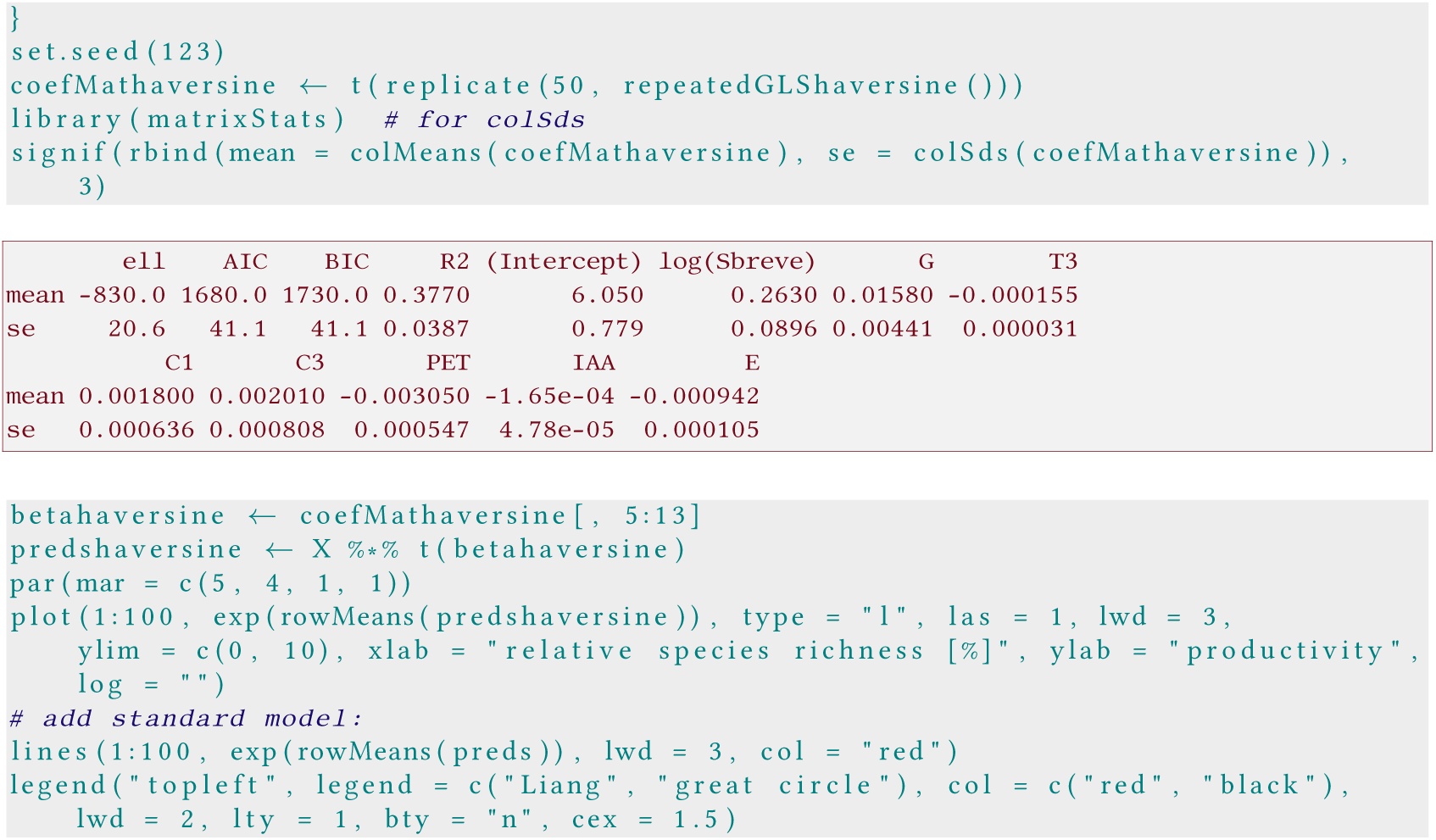

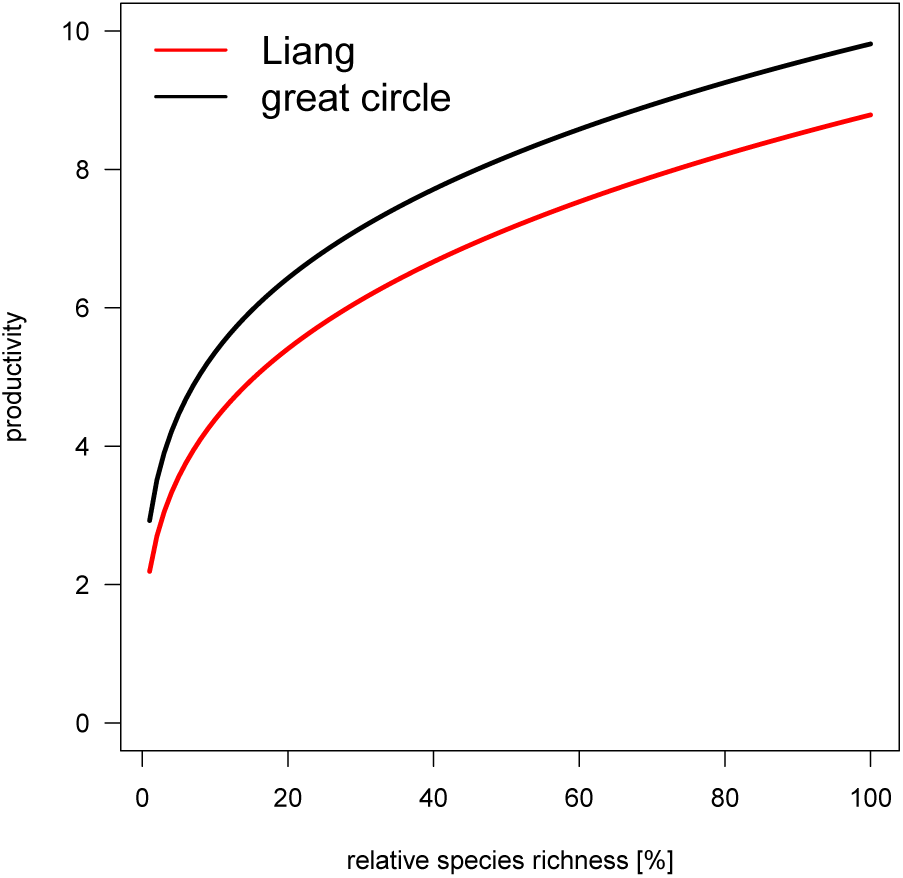

In this case, the correct representation of spatial distances increases overall productivity estimates a bit, but leaves the TSP-P-relationship unchanged.

#### C.5 Define species richness relative to what is regionally possible

The definition of Š in the subsets is relative to the largest value in the subset. When analysing the full data set, by the same logic, all tree species richness values are a fraction of the most diverse plot with 405 species. Thus, the reported relationship must be read as: “If we increase whatever number of species we currently have in a plot to 400, then we are moving on this curve.” Ecologically, this makes no sense. Whatever the reason why there are only a handful species in a plot of 1/10th of a hectare in temperate and boreal forests, it also prevents a “tropic richness” in these sites. Thus, one cannot meaningfully increase species richness outside the tropics to a tropical level. As a consequence, the depicted figure lacks ecological interpretability.

What the reader, and we suspect also some of the authors, probably interpret into the x-axis is “species richness relative to what would be possible at this site”. There are different ways to define “what is possible at this site”: (a) relative to what the maximum recorded value for this biome is, (b) relative to the richness of plots in the vicinity, (c) relative to the number of trees in the local/regional species pool, and possibly others. In the following, we use approach (b), relative to other plots in the region. The biome is, in our opinion, too large, while the third approach would require an analysis far beyond what current data allow us to do.

We define the “region” around a plot as being within a raster cell of 100 x 100 km^2^.

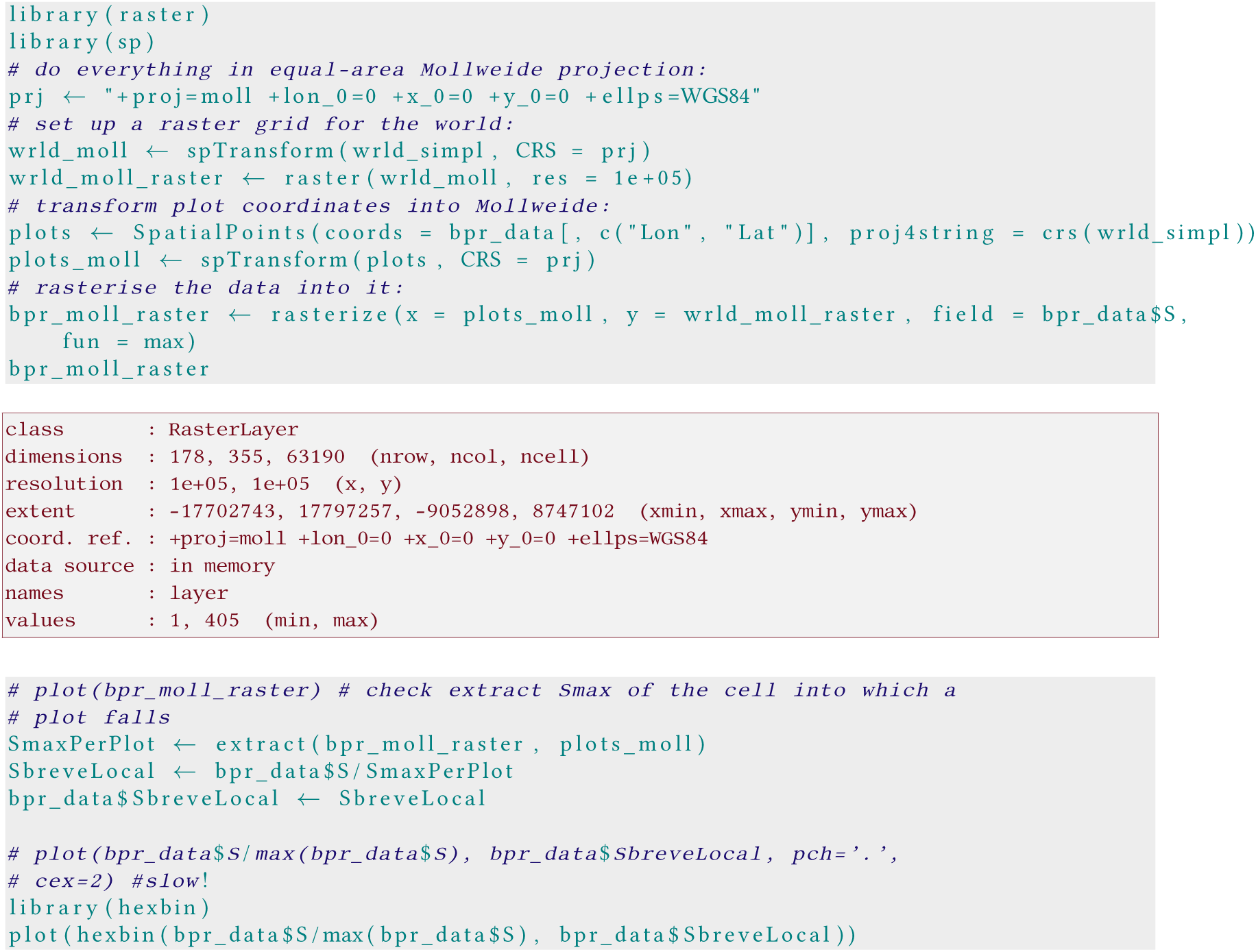

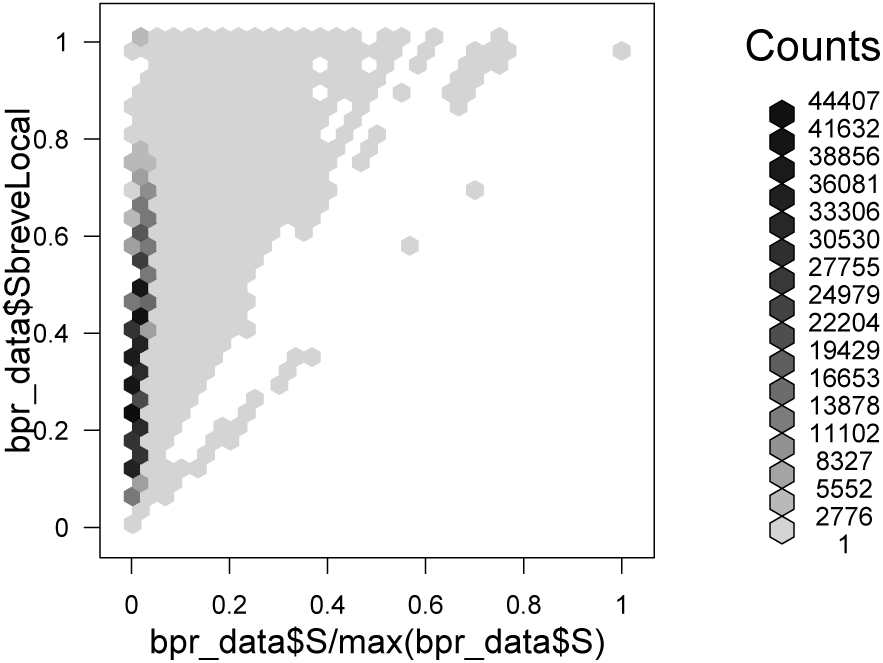

The local relative species richness (Slocal) shows substantially more variability for low-richness plots.

Now we can repeat the above analysis with a different relative tree species richness as response:

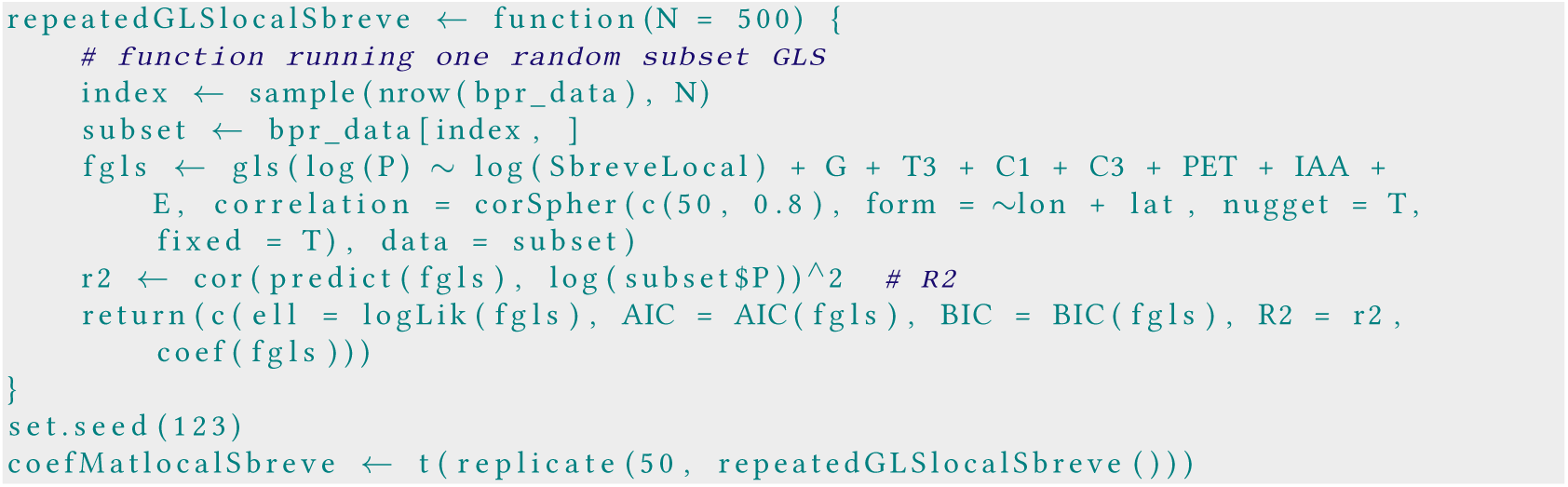

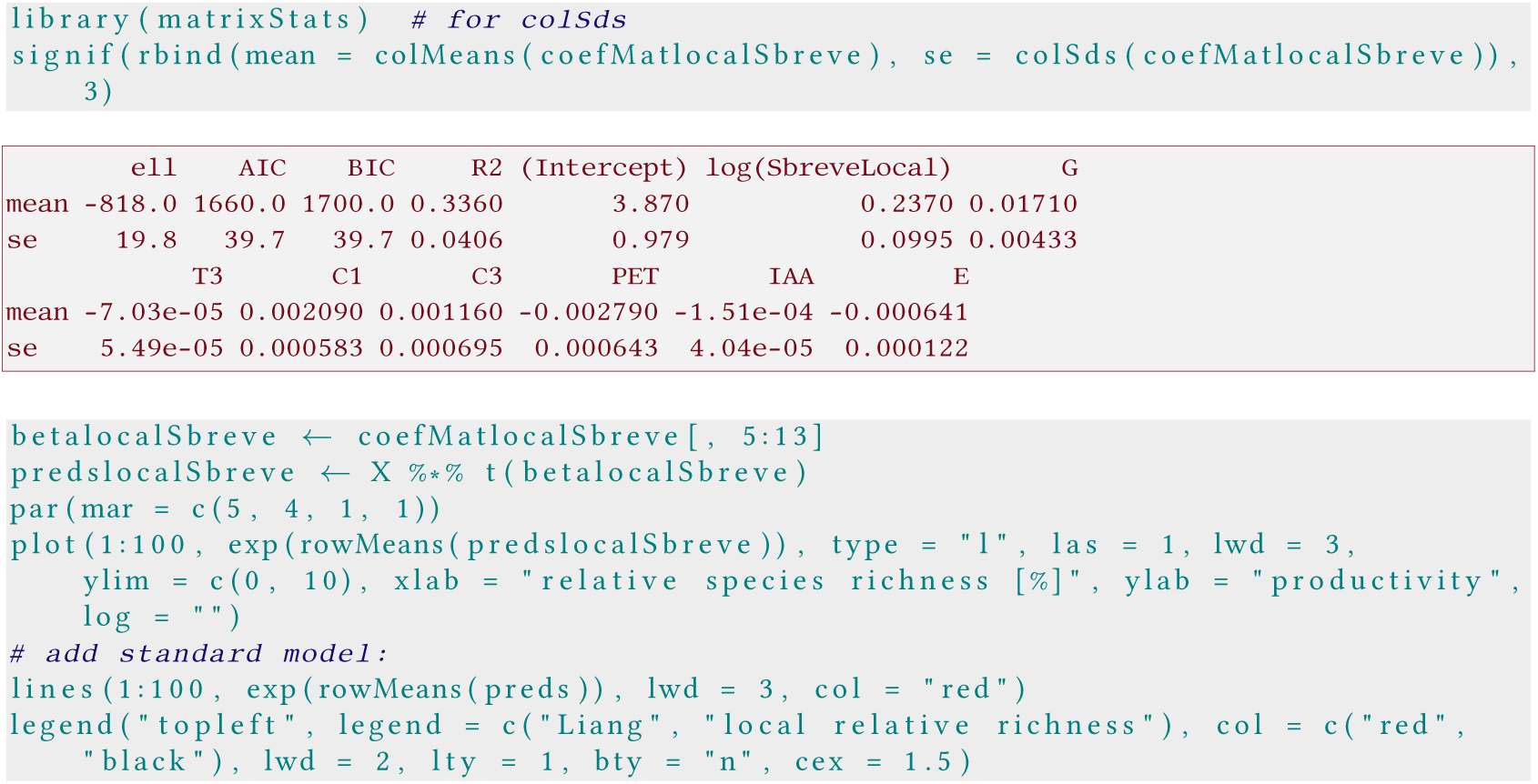

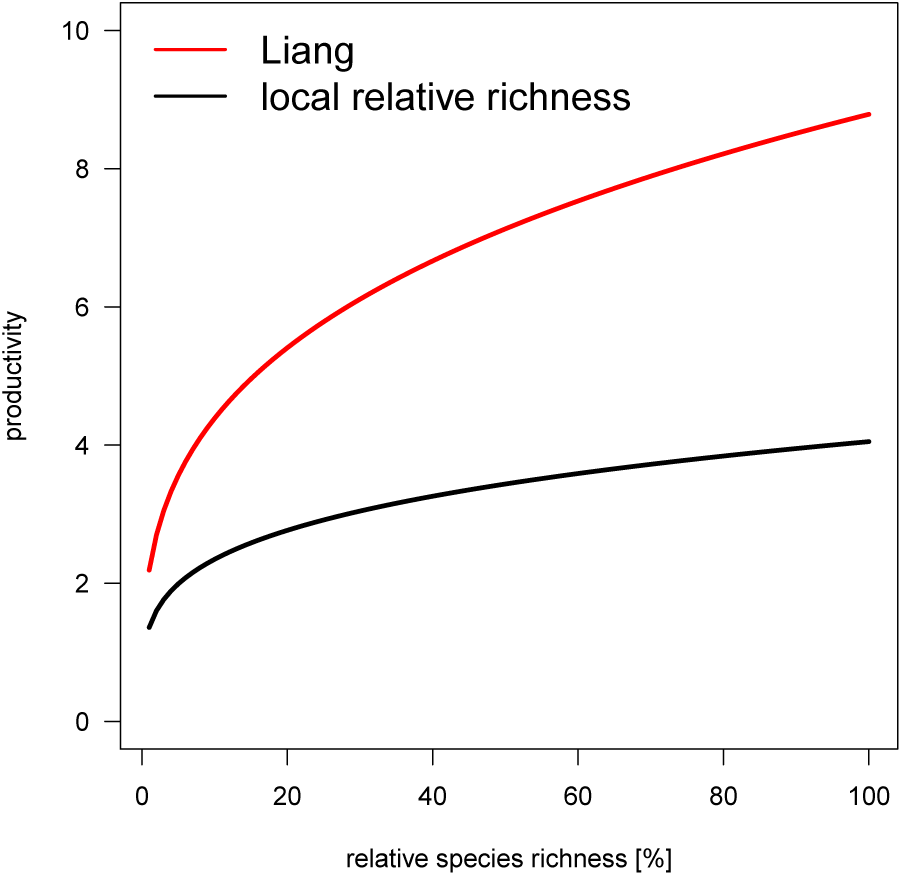

Representing species richness relative to what is possible in that region substantially reduces the effect of species richness on productivity, as well as the error margin of this relationship.

To repeat the message of this last plot: When moving from a plot that only comprises, say, 10% of the potential local tree species to one that contains all 100%, productivity increases from around 2.2 to 4.5 m^3^ha^−1^ y^−1^ (much less than the increase from 2.5 to 8 m^3^ha^−1^ y^−1^ reported by Liang et al. 2016).

#### C.6 Use relative, rather than absolute, productivity

It appears a bit odd to scale species richness relativ to what is locally possible, but not also productivity. Tropical forests can be expected to be far more productive than boreal ones, but the effect of relative species richness on *relative* productivity may be similar.

To explore this idea, we follow the same approach as with local relative species richness and compute local relative productivity, P_local_. This we then analyse with the local relative species richness (Šlocal).

Again we define the “region” around a plot as being within a raster cell of 100 x 100 km^2^.

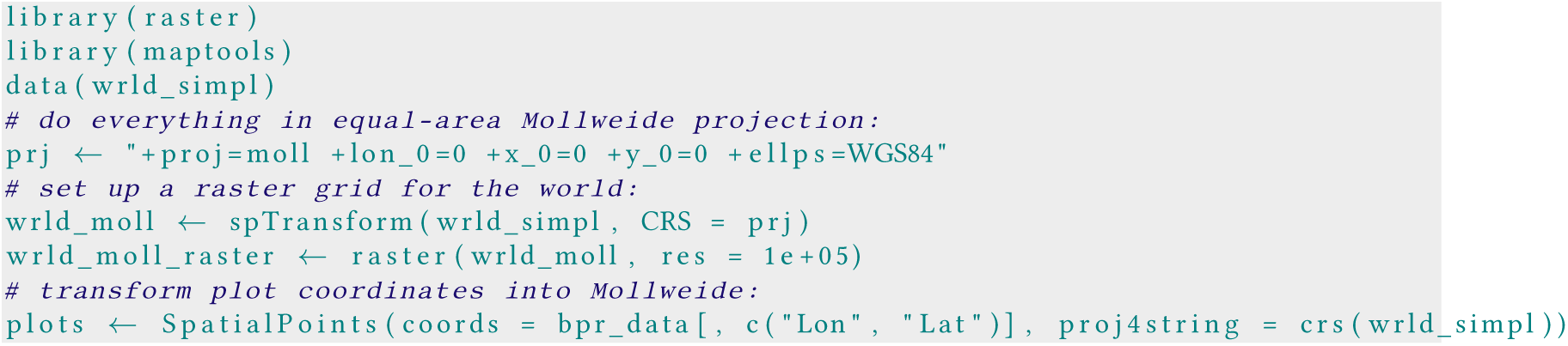

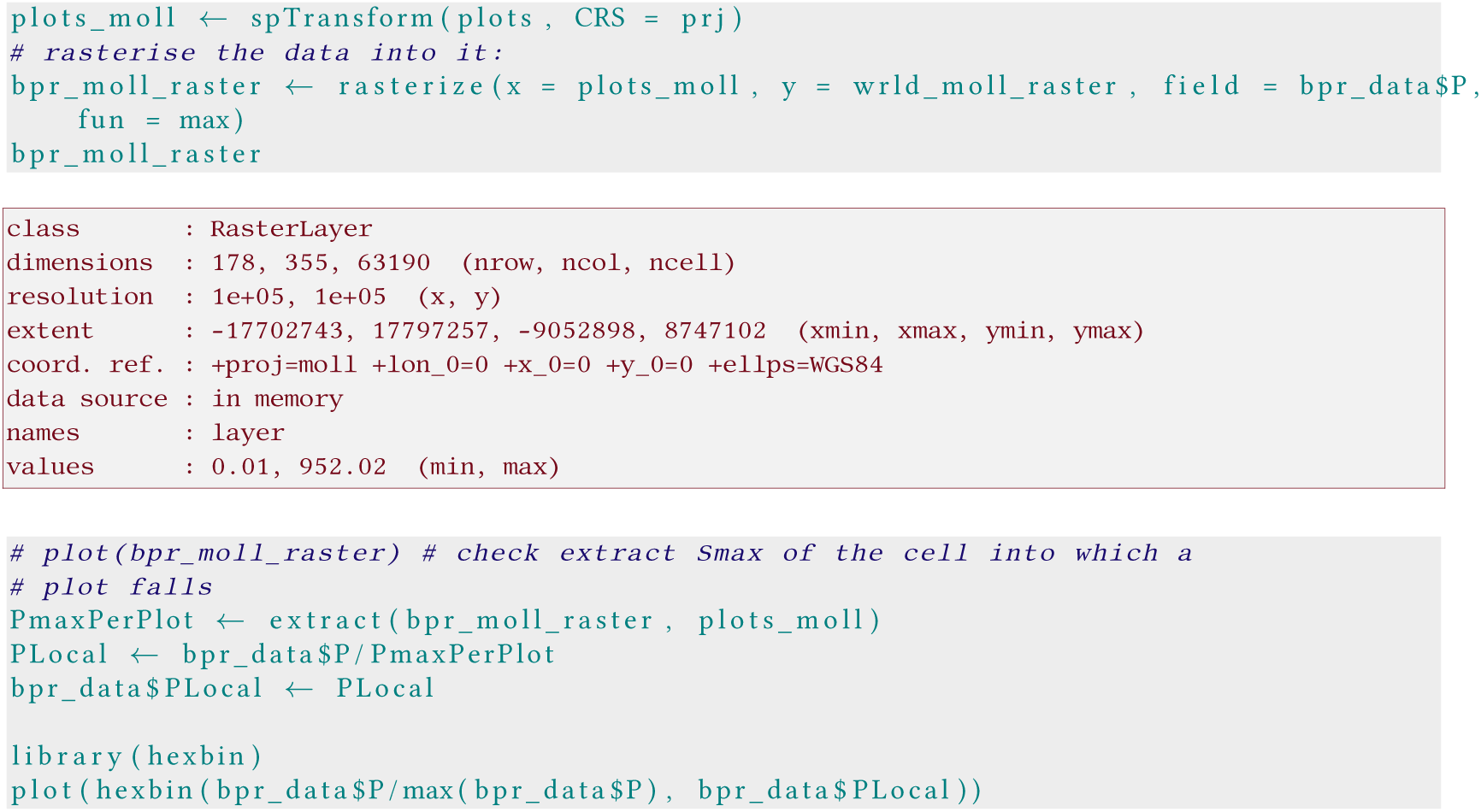

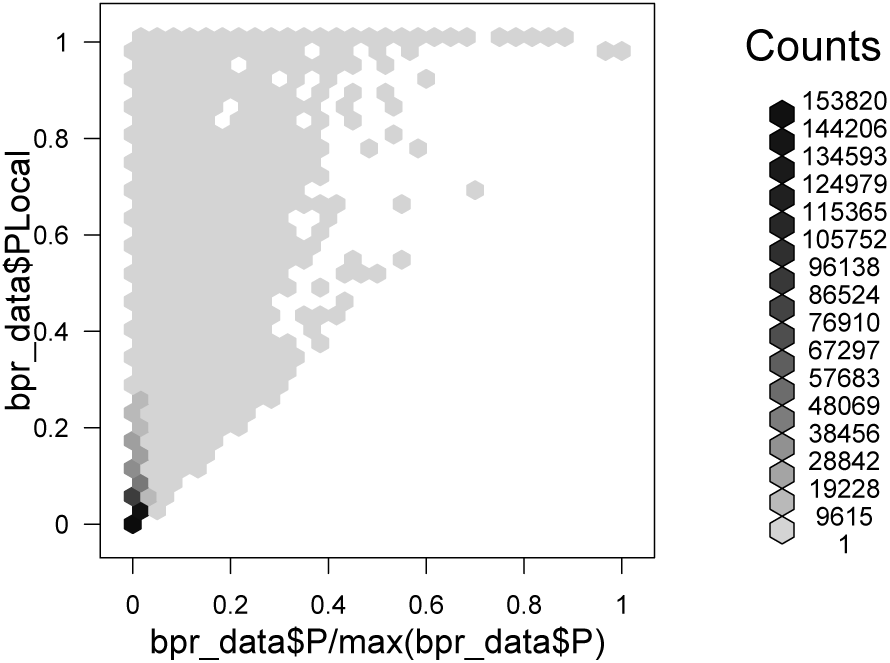

Now we can repeat the above analysis with a different relative (local) productivity as response:

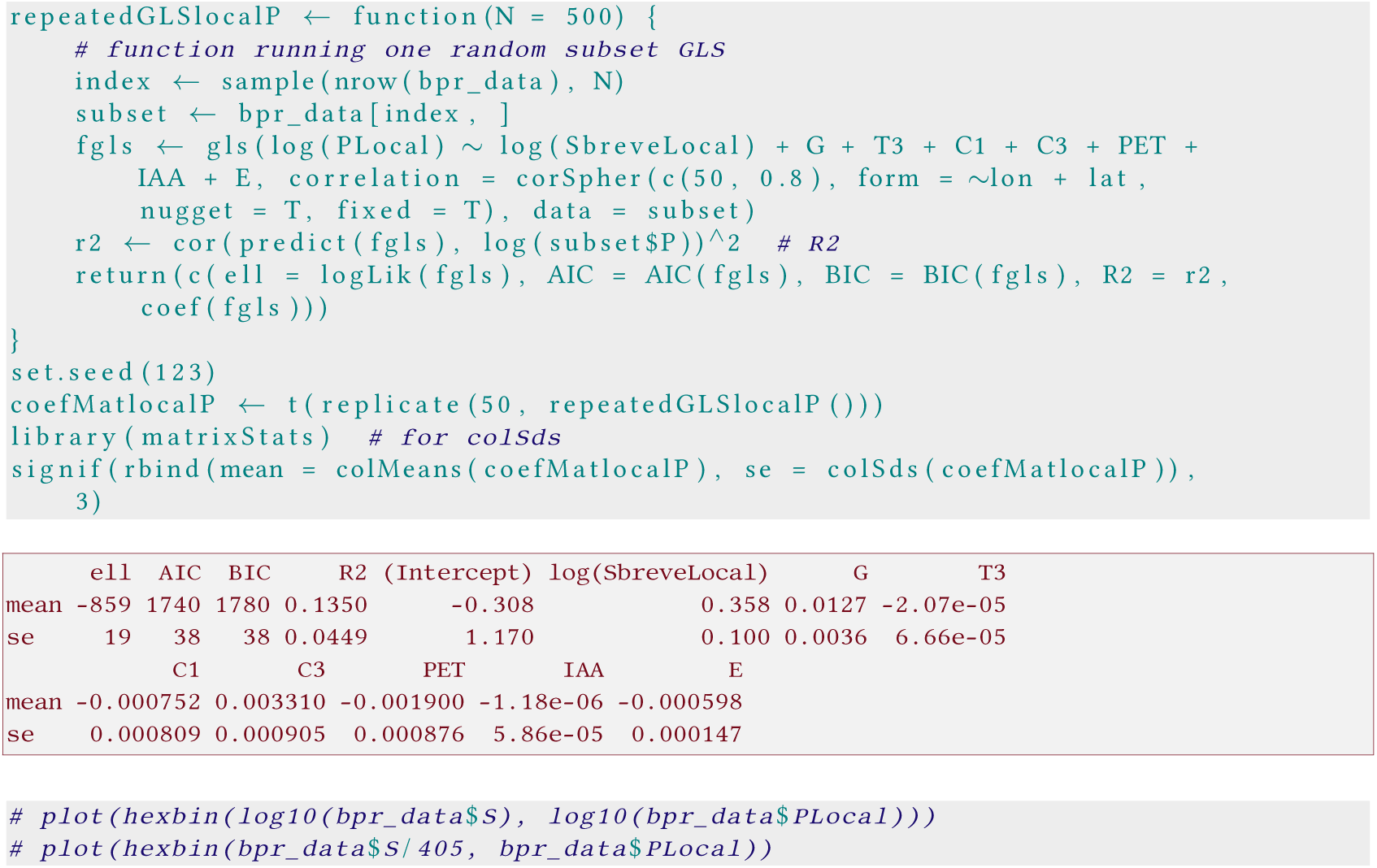

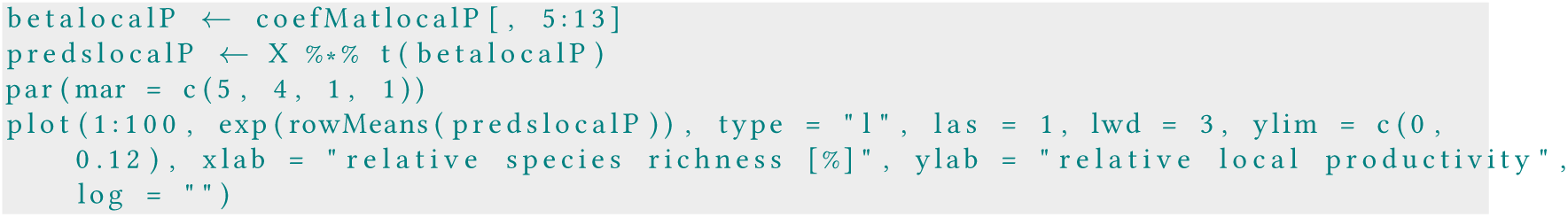

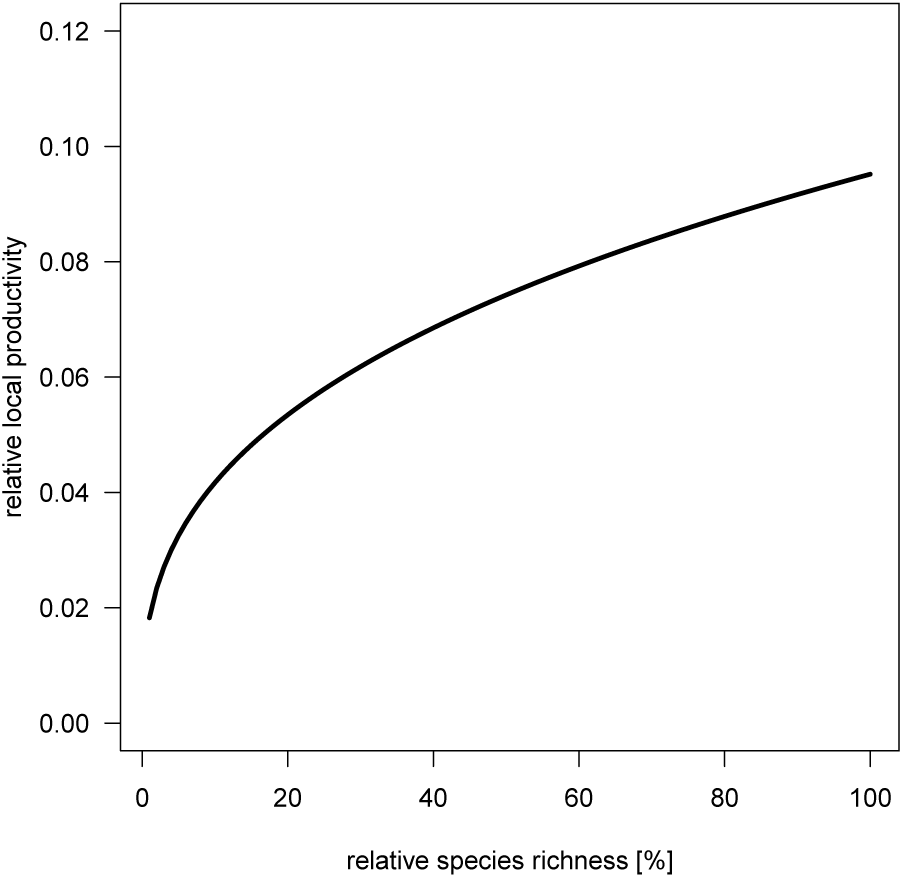

This does not look like an improvement, as the model fit is poorer (R^2^ of 0.135 compared to 0.34). Interestingly, however, (a) the effect of species richness is stronger (larger value of *θ*); and (b) even at maximal species richness only 10% of maximal productivity is reached.

Note that we used the *local* relative species richness ((Š)local); the model with the original Š is similarly bad, and with a lower value for *θ* (around 0.213). While this may actually be interesting in itself, we do not pursue this line of thought further here.

### D A new analysis

In summary, our explorations above have shown substantial effects of (in that order)

1. computing Š relative to maximal local species richness;
2. stratifying the data proportional to an ecoregion’s share of global forest cover;
3. computing spatial distances on a sphere;
4. stratification of samples by ecoregion forest cover;
5. correcting for spatial autocorrelation;
6. correcting standard errors for subsampling.

The relative scaling of productivity did not improve the global model and is not further considered here.

It is quite possible that these alterations of the original model interact, i.e. that the substantial bias we have seen introduced by stratification may be caused by the way Š is computed, and would be removed with a locally computed Š _local_. Exploring all these option combinations is beyond the scope of this re-analysis.

So, in a nutshell, the *new global model* uses a locally-scaled species richness as predictor, computes spatial distances on a sphere when correcting for spatial autocorrelation in the data and stratifies sampling by each ecoregion’s forest cover.

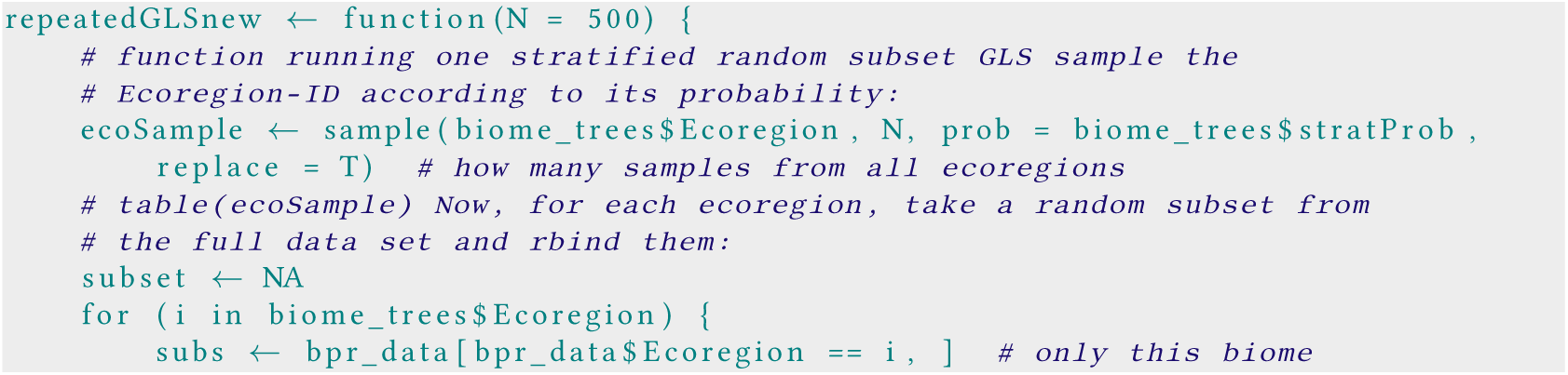

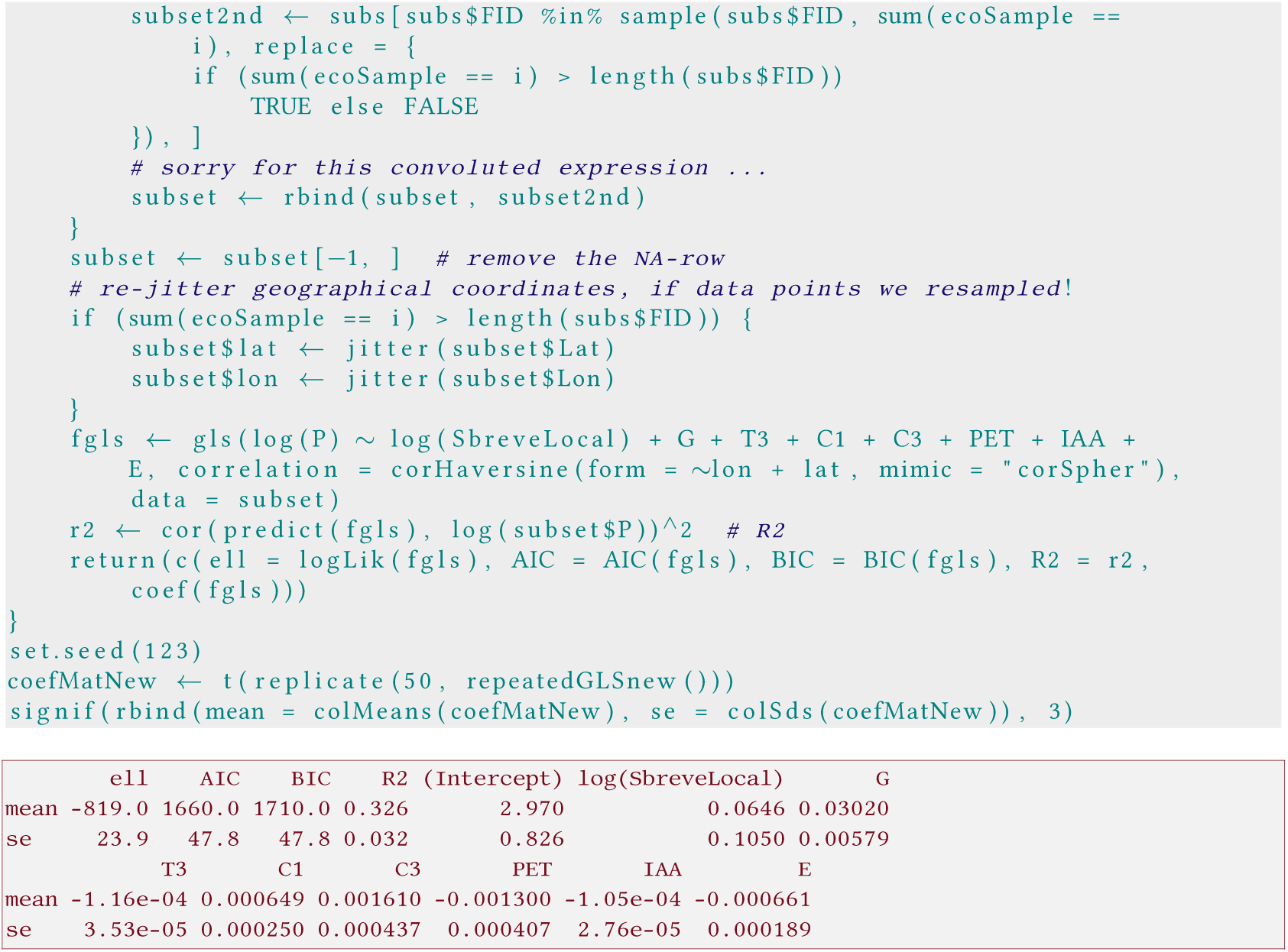

Here we add the correction for subsampling, specifically computing SDs of the predictions from the subsamples and correcting them to the full data. We first do that for the fits provided by the re-analysis in the original form.

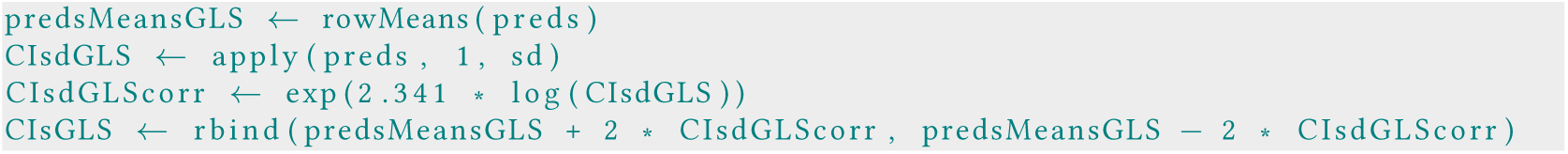

Now we plot the new model, and as comparison the original.

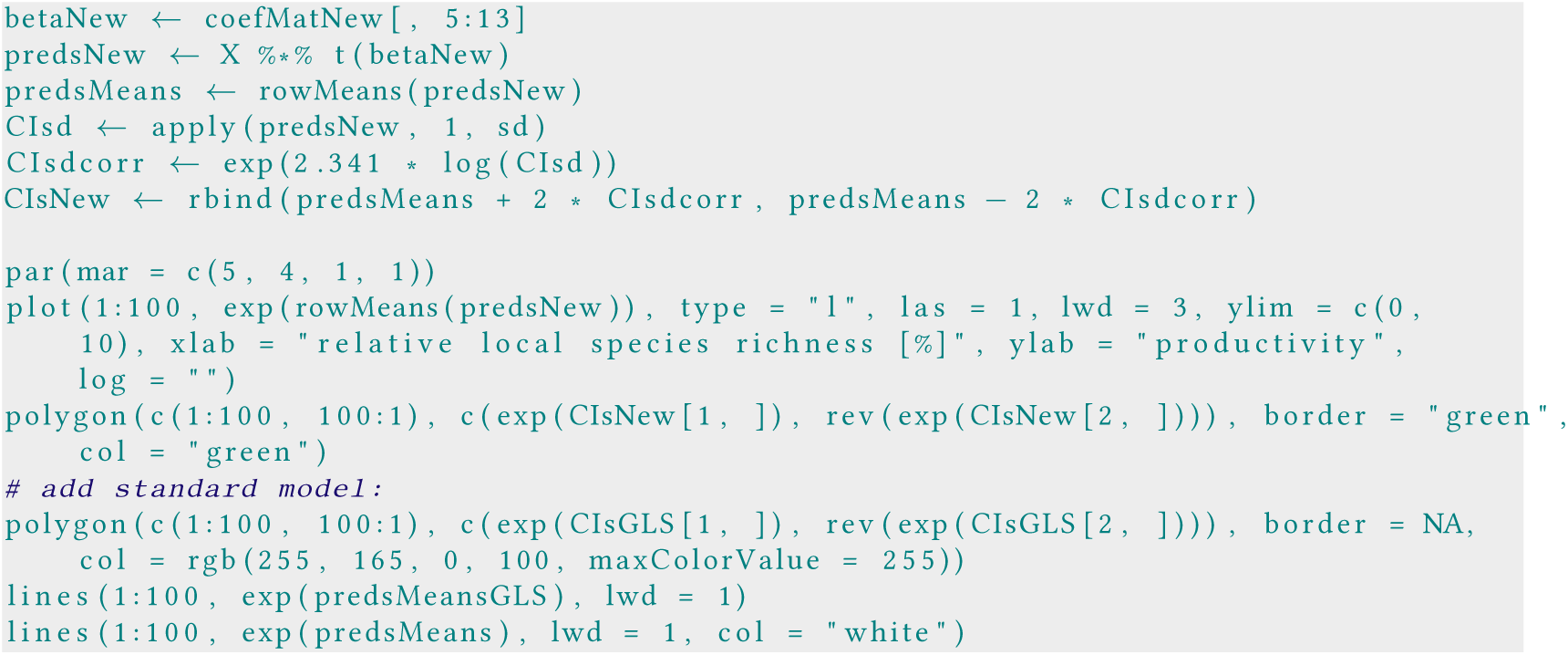

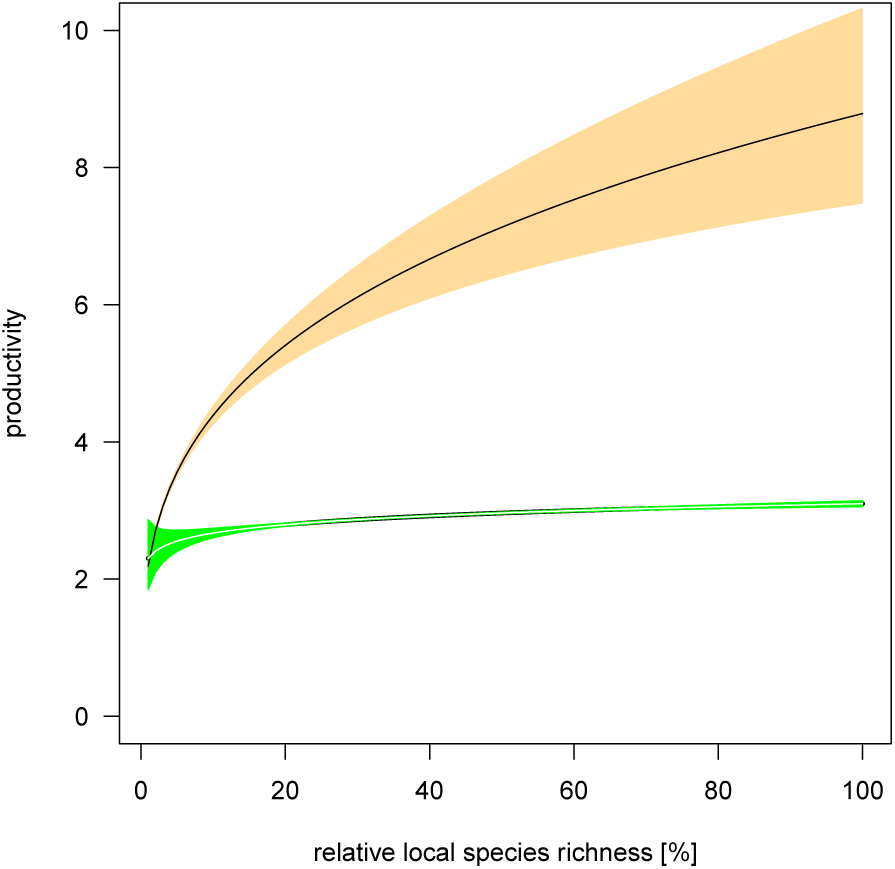

As we can clearly see, the pattern observed by Liang *et al.* (2016) does not hold up to scrutiny when correcting the mistakes of their analysis. The overall effect of increasing species richness to the maximum possible at that site is negligible and levels off at less than 10%.^16^

The absolute levels of productivity presented in the above figure are conditional on the values choosen for environmental covariates and are thus not interpretable “globally”. To do so would require a marginal, rather than a conditional approach. As already the conditional effect at the mean value of the covariates is practically absent, so would probably be the marginal. ^17^Although that was one of our initial questions, it is not worth pursuing under these conditions.

### E Comparison with ecoregion-specific fits

The lack of species richness-effect begs the question whether *within* ecoregions such a species richness-effect is present. It could well be that the variation in species-richness effects across ecoregions leads to a dilution of this signal at the global scale and the overall lack of effect in the full analysis.

We thus recompute with the same approach the fits to each ecoregion separately.

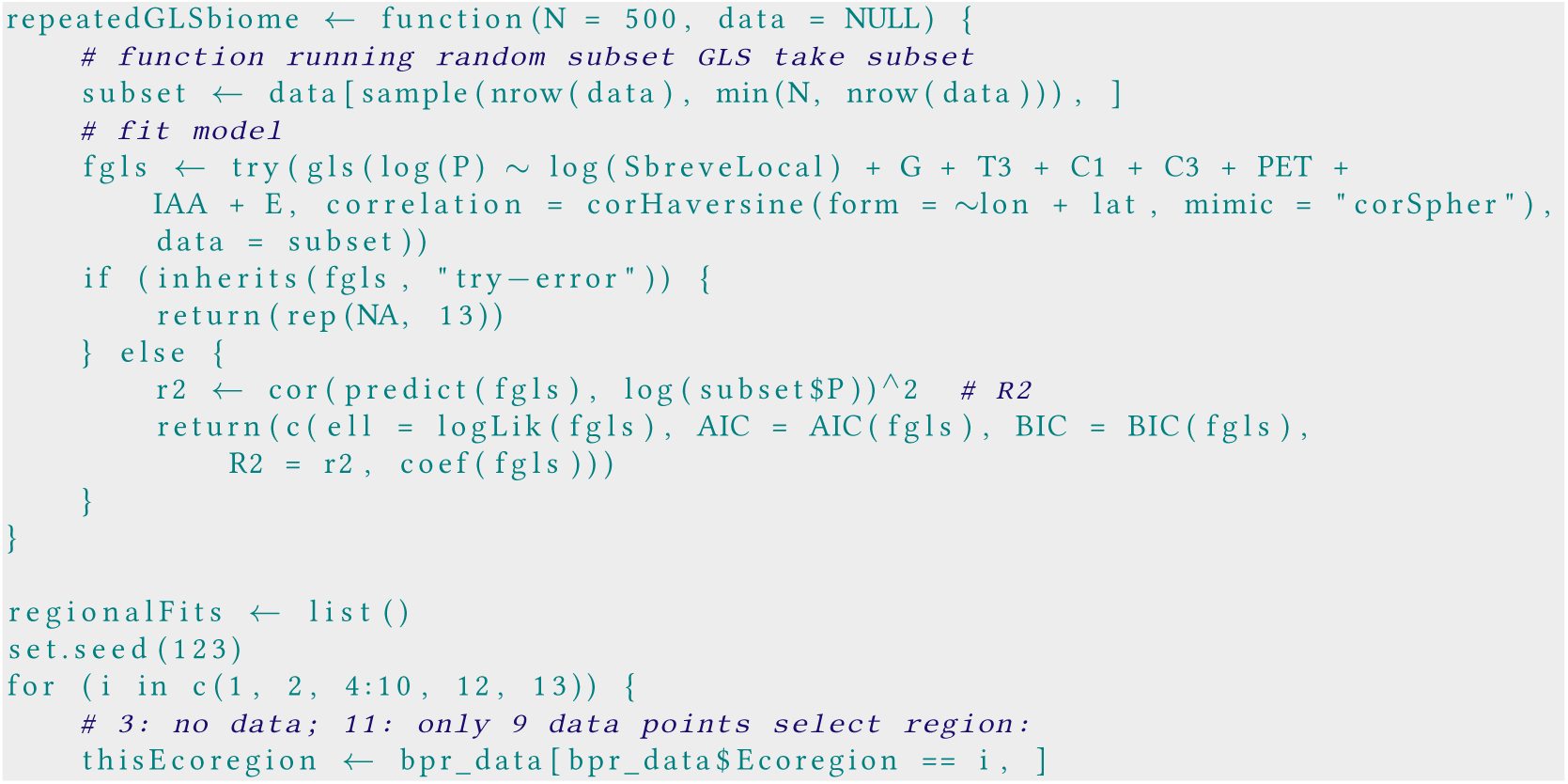

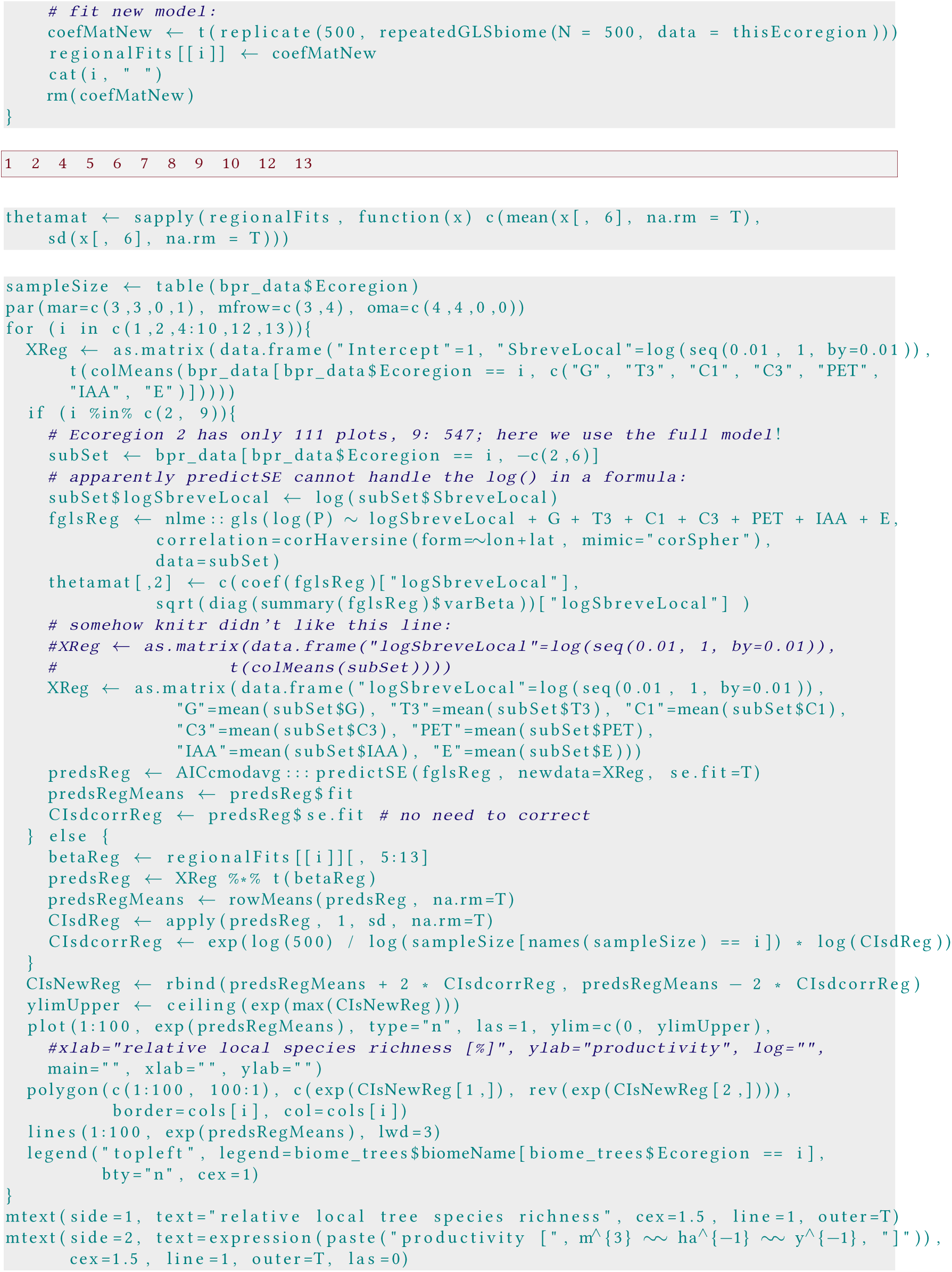

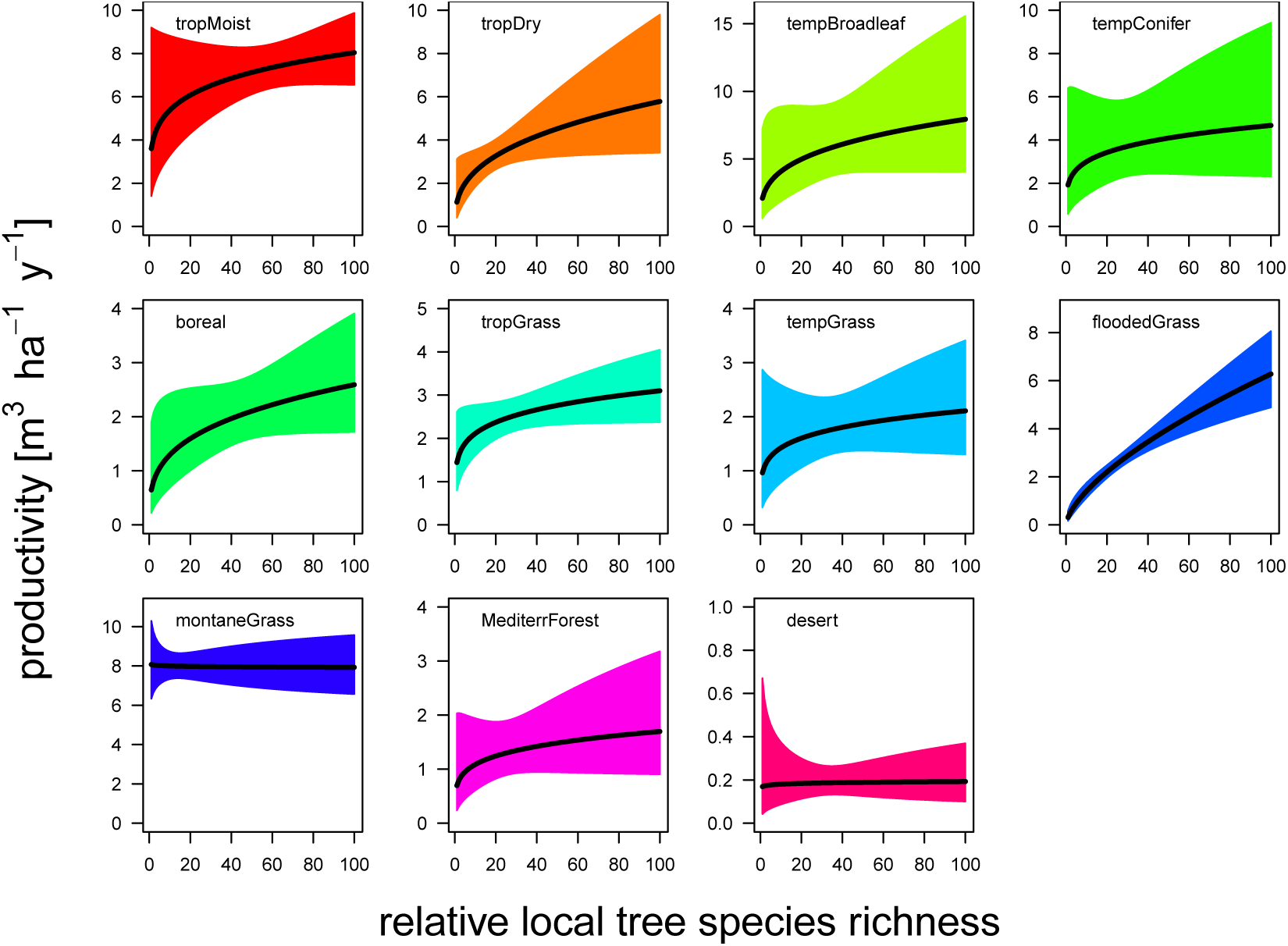

As a summary, we can plot the estimates for each ecoregion.

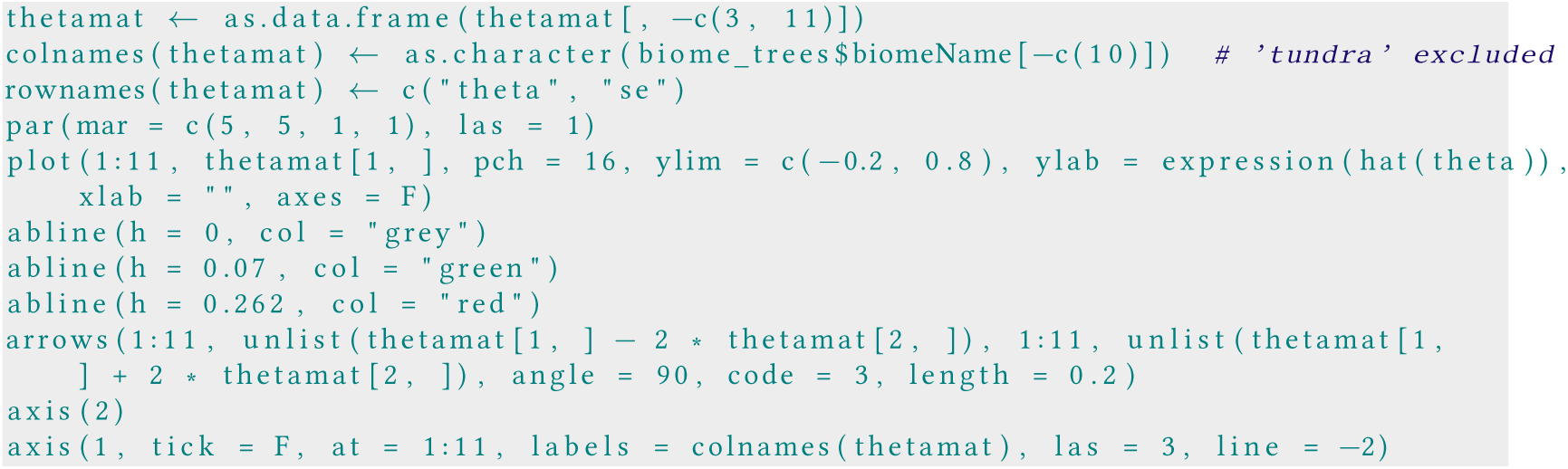

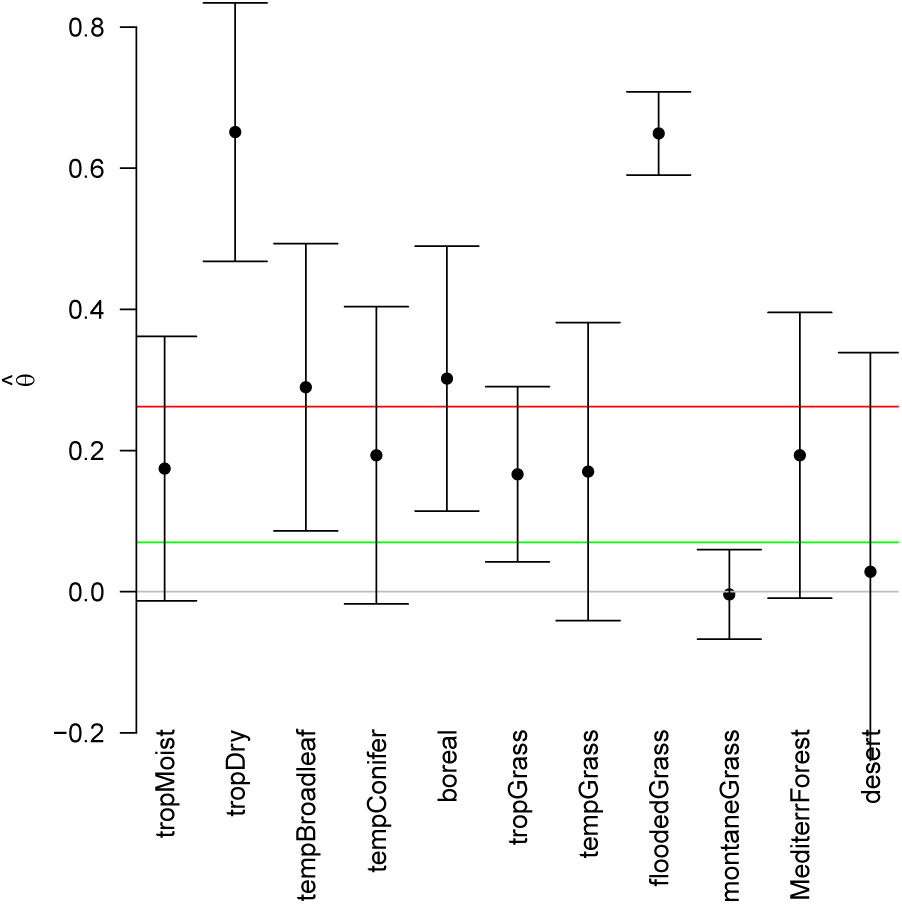

In six out of 11 cases (tropical moist rainforest, temperate coniferous forest, temperate grassland, mountain grassland, Mediterranean forest and desert) the estimates are not significant. All others point at positive effects of relative local species richness of rather different strength (very strong in tropical dry rainforests and flooded grasslands). The original global model of Liang *et al.* (2016) (in red) shows a substantially stronger effect than our re-analysis (green).

#### E.1 Analysis with actual species richness as predictor

One of the interpretational problems of the above analyses, and indeed the key neat idea of the original paper, is that a change from, say, 10 to 30% on the x-axis means completely different things in different ecoregions. In the tropics, this may be a change in species richness from 30 to 90 species, while in the boreal system from “less than one” to “just under two”.

As a final step, we re-analyse the data with the same model as in the previous section, just with log(S) as predictor, rather than a somehow corrected species richness.

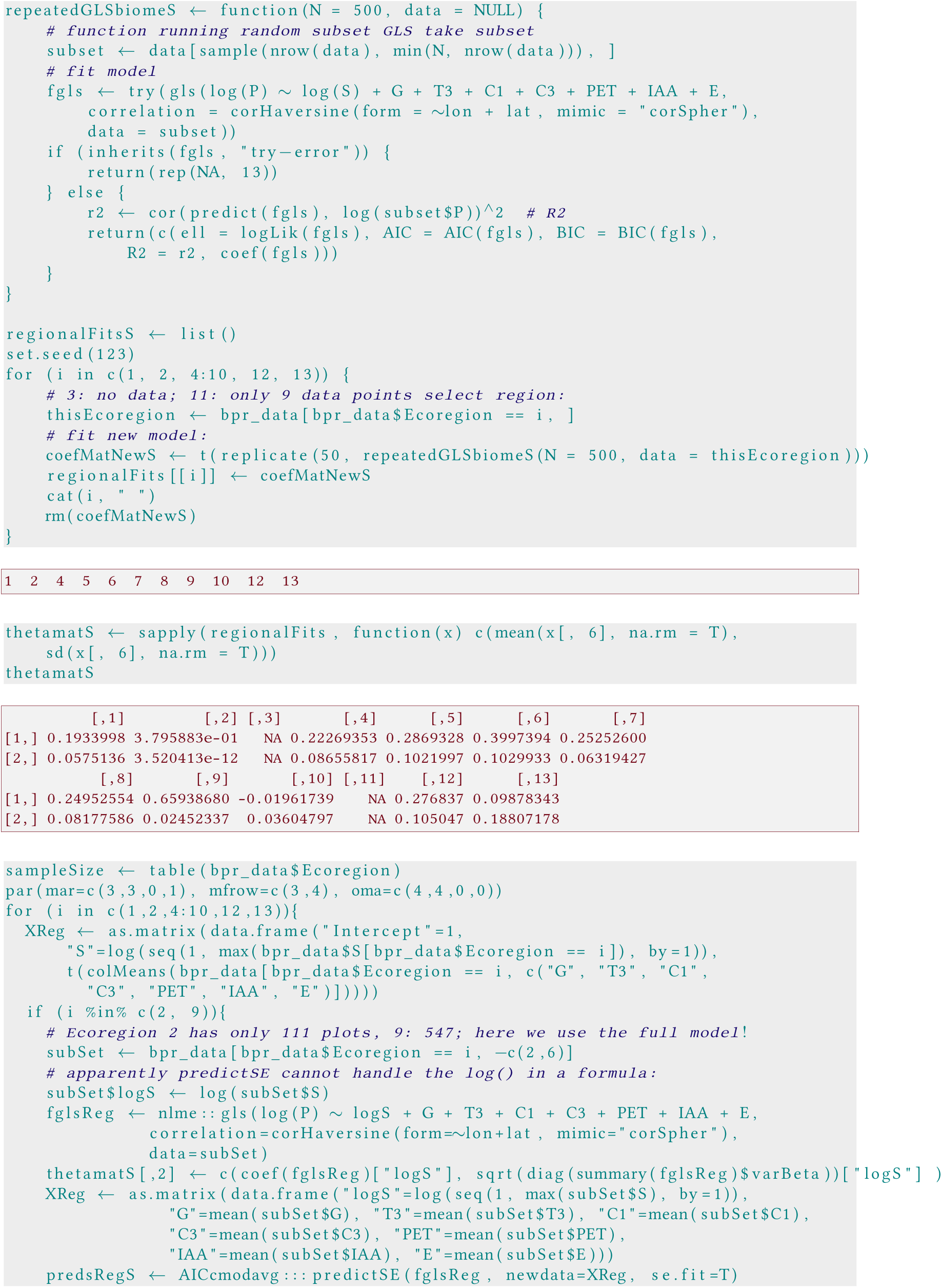

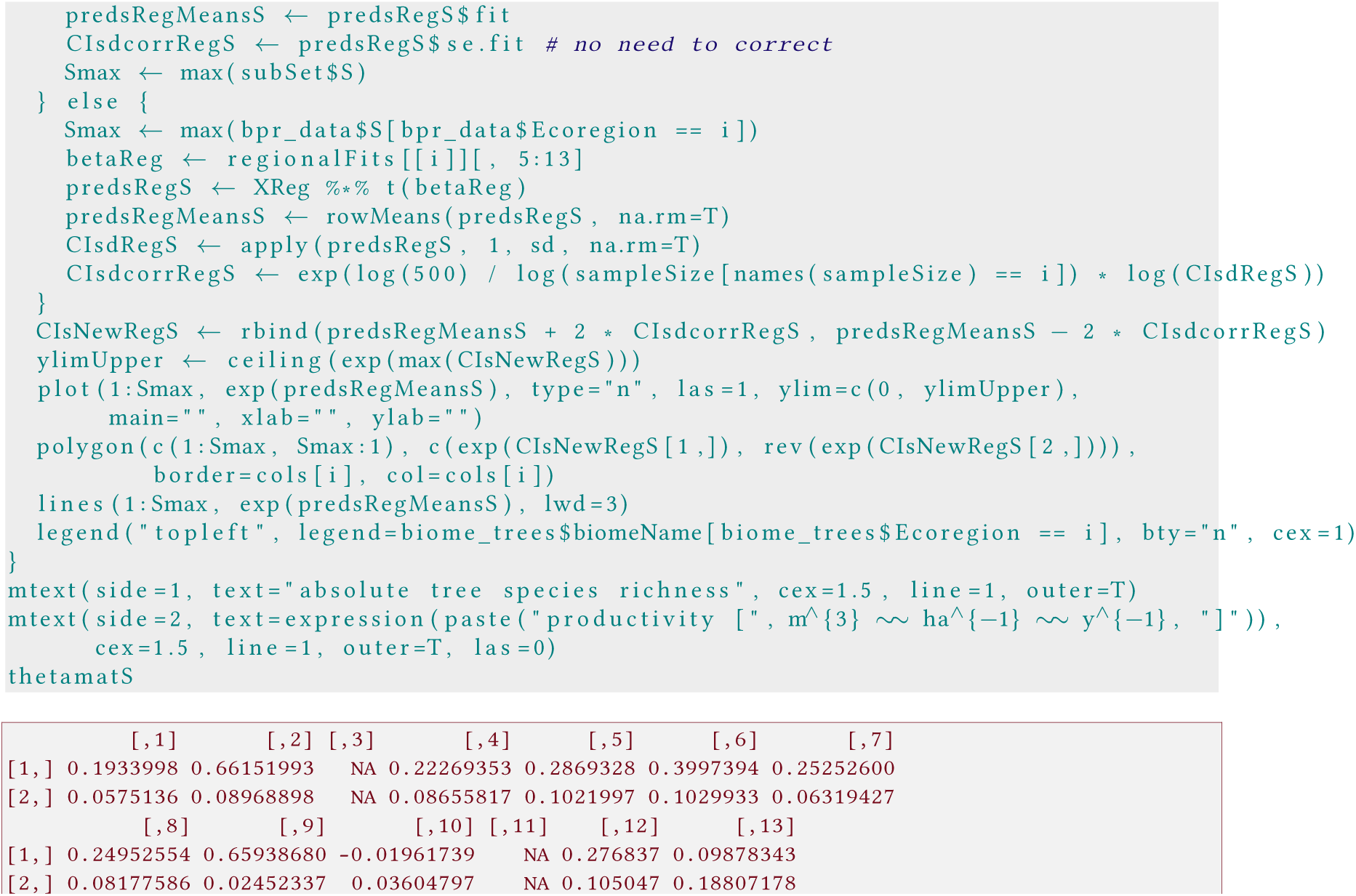

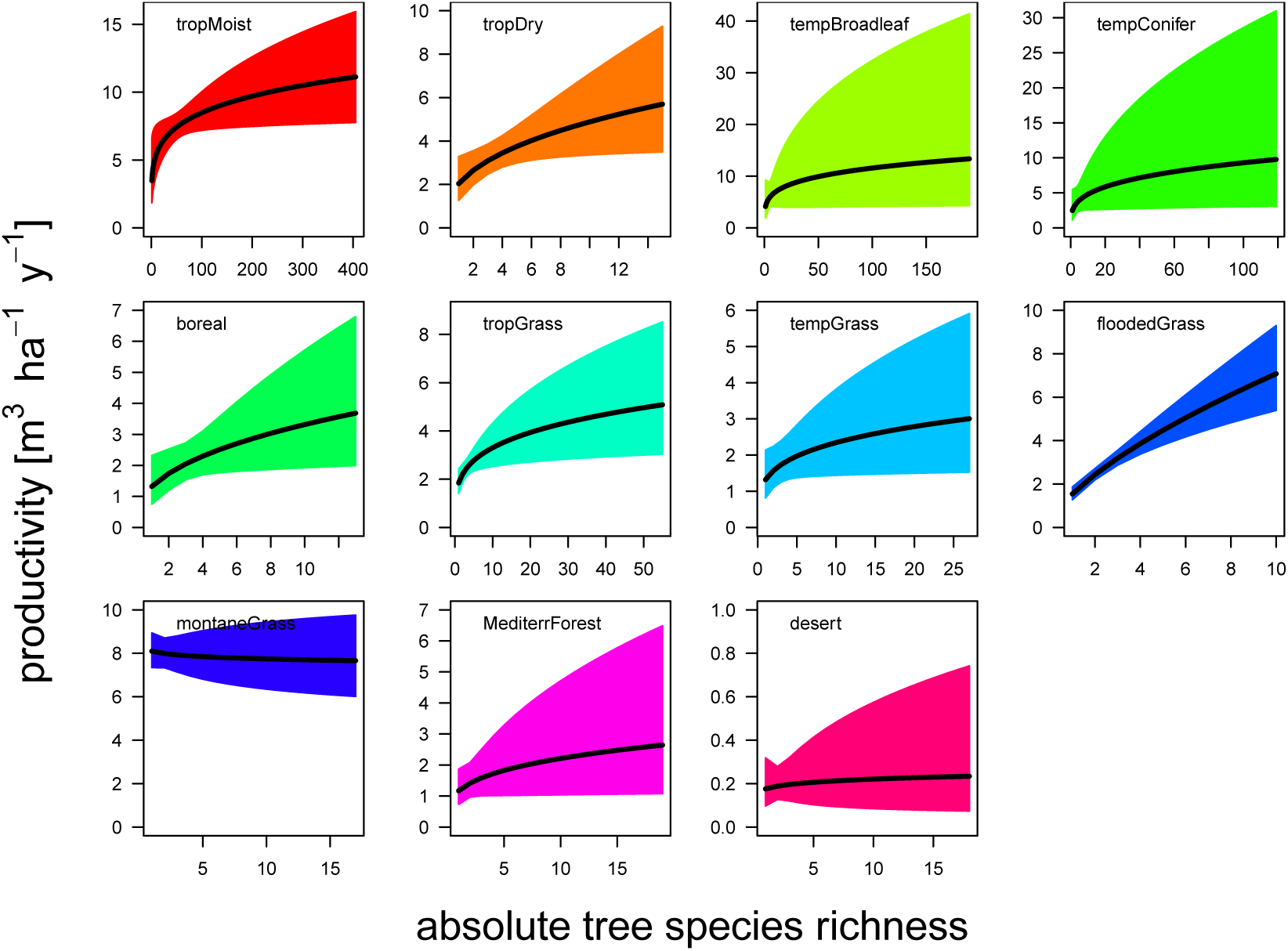

As a summary, we can plot the estimates and their 95%-confidence interval for each ecoregion.

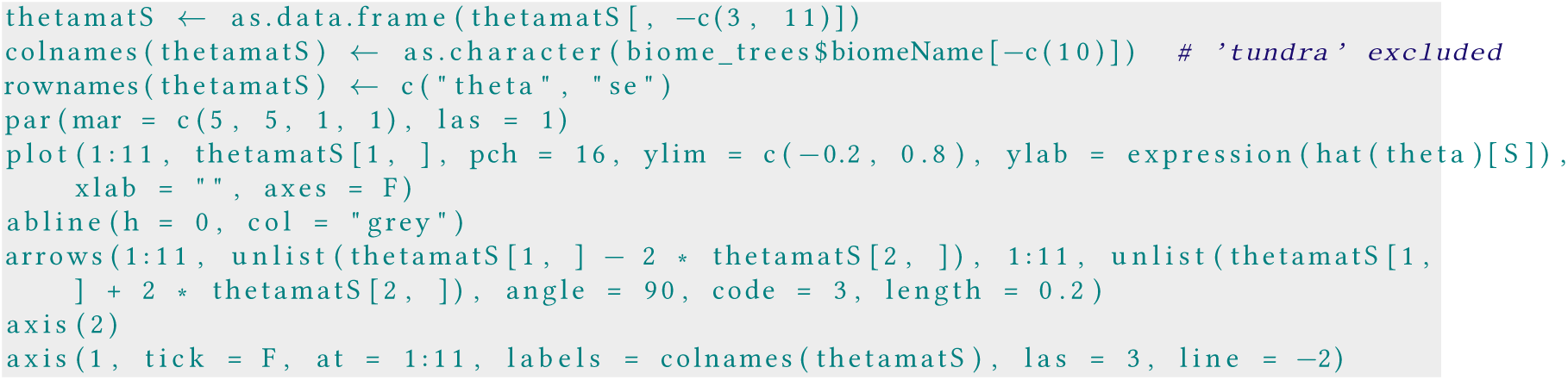

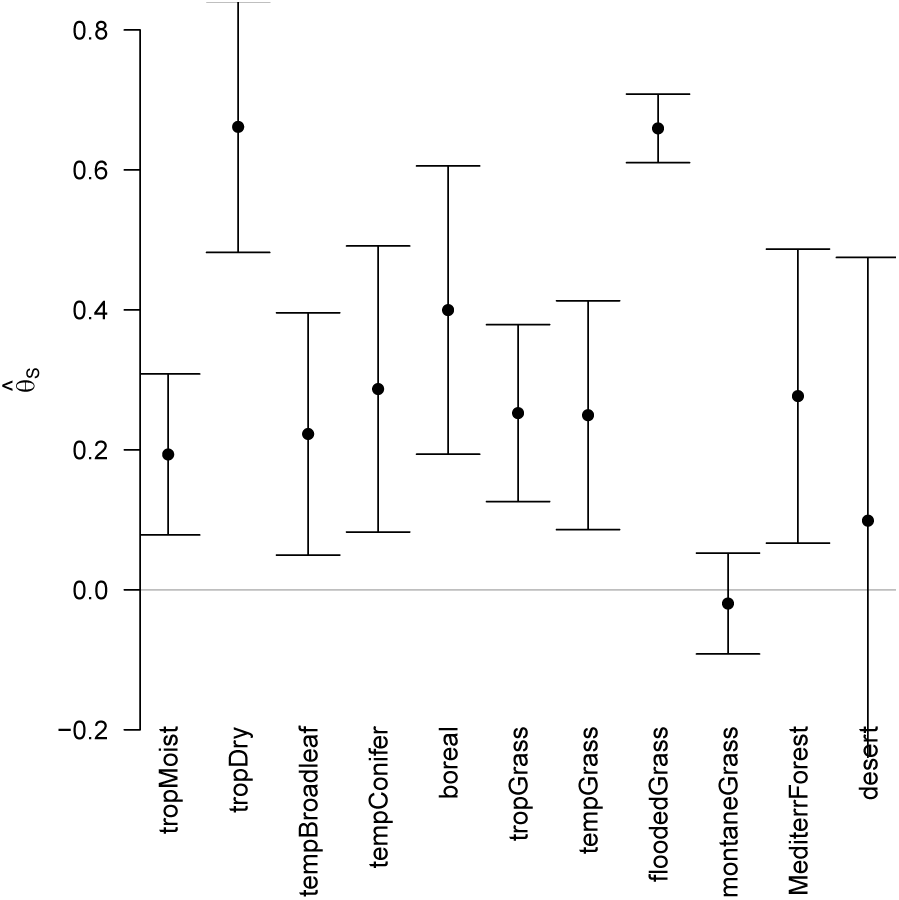

We see that in almost all ecoregions higher tree species richness is correlated with higher productivity. We also see the common pattern of a quick decrease of a distinguishable diversity effect beyond a few species (in tropical moist forests, “few” in this case means a few dozen, while in all other systems “few” means less than 10). ^18^

### F Conclusion

The “global” relationship between “relative tree species richness” and productivity reported by Liang *et al.* (2016) hinges on an analysis with several logical and technical flaws. Removing them yields essentially a flat line and no such global relationship. It is entirely possible that there are mistakes in the above analysis, and that is why we provide this document.^19^

This relationship is *not* globally consistent; rather, ecosystems vary substantially, but most show a positive tree-species richness-productivity relationship. Analysing the data with a relative species richness axis provides no benefit or insight when compared to absolute species richnes analyses.

## Footnotes

1 This R-code of Liang is now available here: https://github.com/jingjingliang2018/GFB1/blob/master/R_script_Github10312008.R

2 We have sent this document on 19 Sept 2018 to the first author and, in response, subsequently (on 31 Oct 2018) he made R-code of (the main part of) their analysis avaiable under https://github.com/jingjingliang2018/GFB1/blob/master/R_script_Github10312008.R. In that code it is clear that plots with productivity larger than 533 m^3^ha^-1^yr^-1^ and with species richness larger than 270 species were removed. The reasoning behind these values is not communicated in that document. Going through their code, we are pleased to notice that we have correctly interpreted their methods section and that our points of content are not mere misunderstandings.

3 Liang *et al*. (2016) invent the name “spatial random forest” for this procedure, which is rather misleading. First, the randomForest algorithm (Breiman, 2001) uses classification and regression trees as building blocks, hence the name “forest” as a large group of “trees”. Second, in randomForest the actual model is altered in each tree, by randomly selecting the variables trialed at each node. Third, while randomForest samples data points, it does so *with replacement*, i.e. as a bootstrap. As a result, the number of data points are the same as the full data set in each tree. What the subsampling approach of Liang *et al*. (2016) and the randomForest have in common is the use of aggregating many models. Since these are not actual bootstraps (because Liang et al. subsample), this is not proper “bagging” either. Thus, the term “spatial random forest” is misleading and suggests a sampling theory which actually does not apply to this approach. Accordingly, we shun this term, and also will not use the term “bootstrapping” unless sampling is carried out *with* replacement.

4 We will return to the computation of relative species richness as a critical flaw later.

5 Sources for each are given in the metadata to the New Zealand data.

6 We will return to the misinterpretation of this correlation as being computed on a sphere later.

7 We shall return to the computation of error bars in a few sections. For now, please note that the classical boostrap’s idea is to use the *entire* data set when bootstrapping; only then will the bootstrapped standard errors be correct. Using the bootstrap on subsamples, in the way done by Liang *et al.* (2016), does not constitute a valid statistical approach for computing standard errors for the full data set, neither in their way, nor in the way presented here (with the multiplication by 100). What this subsampling-bootstrapped standard errors deliver is an estimate of where the line would be with 500 data points, not for the full data set. There are subsampling procedures (see Politis *et al*., 1999), but they require additional corrections on top of the subsampling itself.

8 We do not know how Liang computed these, as he did not reply to our emails on this topic.

9 This is crucial to understand, thus we try to repeat with other words. If we draw 500 plots randomly, with 97% of them coming from temperate forests, then the highest species richness will be set by the very few plots from the tropics, more precisely ecoregion 1: tropical moist broadleaf forest. Thus, we divide the species richness of our temperate plots, say something between 1 and 10, by 80-400. Thus, low Š-values will inevitably be from the temperate or boreal forests, while high Š-values will be from the tropics.

10 Clearly, this is still not an ideal relative species richness, as we can imagine local conditions to vary at relatively small scale. However, we think that this step is already a major improvement in the actual meaning and interpretability of “relative species richness” and will come closer to what we think most of the authors actually understood Š to be.

11 Beyond 5000 the GLS becomes extremely time consuming, running for many hours per model run.

12 The origin of this bias has probably to do with the re-scaling of species richness in each subset. If no high-diversity plot is present, then the left part of the relationship is stretched out to the right hand side without higher P-values, thereby pulling the entire curve down.

13 This is implicit in the corSpher-argument (Pinheiro & Bates, 2000). The different correlation structures (exponential, Gaussian, linear and spherical) describe how the covariance between two locations decreases with their distance. Spherical, for example, is similar to exponential, but with a more linear shape.

14 https://en.wikipedia.org/wiki/Great-circle_distance

15 Taken from https://stackoverflow.com/questions/18857443/specifying-a-correlation-structurefor-a-linear-mixed-model-using-the-ramps-pack.

16 As written above, it is probably in the interaction of the changes that the effect disappears. Changing, for example, the Haversine back to corSpher has hardly any effect in itself. One could now start to remove the changes one at a time to identify the step(s) that caused the original signal to disappear.

17 The marginal effect asks the question: “I’m in a forest somewhere on this planet. I know that this plot has 50% of its potential tree species richness. How productive is this plot?” In other words, the marginal effect is ignorant of the values of the environmental covariates, and as a consequences is far less certain. The conditional effect (“I’m standing in a forest patch in north Sumatra, with that much rainfall, tree density and so forth, and tree species richness is at 50% of its maximum potential: How productive is this site?”) obviously requires more information, but is thereby, by deffnition, not global anymore.

18 These curves are not identical to those from the previous analysis with relative local species richness, as the re-scaling of the x-axis is non-linear: species richness enters the model after log-transformation, thereby also affecting estimation of *θ*.

19 Note that some steps require substantial computing time (all those running a model often).

